# Accessible, Open-source Hardware and Process Designs for 3D Bioprinting and Culturing Channels Lined with iPSC-derived Vascular Endothelial Cells

**DOI:** 10.1101/2021.10.29.466348

**Authors:** Andrew R Gross, Roberta S. Santos, Dhruv Sareen

## Abstract

Indirect bioprinting for cell culture requires the use of several technologies and techniques which currently prevent many researchers not specialized in electrical engineering or materials science from accessing these new tools. In this paper, a printer and all necessary associated hardware was developed and tested for the purpose of seeding human induced Pluripotent Stem Cell (iPSC)-derived endothelial cells (iECs) onto all surfaces of a fibrin-gelatin channel. Immature iECs were seeded onto all channel surfaces and completed differentiation along channel walls. All required tools and methods, including engineering drawing, printable files, code, and hand-tool templates, have been provided with sufficient clarity to enable full, open-source replication of all technique employed.

## 1. Introduction

The rapid growth of the field of 3D bioprinting is driven in part by the development of cutting-edge approaches, but also by method refinements that allow these cutting-edge technologies to become accessible, routine techniques. While specialists push the boundaries of bioprinting technology, the field’s current growth in size relies heavily on adoption of techniques across biology labs without a primary focus in electrical and mechanical engineering. These labs require onramps to bioprinting which provide the greatest benefit to their research at an attainable investment of time and money. This paper describes strategies aimed at a broad audience which integrate and refine emerging methods for indirect 3D bioprinting into a collection of accessible step-by-step protocols to bioprint vascularized constructs using human iPSC-derived vascular endothelial cells (iECs). These protocols allow these iECs to be generated and subsequently live-imaged within a continuously perfused enclosure over an extended time period as a platform for developing assays such as a diffusional barrier assessment. They are composed with special attention paid to common obstacles and the means to overcome them.

The methods described in this paper were developed to assay the barrier function of iECs, but were prepared to make up a complete system capable of being applied to a broad range of experimental needs, such as multicellular co-culture and multi-organ system models. Vascularized 3D constructs provide a much-needed experimental and clinical tool. They’ve been identified as a potential means of assessing drug permeability across the blood brain (Shin *et al.*, 2019; Bhalerao *et al.*, 2020), modelling liver toxicity (Nguyen *et al.*, 2016; Ma *et al.*, 2020), and improving survival of cell grafts (Derakhshanfar *et al.*, 2018; Cidonio *et al.*, 2019; Erdem *et al.*, 2020).

Among bioprinting approaches, one of the most modular and accessible is indirect printing, in which the printed material is a sacrificial scaffold which produces hollow cavities in a bulk material once the scaffold is dissolved. Among the refinements described here are the use of iPSC-derived endothelial cells (iECs) differentiated from a line expressing a constitutive nuclear GFP marker, seeded in a housing designed for live-cell monitoring. Prior studies have seeded indirectly printed channels with HUVECs (Kolesky *et al.*, 2016; Yang *et al.*, 2016; Costa *et al.*, 2017; Fitzsimmons *et al.*, 2018), however iECs offer several advantages. Unlike HUVECs, these cells can be differentiated into specific, relevant endothelial lineages, and they can be prepared from stem cells reprogrammed from patients possessing unique disease genotypes. Most importantly, the use of a GFP-expressing cell line was found to radically improve the process of developing bioprinting processes by allowing experimentalists to monitor the morphology and placement of cells throughout the duration of experiments in a way that was impossible under brightfield or phase contrast microscopy. Additionally, iECs can be cultivated from a diverse source of patient lines to model genetic diversity or genetic disfunction. They can be co-cultured with other cell types derived from the same patient line. They can also eventually be prepared for autologous human engraftment using a patient-specific source. Based on their versatility and ease of culture, iECs would make a fitting replacement in the many instances in which HUVECs (Noor *et al.*, 2019; Skylar-Scott *et al.*, 2019) or dermal fibroblasts (Jang *et al.*, 2017) are used as a default endothelial cell placeholder.

Here, a modular and flexible bioprinting system is described which was used to produce channels in a fibrin-based construct which were subsequently seeded with immature endothelial cells which underwent maturation and staining within channels. Special attention has been paid to elucidating steps which were found under-described within the literature and demonstrating low-cost tools to overcoming these barriers. The complete methodological process can be found in the supplementary methods.

## 2. Materials and Methods

In order to prepare a cell-laden construct or an endothelial cell-lined vascular model a commercial desktop 3D printer (Vertex k8400) was modified to use a compact mechanical extruder and a build plate adapter to hold print surfaces, along with a confined perfusion housing. Sacrificial scaffolds were designed in Autodesk’s Fusion360 CAD software and printed in 30% Pluronic F-127 sacrificial ink before casting a fibrin-gelatin matrix around the scaffold and flushing it out. Endothelial cells differentiated from iPSCs were introduced by filling the channels with a suspension of iECs. These cells were maintained for seven days before being stained for endothelial marker proteins VEGFR2 and CD31. Additionally, modifications to the bioprinted construct housing to accommodate live imaging was determined to be critical to the development process.

Figure 1 outlines the workflow to set up this system. Special focus has been paid to outlining practical steps in bioprinting which have often been insufficiently described in prior literature. While approaches to modifying low-cost commercial desktop 3D printers have been described elsewhere (Banović and Vihar, 2018; Fitzsimmons *et al.*, 2018; Pusch, Hinton and Feinberg, 2018; Kahl *et al.*, 2019), our efforts to assess iPSC-derived endothelial cell (iEC) barrier function in 3D bioprinted constructs identified the most commonly overlooked barriers to 3D bioprinting once the printer itself is constructed.

**Figure 1:**
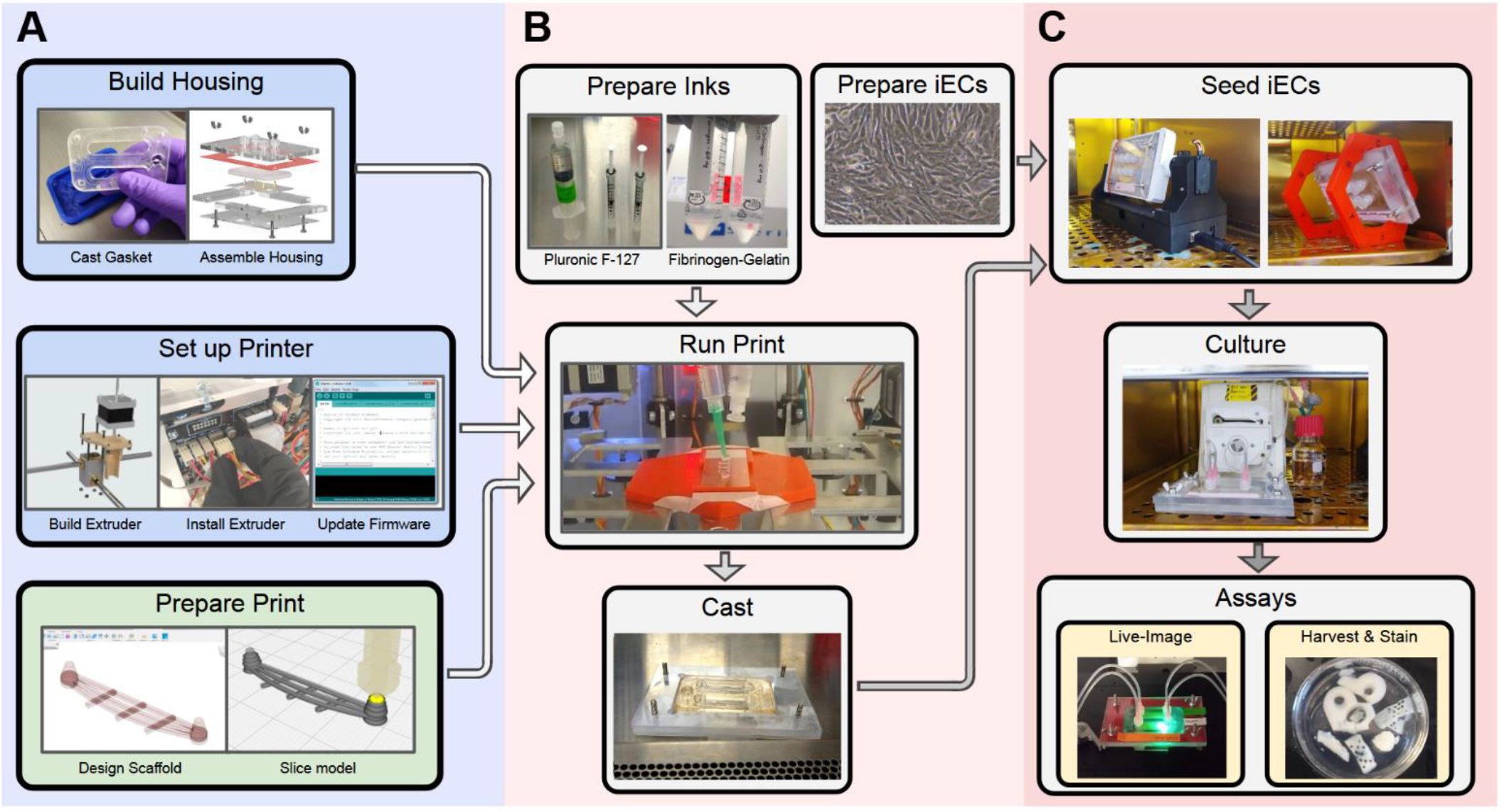
A) Preparation steps include building the housing, installing a compact mechanical syringe pump on an FDM printer, and generating the tool paths from a 3D file in Cura. B) Steps required to produce the construct and cells include ink preparation, initial iEC differentiation, printing, and casting. C) Once the construct and cells are ready, the cells can be seeded, cultured, and eventually analyzed either within the housing or by removing the construct and sectioning it.

The first challenge in bioprinting that was found to be under defined within literature was the need for a confined housing that supported sterile, continuous long-term flow. This support hardware needed to not only maintain cells, but also provide means of observation for the purpose of performing useful experiments in a manner flexible enough to be applicable between experiments. The construction of such a housing from a polycarbonate base and lid sandwiching a PDMS gasket is described in detail in Supplementary methods part 1. Second was the assembly of a suitable bioprinter for printing small, sterile volumes. Supplementary methods part 2 describes the construction and coding of such a printer with emphasis on the specific functionality required to print inks as described in this method. Third, the process of translating an intended geometry into commands a printer can use were found to be almost entirely overlooked in literature. This made the process of designing and printing hollow channels described in published studies unreplicatable. Supplementary methods part 3 describes an accessible means of designing scaffolds to produce hollow channels and converting those designs into g-code machine instructions broadly compatible across printers using free, user-friendly software that requires no custom scripts and no programming experience.

The process of preparing the necessary inks – primarily Pluronic F-127 with FITC-bound dextran and observing the change in fluorescence in the surrounding matrix via fluorescence microscopy. These methods were developed with an emphasis on improving repeatability, so each is described in extensive detail in the supplementary materials.

### 2.1 Constructing the Microscope-compatible Confined Perfusion Housing

Recapitulating the 3D qualities of living animals requires flow of medium, and this this requires confinement. Here, this was achieved by sandwiching a PDMS gasket between two polycarbonate sheets and then iterating the design to meet several crucial design requirements identified during development. The first was that printing had to be performed on a separate surface and then installed into the housing rather than requiring printing to occur within the housing. This was because bioprinting – as with plastic FDM printing – is an imperfect process, and an unsuccessful first attempt should not halt an experiment and require the cleaning and resterilization of major hardware. The use of glass microscope slides as a separate print surface also simplified sterility requirements by reducing the need to sterilize the much larger structural base of the housing.

Next, ports to the housing consisted of polycarbonate Luer-Lok fittings that were solvent-welded into a polycarbonate lid to ensure that flow lines made firm connections that could not accidentally separate if tugged and reduced the risk of leaks and contamination that come with connecting flow lines through fittings held by the elastic pressure of a rubber gasket.

Third, it was found that a microscope-compatible platform was needed in order to allow users to monitor cells, as a bulky enclosure deprived users of the ability to monitor cells as they would in a planar culture and allow for microscope-based assays such as the permeability assay used to assess barrier function of endothelial cells. This was achieved by printing sacrificial scaffolds onto glass slides which were installed into a base with the same footprint as a standard tissue culture plate. The gasket was placed on top of the glass slides then the lid on top of that, and the lid was screwed down with thumb screws to create a fluid-tight environment (Supplementary methods part 1).

### 2.2 Constructing the Compact Cartridge Extruder

Basic bioprinters are increasingly available commercially, however the custom-built route remains an order-of-magnitude cheaper and allows for a level of control, customization, and repairability that makes them an attractive alternative. Several designs have been described previously (Banović and Vihar, 2018; Pusch, Hinton and Feinberg, 2018; Bessler *et al.*, 2019; Kahl *et al.*, 2019) however the design described here was designed to provide a specific, critical functionality missing elsewhere: accommodation of small, sterile volumes. These requirements make any tubing between a syringe and a printhead a non-viable option, which required a syringe-mounted printhead that could be loaded and unloaded without tools (Fig. 3). A mechanical syringe pump was found to be highly preferable to a pneumatic driver, as air pressure was much harder to control with software, required much more specialized hardware, and was prone to ejecting the entire contents of a syringe if pressure was momentarily too high. These considerations led to the design and construction of a snap-fit cartridge made of a 5 mL syringe capable of being loaded into the printer rapidly without tools and without compromising its sterility (Supplementary methods part 2).

### 2.3 Integrating the Compact Cartridge Extruder into the Printer

The new printhead was installed by affixing it to the existing gantry and attaching a linear stepper motor. This motor was plugged into the printer’s mainboard as a direct substitution to the original filament extruder stepper motor. The printer’s firmware was then modified to disable the default setting which prohibits extrusion unless the nozzle is heated to the melting temperature of common plastics and to set the correct extrusion distance per turn of the motor (Supplementary methods part 2).

### 2.4 Designing Constructs and Their Files

A 3D printed construct’s external geometry is defined by the shape of the surrounding gasket, and the geometry of its vascular channels are defined by the printed sacrificial scaffold. The gasket was designed to sit within a surrounding polycarbonate base with cavities where the constructs would fill. The gasket was prepared by printing or casting a mold and then casting the gasket in PDMS in the mold.

The sacrificial scaffold was designed as a series of horizontal tubes the diameter of the printer nozzle strung between two posts. This design was selected to accommodate commercial 3D printer slicing software. The slicer interpreted the sacrificial scaffold as a series of slices with their height equal to the diameter of the channels, which was also the width of the nozzle. If the construct was sliced with no outside walls and 100% infill, the Ultimaker Cura Slicer would produce towers with single channels strong between them. Details are provided in Supplementary methods Part 3.

### 2.5 Preparing Inks

The sacrificial Pluronic F-127 ink was prepared by weighing out 7.5 g of Pluronic F-127 and then alternating between sprinkling the powder into a 50 mL conical and wetting the powder with 18 mL of PBS supplemented with 2% pen-strep anti-biotic antimycotic. The Pluronic F-127 strongly resisted mixing and diffusion, so its preparation required that the powder and PBS be alternated so that all powder was wet and no pockets of PBS or dry powder were present in the tube. The final volume was brought up to 25 mL to produce a 30% Pluronic F-127 solution. The conical tube was then chilled to 4 degrees and placed on a rotator in a cold room or refrigerator overnight.

Fibrin-gelatin bioink was prepared to a final concentration of 10 mg/mL by initially dissolving 60 mg of fibrinogen from bovine plasma (Sigma 341573-1GM) in 4.5 mL of PBS supplemented with 120 uL of penstrep antibiotic antimycotic and 60 uL of Calcium Chloride. The solution was thoroughly mixed and then warmed to 37^°^C for 15 minutes. Then, 1.5 mL of 100 mg/mL of autoclaved gelatin (Sigma G2500-100G) was added, followed by 360 uL of 50mg/mL Transglutaminase (Modernist Pantry 1203-50). The ink was then mixed again and filtered using a Millipore Steriflip filter (Millipore). This ink was allowed to sit at room temperature for at least 30 minutes and no more than 90 minutes, which resulted in a firm, clear ink once crosslinked. Considerations for selecting concentrations of fibrinogen and gelatin are described in detail in Supplementary methods part 4.1.

### 2.6 Culturing iPSCs and iECs

IPSC-derived endothelial cells were differentiated from induced pluripotent stem cells in a three-stage differentiation based on Harding et al. 2017. First, human iPSCs were cultured in mTeSR+ stem cell medium (Stem Cell Technologies) on Matrigel coated plates (Corning) prior to passage using EZ-pass passage tools onto fresh Matrigel-coated 10 cm dishes at a density of approximately 5%. After two or three days, when colonies were approximately 1-2 mm across, differentiation was initiated using the stage 1 medium consisting of STEMDiff APEL base medium (STEMCELL Technologies 05275) supplemented with 6 uM CHIR99021 (Xcess Biosciences m60002). After 48 hours the cells were fed with stage 2 differentiation medium consisting of STEMDiff APEL supplemented with 50 ng/mL of VEGF 165 (R&D Systems, 293-VE-010), 25 ng/mL of BMP4 (R&D Systems, 314-BP-010), and 10 ng/mL of FGF-basic (FGF-2, PeproTech, 100-18B). After another 48 hours, the cells were dissociated for five minutes in of Accutase (STEMCELL Technologies, 07922) and resuspended in MV2 base medium (Harding *et al.*, 2017). The cells were counted and either seeded directly into constructs or another 10 cm dish or frozen. Seeding into constructs on day 4 was found to provide a faster development cycle to allow testing after seven days in culture after only eleven days of culture, as opposed to eighteen days of culture if cells were dissociated and seeded into a dish on day 4 and then dissociated again on day 11.

These cells were differentiated from a female hiPSC line, CS83iCTR-33n1_AAV#46, which had a GFP-expression gene stably inserted into the AAVS1 locus. This gene was inserted downstream of a β-actin promoter with a nuclear localization sequence which ensures that GFP is constitutively expressed in the nucleus of the cell line (Hatada *et al.*, 2015).

### 2.7 Printing, Casting, and Evacuating Sacrificial Ink

The g-code toolpath file was loaded into Repetier host software according to the printing instructions in supplementary methods part 4. Scaffolds were printed in 30% Pluronic F-127 on sterile glass slides at room temperature using the mechanical extruder, then placed within the perfusion housing base. Food-grade silicone oil (McMaster 3025K15) was gently applied to the face of the PDMS gasket that would contact the slides and the PDMS gasket was set in place. In cell viability tests, dissociated HEK293T cells were spun down and resuspended in complete fibrinogen-gelatin ink at 6 E6 cells/mL.

Thrombin crosslinker was added at a concentration of 50 U/mL to the conical of room-temperature fibrinogen bioink, applying 60 uL of crosslinking solution per mL of bioink. This solution was mixed for 10 seconds via pipetting while taking care to avoid creating bubbles, then was dispensed into the housing. The ink has a typical working time of around 30-40 seconds after the introduction of the thrombin crosslinking solution at room temperature.

### 2.8 Seeding iECs into Channels

IPSC-derived endothelial cells seed readily onto fibrinogen-gelatin ink in a dish and will propagate to cover surfaces. Seeding cells onto the walls and ceiling of channels, however, requires reorienting the bioprinted construct so that the cells settle onto the desired surface. To do so, 1E6 day 4 iECs were resuspended in a volume equal to or greater than the void space of the channel (200 μL) and introduced via one port. The cells in suspension settled in minutes, and within one hour began to attach. Seeding was achieved using two approaches. Initially, the construct was inserted into a hexagonal bracket (Fig. 4D) that allowed for its placement in six different orientations. Later, this seeding process was automated using a custom designed rotator (Fig. 4E) which would cycle through positions. When rotating manually, the housing was rotated 120 degrees after one hour and then the cells were allowed to attach to a second surface and left overnight. The automated rotator rotated through eight positions, changing hourly, which produced significantly more coverage during each seeding.

By using GFP-labeled iPSCs, cell placement and shape can be viewed more readily than when relying on phase contrast microscopy. The next day cells were fed and additional cells were seeded as needed to sufficiently introduce cells onto all walls, after which cells would proliferate to close small gaps and cover surfaces.

### 2.9 Staining cells for markers

When staining cells lining a channel, fixation was performed inside the housing. Seven days after the last seeding the channel was gently aspirated and filled with 4% paraformaldehyde for 15 minutes, then gently aspirated and washed with phosphate-buffered-saline (PBS) three times. It was then blocked for 2 hrs with PBS containing 10% donkey serum (Millipore) and 0.15% Triton X-100 (Bio-Rad). Primary antibodies were diluted in blocking buffer and placed on a rocker for 1 hour at 25 rpm, then left in the channel overnight at 4oC. Primary antibodies were VEGFR2 (1:50, Cell Signalling #2479) and CD31 (1:50, Cell Signalling #3528). The channels were then aspirated and washed three times with PBS with 0.1% TWEEN-20 (Thermofisher) and then stained with secondary antibodies AlexaFluor 568 anti-mouse (Fisher Scientific A10037) or AlexaFluor 568 anti-rabbit (Fisher Scientific A10042) and incubated overnight at 4^°^C. The channel was then washed and imaged on a Nikon confocal microscope in the Nikon Elements Software.

### 2.10 Performing FITC Diffusion Assay

In order to assess the barrier function of cells lining a channel, a 0.5 mg/mL solution of 4 kDa FITC-bound dextran (Sigma FD4-100MG) was flowed through a channel at 30 mL/hr for thirty minutes during a confocal timeseries captured every 20 seconds for 30 minutes as described in Supplementary methods part 7.

## 3. Results

### 3.1 Compact Cartridge Extruder Allows for Reliable Printing of Sacrificial Scaffolds

A compact linear actuator assembled from simple, low-cost materials can accommodate a syringe cartridge which can be loaded instantly without tools (Fig. 1A). In the process of developing a bioprinter capable of performing routine, daily-use printing, several design limitations were identified among existing tools. Among these were a requirement that the ink be housed in a vessel which could be prepared within a hood in small volumes and loaded into the printer elsewhere quickly without compromising sterility and without requiring tools (Fig. 2A/B). This was achieved by using a simple, compact linear actuator consisting of a stepper motor with a threaded rod and then designing a snap-fit coupling. A 3D printed socket end of the coupling receives a 3D printed plug end that replaces the shaft of a 5 mL syringe. This design was capable of precise, sterile deposition of sacrificial Pluronic F-127 ink without thermal control (Fig. 1B, Fig 2E)

**Figure 2:**
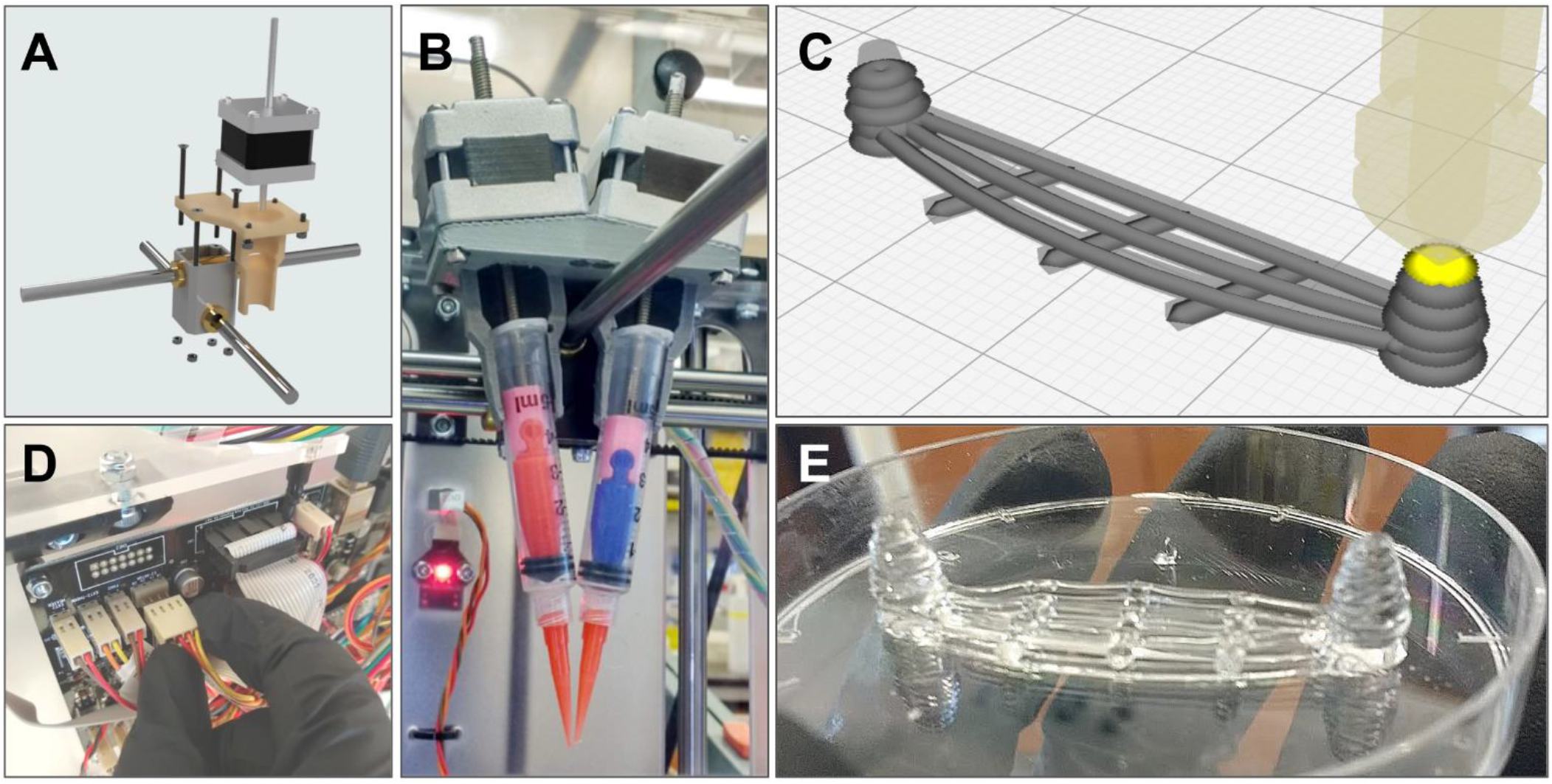
Steps to modify an FDM desktop 3D printer and design a sacrfificial scaffold. A) An exploded view of a mechanical syringe pump printhead. B) The finished dual-extrusion printhead with syringe print cartridges. C) Toolpath visualization in Ultimaker Cura. D) Physically Integrating a linear stepper motor into the printer’s control board. E) A multi-level scaffold with suspended channels.

### 3.2 G-code Extrusion Toolpaths Can be Generated Using User-friendly Open Source Software

Machine g-code extrusion tool path commands were able to be generated within Ultimaker’s Cura open source slicing software by setting several parameters in the user settings and designing scaffolds with printability in mind. Designing vascular networks and translating those designs into printable g-code extrusion toolpath commands has been a challenge largely overlooked in existing literature. Most published examples use either proprietary software packaged with expensive commercial bioprinters (Noor *et al.*, 2019) or custom code written in Matlab, C++, or Python, which are inaccessible to most users (Homan *et al.*, 2016; Kang *et al.*, 2016; Kolesky *et al.*, 2016; Jang *et al.*, 2017; Skylar-Scott *et al.*, 2019). Here, we designed vascular networks using Autodesk’s popular Fusion360 CAD software, which is available as a free package for non-commercial use (Fig. 1A). We then converted our model into a g-code toolpath in the popular opensource Cura slicer by setting the line count to 0 and the infill to 100 (Fig. 2C). Although this slicer is not designed to produce a series of narrow individual strings, toolpaths can be quickly generated by inputting the correct user settings. The placement of features (such as the distance between towers) is set by dimensions within the CAD model, while the width of channels can be adjusted by modifying the flow ratio in the slicer. Using the same model, one can extrude either a 0.9 mm diameter channel or a 1.5 mm channel using the same 21-gauge (0.5 mm ID) dispensing needle just by adjusting the Flow parameter in Cura that modifies the volume of material extruded relative to the distance. Because extrusion is performed by a stepper motor that replaces the original FDM motor through a simple substitution on the motherboard (Fig. 2D), the commands execute correctly and produce high quality prints (Fig. 2E) without the need to convert these commands to be executed by a pneumatic system.

### 3.3 GFP-expressing cells in a microscope-compatible housing provided critical insights

The ability to monitor the position and morphology of iECs within the construct at all times throughout the experiment was critical for process development. Initial attempts to seed cells frequently resulted in unseeded or partially seeded channels that provided insufficient data for further process development and took two weeks to yield results (Fig. 4A). The use of a microscope-compatible housing (Fig. 3A/B) allowed for observations during the experiment, which proved critical, and the incorporation of the nuclear-GFP expressing cell line CS83iCTR-33n1_AAV#46 provided a break-through by making it possible to see cell height by panning through the construct with an ECHO Revolve fluorescent microscope. The GFP expression also communicated the cells density, shape, and viability far beyond what was possible with unlabeled cells (Fig. 4 F/G). The use of GFP-expressing cells precluded the use of available Live-Dead stains, however this trade-off appeared worthwhile as it allowed for an effective confirmation of cell health throughout the course of the experiment in exchange of a more precise assessment at a final timepoint. The presence of GFP was determined to be a reliable indicator of cell viability, as GFP expression rapidly decreased within 24 hours of widespread cell death in tests.

**Figure 3:**
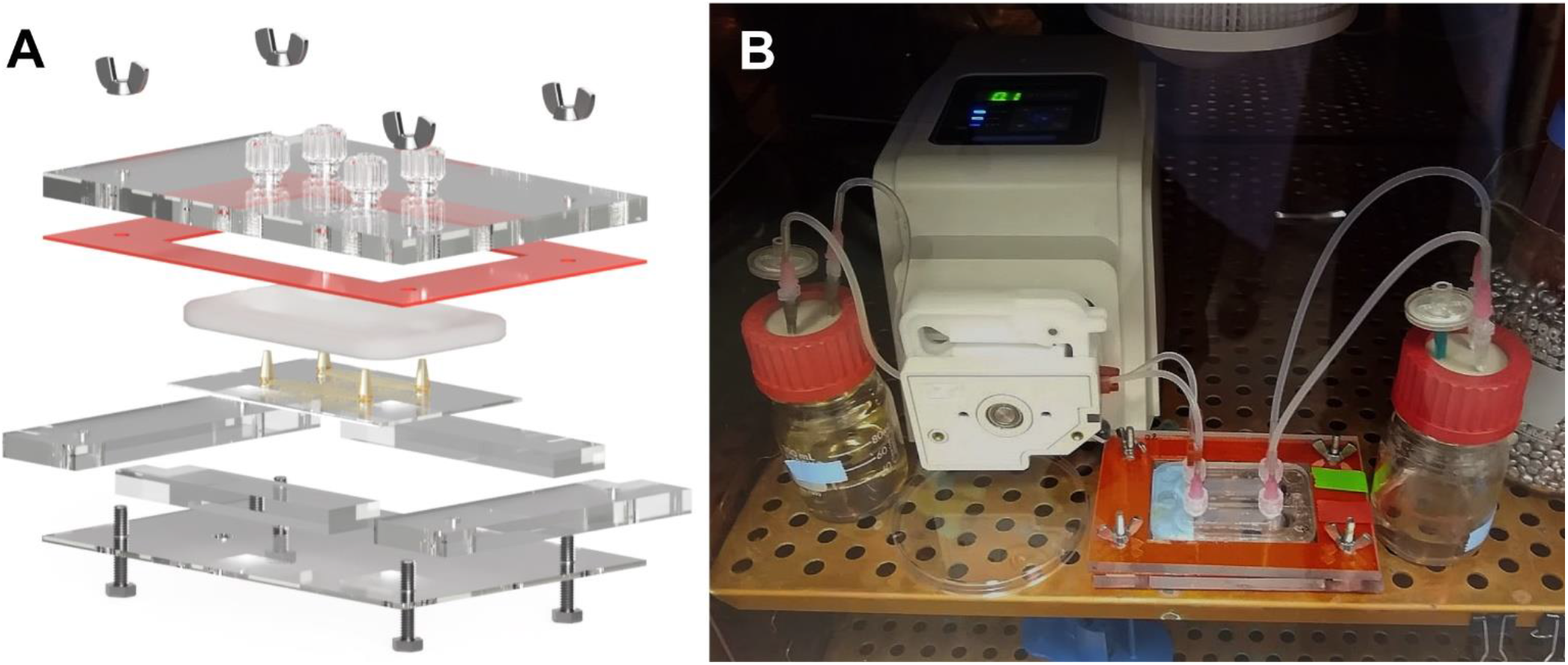
The perfusion system consists of the housing and the accessory hardware, such as the pump and resevoirs. A) An exploded view of the microscope-compatible housing. B) A non-cycling perfusion system moves media from an inlet reservoir to an outlet through two channels in parallel.

**Figure 4:**
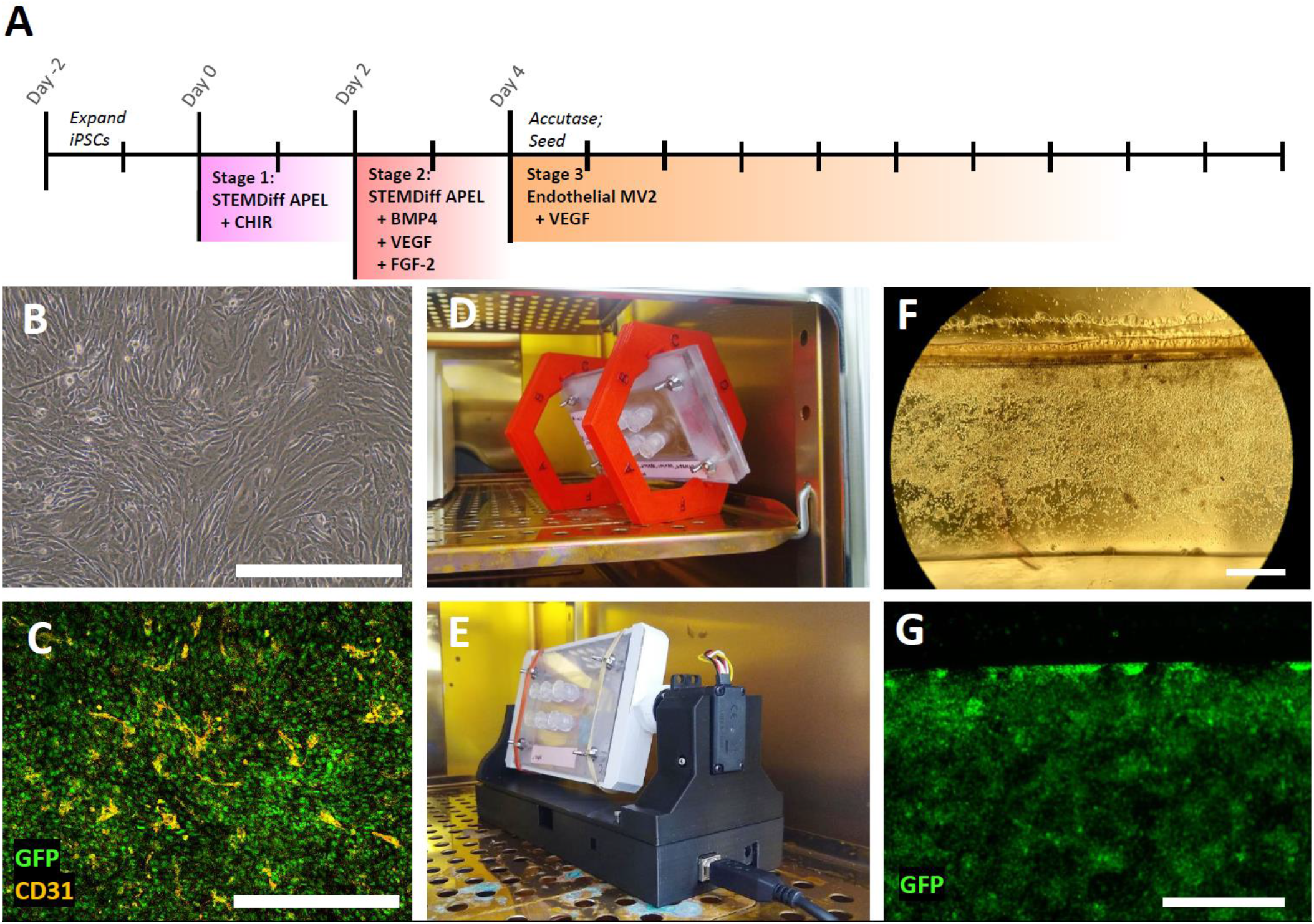
Seeding iECs into a channel. All scale bars are 500 μm. A) Schematic of iEC differentiation. Cells were fixed and stained seven days after the last seeding. B) iECs in a dish on day 11. C) iECs in a dish on fibrin-gelatin basement matrix stained with CD31. D) Seeding using a manual bracket. E) Seeding using a custom rotator. F) Brightfield of iECs the day after seeding. G) 488 nm-channel image of GFP-expressing iECs the day after seeding.

### 3.4 iECs anchored securely to 3D fibrin matrix channel walls

Endothelial cells differentiated from iPSCs into iECs (Fig. 4B) took up residence along the interior of hollow channels and propagated to cover available surfaces when the construct was rotated during seeding and seeded repeatedly over three subsequent days to allow cells to settle on all sides (Fig. 1 C, Fig. 5). These iECs were generated using a 3-stage differentiation over 10 days based on a differentiation described by Harding *et al.* 2017 (Fig. 4A). They were then dissociated with Accutase and introduced into the construct in suspension. If seeded without rotation, the cells settled and attached overnight. By seeding successive days at different orientations, cells were induced to attach and proliferate on all faces of the channels.

**Figure 5:**
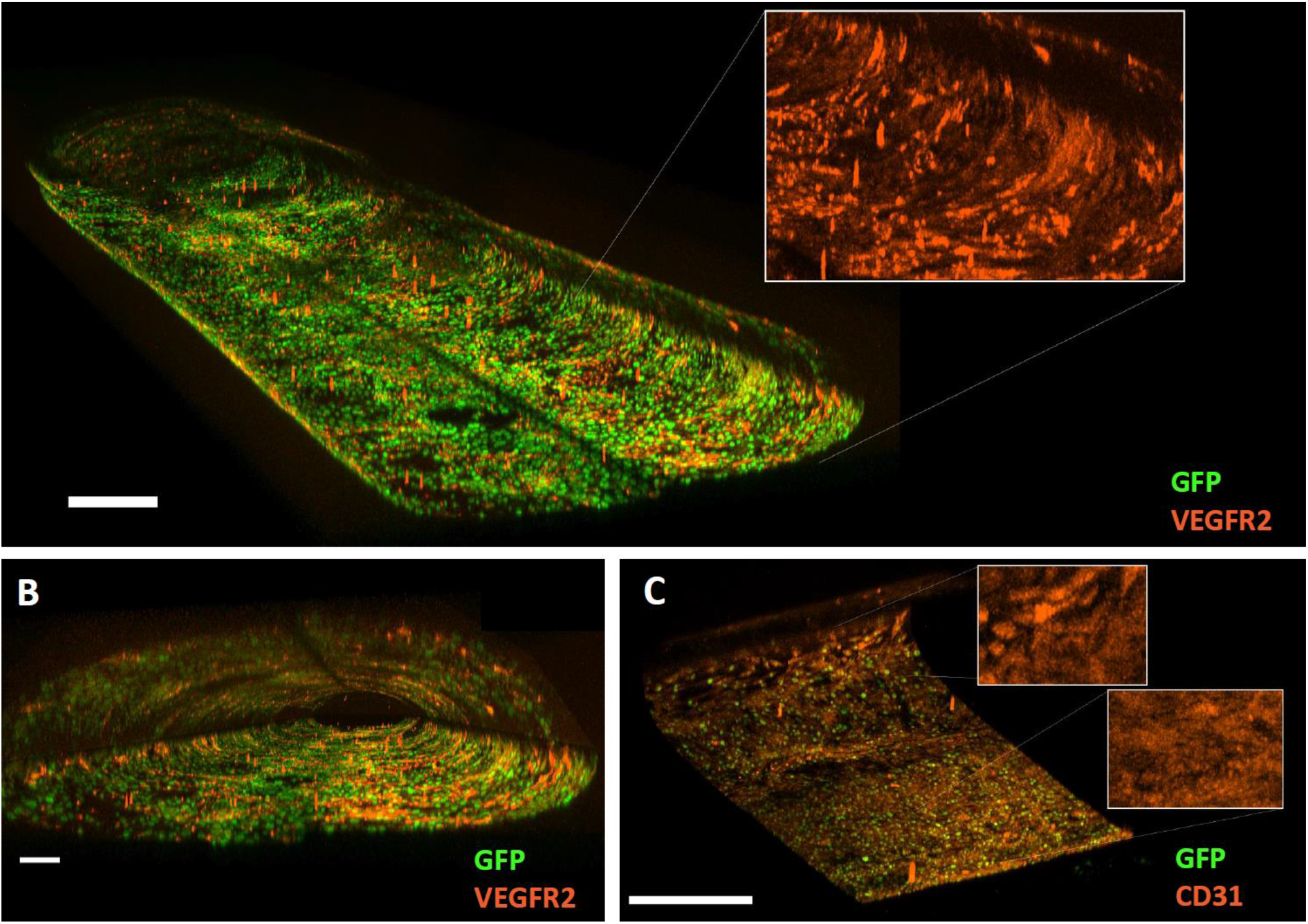
IPSC-derived endothelial cells in bioprinted constructs. A) Nuclear GFP-expressing iECs stained for VEGF receptor 2 after 7 days in culture with enlarged 568 nm channel inset. B) The same construct from stained for VEGFR2 viewed axially. C) Nuclear GFP-expressing iECs stained for CD31 in a channel 7 days after seeding.

Although attachment efficiency varied between batches of iECs produced, the fibrin matrix proved to be highly biocompatible with the endothelial cells, and cells seeded onto fibrin in a dish expressed endothelial cell marker CD31 after 7 days after seeding (Fig. 4C).

Cells demonstrated an ability to propagate to fill gaps over short distances. This required initial seeding to be homogenous, however. A dense mat of cells along the bottom of a channel did not travel to cover the ceiling of a channel within the duration of the experiment. For this reason, cells were allowed 60 minutes to settle on one face of the channel, then the housing was rotated.

Although the manual bracket allowed for successful seeding, an automated rotator was found to be far more efficient. When using the manual bracket, the housing was rotated 120 degrees and allowed 60 minutes to settle in the new orientation, after which time the housing was left overnight. The next day the construct was observed and iECs were suspended and seeded again to cover the uncovered faces and the process was repeated if needed. The seeding process was similar when using the automated rotator, however it was much more effective at covering faces because it rotated through eight positions for 60 minutes each overnight. This allowed for greater coverage in a single seeding, and better distribution.

### 3.5 iPSCs seeded onto channel walls and cultured for one week expressed VEGFR2 and CD31

IPSC-derived endothelial cells seeded at differentiation day 4 and cultured within a channel until day 11 expressed endothelial markers. Cells within the channels exhibited expression of vascular endothelial growth factor receptor 2 (VEGFR2) (Fig. 5 A/B) and platelet endothelial cells adhesion molecule CD31 (Fig. 5C). The widespread expression of these critical endothelial markers indicated their fulfilment of an endothelial cell identity within these channels, which occurred between their seeding on day 4 and the end of the experiment one week after the construct’s last seeding.

## 4. Conclusions

Bioprinting offers a broad range of new possibilities in 3D culture. To researchers seeking an entry point – especially those with a background in cell culture but not engineering or materials science – the biggest challenge is identifying an existing use case they can adapt to fit their research. Such use cases must be instructionally clear enough to allow a broad audience of resourceful non-engineers to replicate the hardware and techniques required. This paper is meant to provide clear tutorials for such labs.

This study identified a series of process improvements that allowed 3D bioprinting to be employed in a variety of experiments. First among these was a reliable, compact mechanical extruder coupled with flexible software. These features allowed routine use with an attainable level of training by most lab members. Secondly, a housing which allowed for observation of cells during the course of the experiment was critical. The use of cell line CS83iCTR-33n1_AAV#46, which constitutively expressed GFP, augmented the ability to observe cells substantially. The presence of a fluorescent marker allowed for a significantly higher degree of observation of placement and morphology than was possible with brightfield or phase contrast microscopy. These two developments were the most significant lessons provided by this project to anyone seeking to begin process development in the bioprinting space.

The use of iECs confirmed their suitability for seeding on the interior of indirectly printed channels and offers a guidepost for further development. Within bioprinting, iPSC-derived cell types are a well-recognized tool. Their appeal derives from their ability to provide an abundant source of diverse cell types otherwise unavailable, as well as the ability to generate cocultures of multiple cell types which share the same genome. These cells can be derived from patients of relevant genetic disorders and potentially be utilized in autologous transplants. While the use of iECs is expanding in bioprinting, these have been primarily limited to cell-laden inks (Zhang *et al.*, 2014; Bulanova *et al.*, 2017; Gao *et al.*, 2017), while most attempts to seed endothelial cells into indirect channels relied on HUVECs (Yang *et al.*, 2016; Costa *et al.*, 2017). Here, we describe a toolset that will hopefully encourage the wider use of iECs in indirect bioprinting experiments.

Fibrin as a substrate demonstrated great use in its biocompatibility and its amenability to casting. It’s 30 – 60 second working time made it suitable in use cases incompatible with materials requiring UV crosslinking or which experienced unsuitably fast crosslinking. Pluronic 127, however, must come with a warning: it’s high printability and appealing melting properties come at the cost of major challenges to sufficient removal required for seeding cells.

Finally, it must be emphasized that even more so than in open software and open hardware development, effectively disseminating methods of 3D cell culture and handling require a proactive determination of communicate the nuanced challenges of hands-on wet work. A willingness to share alone is often not enough to transfer the arcane but crucial tricks to successfully culture sensitive living cells in novel formats. This paper is meant to contribute to what appears to be a rapidly growing movement: to take the principles of open software design that have already moved from the computer science field into an indispensable role within bioinformatics and adopt them further into the development of methods and hardware for wet lab work. This model of information sharing may be crucial to the popularization of bioprinting, as experiences with the alternative approach – closed platforms – have shown a concerning pattern: often a polished and feature-rich hardware solution is rendered incompatible with a broader workflow due to an inability to customize its hardware or software. Additionally, the field of bioprinting requires greater attention to the accessory hardware necessary for cell seeding and confined, sterile flow. Fortunately, the adoption of open source hardware is well underway, especially within the field of microfluidics (Kong *et al.*, 2017; Walsh *et al.*, 2017; Stephenson *et al.*, 2018; Gao *et al.*, 2020; Felton, Hughes and Diaz-Gaxiola, 2021).

The rapid proliferation of innovative open-source hardware designs – especially during the COVID19 pandemic -- can be seen in the growth of repositories (metafluidics.org, 3dprint.nih.gov, github.com/MakerTobey/OpenMicrofluidics), journals (The Journal of Open Hardware, HardwareX), consortia (Gathering for Open Science Hardware, Open Source Hardware Association), businesses (CHAI, Opentrons, Lulzbot) and many, many projects such as this one.

## Supporting information

Full Supplementary Methods Sareen Lab Bioprinting Protocols

## Author Contributions

ARG: Design of experiment, hardware, and methods; construction and performance of experiment; data analysis; writing

RSS: Design of experiment

DS: Design of experiment

## Acknowledgements

The authors wish to thank Allison Tran, who assisted with tests on enzymatic digestion of bioprinted constructs as a summer internship project as well as Dr. Melodie Metzinger and Trevor Nelson for their insights during discussion on the mechanical behaviors of protein-based hydrogels.

## Statement of Ethics

The stem cell lines used this research were conducted under the supervision of Cedars-Sinai IRB study Pro00036896.

## Data Availability

All necessary 3D model files and code is available at https://github.com/andrewrgross/3D-Bioprinter-parts-and-accessories.

