## Supplementary material for "Accessible, Open-source Hardware and Process Designs for 3D Bioprinting and Culturing Channels Lined with iPSC-derived Vascular Endothelial Cells": Full Supplementary Methods Sareen Lab Bioprinting Protocols

#### Contents

**I. Designing Confined Perfusion Housing for 3D Culture 4**

**1. Cut Polycarbonate for the Base Layer and the Lid Layer 5**

**2. Cut and assemble the support layer outside of the gasket 5**

2.1. Cutting out the support layer 5

2.2. Assembling the complete base layer 6

**3. Make the Gasket 9**

3.1 Modeling the Gasket 9

3.2 Preparing a Mold 12

3.3 Casting PDMS in the mold 12

3.4 Demolding the gasket 13

4. Install the Luer-Lok Ports in the housing lid 13

4.1 Drill holes 13

4.2 Tap the Holes 14

4.3 Solvent Weld the Ports 14

**5. Assemble the Flow System 14**

**II. Building and Integrating Compact Mechanical Extruder 17**

**1. Designing the Compact Cartridge Extruder 17**

1.1 Syringe dimensions 17

**2. Modeling the Coupler 19**

**3. Integrating the Compact Cartridge Extruder into a Printer 22**

3.1 Mechanically integrating the extruder 22

3.2 Updating the printer's firmware 22

3.3 Electrically wiring in the new extruder 24

**4. Integrate a Stage Bracket 25**

4.1 Designing a Stage Bracket 25

**5. Setting up a Printer Control 26**

**III. Designing Scaffolds and Preparing Printable G-Code Files 29**

**1. Designing and casting Silicone Gaskets 30**

1.1 Modeling the gasket 30

1.2 Modeling the printable gasket mold 31

**2. Designing the printable scaffold 32**

2.2 Generating printable files 33

2.3  Optimizing the model for printing 34
2.4 Refining the print conditions 36

**IV. Preparing Bioinks, Printing, and Casting Constructs 37**
**0. Preparing to print 37**
**1. Preparing bioinks 38**
1.1. Preparing 30% Pluronic F-127 38
1.2. Preparing Fibrinogen Bioink components in advance 40
1.3. Preparing complete fibrinogen-gelatin bioink on the day of use 41
**2. Printing a scaffold 43**
**3. Casting fibrinogen-gelatin ink around a sacrificial scaffold 44**
3.1. Prepare the housing to receive the FG ink 44
3.2. Add thrombin to initiate crosslinking and dispense quickly 45
**4. Flushing out the sacrificial scaffold 46**

**V. Generating iECs and Seeding Constructs 47**
**1. Generating iPSC-derived Endothelial Cells 47**
1.1 Culturing iPSCs 48
1.2 Differentiating Cells 48
**2. Seeding iECs into channels 48**
2.1 Determining the volume and surface area of a construct's void spaces 49
2.2 Seeding cells into a construct 49
2.3. Assessing seeded cells and feeding 50
Bioprint Construct Assessments 52

**VI. Culturing, Harvesting, and Imaging Constructs 54**
**1. Maintaining Bioprinted Constructs in Culture 54**
1.1: Flow system setup 55
1.2. Feeding manually 56
**2. Live-Imaging Constructs 57**
**3. Harvesting 57**
3.1. Sectioning for staining 57
3.2. Digesting fibrin for RNA extraction 58
**4. Staining and Imaging Fixed Constructs 58**
4.1: Immunohistological staining solutions 58
4.2: Immunohistological staining procedure 59
4.3: Viability staining solutions 59
4.4: Viability staining procedure 60
**5.  Confocal fluorescence imaging 60**

**VII. Assessing Diffusional Barriers 61**

**1. Prepare the flow system and reagents** **62**
1.1: Prepare the FITC flow system 62
1.2: Prepare the FITC-Dextran solution 63
**2. Calibrate the microscope settings** **63**
2.1: Frame the point of interest in their X-Y positions 64
2.2: Locate the ideal Z-position for capture 64
**3. Capture diffusion** **64**
3.1: Preparing the channel 64
3.2: Flush channel and begin capture. 65
**4. Post-process image stack** **66**
**5. Analyze and quantify the rate of diffusion** **66**
**6. Compare the speed of diffusion between time serieses** **67**

### I. Designing Confined Perfusion Housing for 3D Culture

**1. Cut Polycarbonate for the Base Layer and the Lid Layer 5**
**2. Cut and assemble the support layer outside of the gasket 5**
2.1. Cutting out the support layer 5
2.2. Assembling the complete base layer 6
**3. Make the Gasket 9**
3.1 Modeling the Gasket 9
3.2 Preparing a Mold 12
3.3 Casting PDMS in the mold 12
3.4 Demolding the gasket 13
4. Install the Luer-Lok Ports in the housing lid 13
4.1 Drill holes 13
4.2 Tap the Holes 14
4.3 Solvent Weld the Ports 14
**5. Assemble the Flow System 14**

A perfused construct must be confined so that flow is only able to in, through, and out of intended positions. The following guide outlines the design and construction of two confined perfusion housing systems which can be used as is or modified and iterated upon. One is optimized for simplicity, while the other is designed to allow for live imaging on a microscope.

Both housings are made by sandwiching a PDMS silicone gasket between two pieces of polycarbonate, with printing and casting taking place on a separate surface..

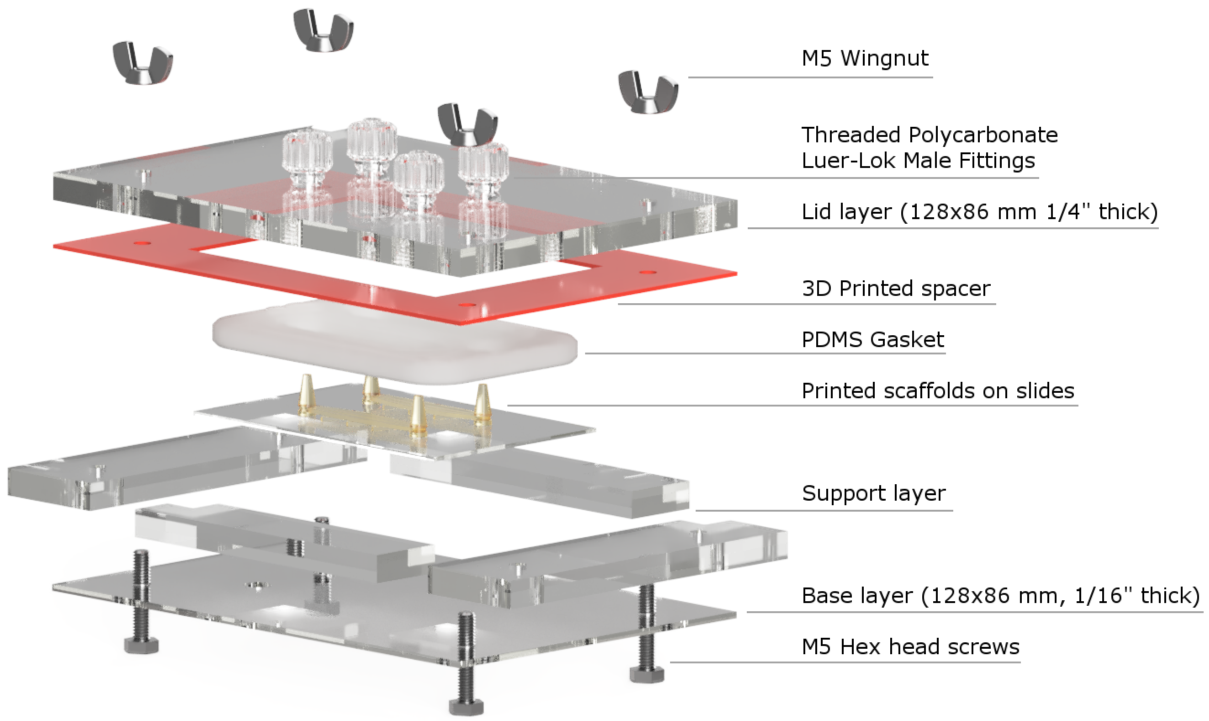

*Supp. Fig. 1.1: An exploded view of the microscope-compatible housing*

### 1. Cut Polycarbonate for the Base Layer and the Lid Layer

The basic model can be cut by dividing a 6” piece of 1/4 “ polycarbonate into quarters.

The microscope-compatible housing is assembled from a 1/16” thick polycarbonate base layer, solvent welded to 1/4" thick support layer, each 128 x 86 mm.

### 2. Cut and assemble the support layer outside of the gasket

#### 2.1. Cutting out the support layer

The PDMS gasket is too soft to hold the acrylic plates apart, so a spacer layer must be made to sit outside the gasket. When the screws are tightened, the lid and base layers will compress the spacer and the gasket.

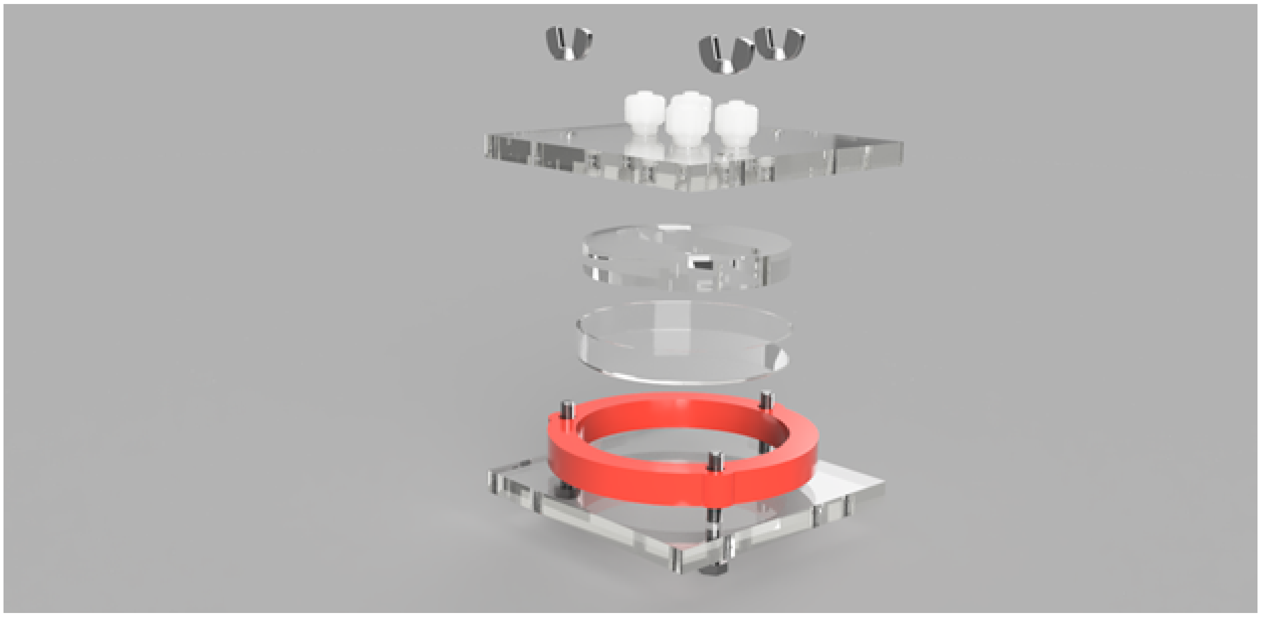

*Supp. Fig. 1.2: An exploded render of the basic microfluidic housing*

For the basic housing, the support layer is a 3D printed ring that encircles the 60 mm dish containing the printed scaffold. This is extremely easy to model and print.

For the microscope-compatible housing, the support layer must be bonded to the thin bottom layer to provide adequate rigidity. This can be prepared either by cutting out the support layer as a single piece (which is the recommended approach), or by cutting two or more separate pieces, which may be easier given the tools available.

To prepare a single piece, cut out it's outside dimensions, then drill the corners of the inner region with a drill press or by printing the drilling jig included in the printable supplementary files, then cut the inside dimensions with a reciprocating saw, a rotary tool, or jigsaw. The corner holes are 7/32" in the printable template to maintain consistency with lid holes that must be tapped, but these corner holes can be enlarged as necessary to accommodate the size of a saw blade.

#### 2.2. Assembling the complete base layer

Once cut, the thin base layer and the support layer (or pieces of the support layer) are then solvent welded using SciGrip 4 acrylic cement. Solvent welding is achieved by clamping polycarbonate components tightly together and depositing solvent along the seam with a needle-tipped applicator, as described in the manufacturer's instructions. The solvent wicks into the space between, dissolves both sides, and then evaporates leaving the two components molecularly bonded.

**Solvent welding large surface areas**

The solvent will wick well into surfaces in close contact, but when joining such a large surface area it is typical that the solvent will not penetrate into all areas. Wicking can be maximized by clamping the surfaces tightly, sandwiched between two other hard surfaces like blocks of wood or more polycarbonate and applying the cement meticulously along the outside perimeter. A bit of very gentle flexing may be applied to distribute the cement.

The solvent cement can also be applied on the interior, however the solvent can distort the junction where the support layer meets the base layer, and this can affect the fit of glass slides inside the support layer. One way to avoid this is to simply apply the cement into the drilled corners. This, in conjunction with application along the outer perimeter typically achieves excellent wicking penetration. Afterwards, allow the cement to evaporate fully before handling, either by leaving it overnight or placing it in a drying oven.

Once the bonding is complete, be aware that water which enters into gaps or pockets between the two layers can cause warping during autoclaving, so it's advisable to dry the base layer thoroughly between washing and autoclaving.

An STL is included for those interested in attempting to print the lower base. This was attempted, but doing so restricts microscope viewing to the view holes rather than through the polycarbonate, and sacrifices the durability of the polycarbonate.

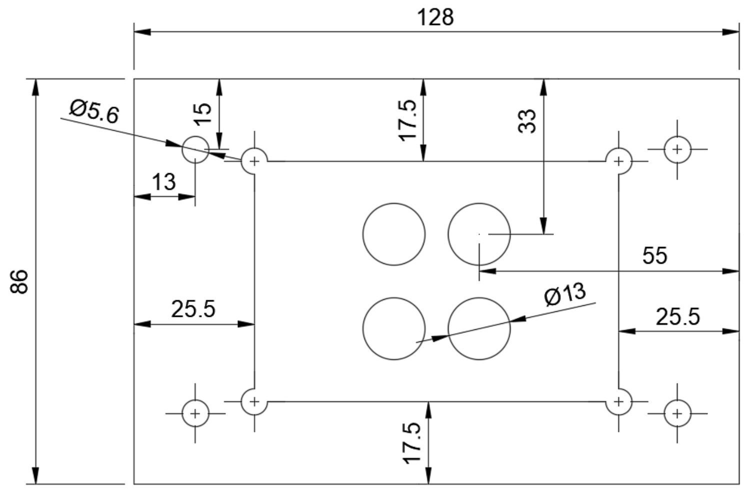

*Supp. Fig. 1.3: Dimensional diagram of 3D printable or single-piece support layer.*

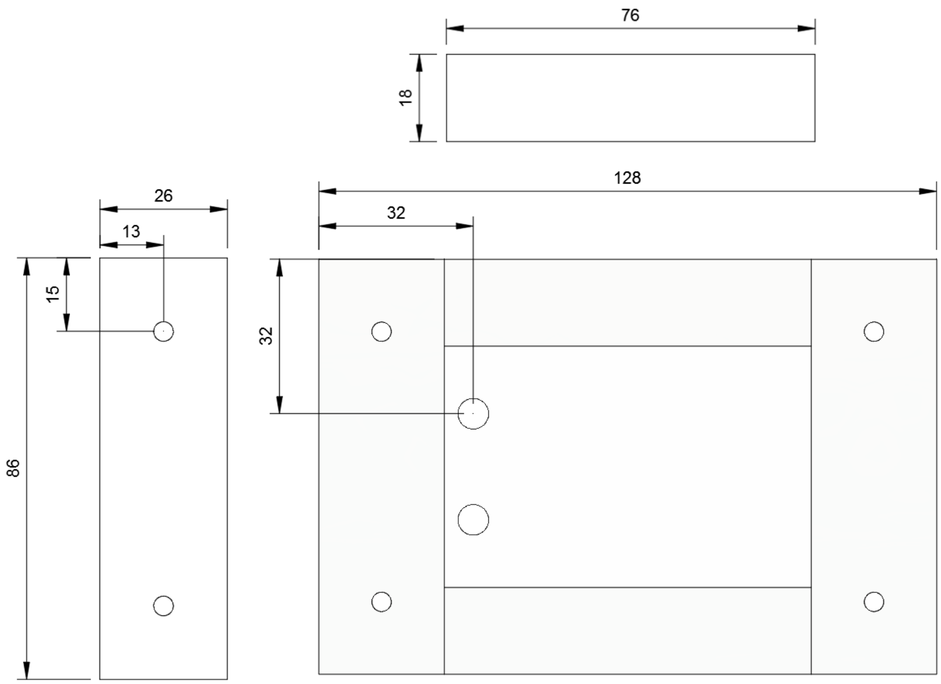

*Supp. Fig. 1.4: Dimensional drawing of multi-piece support layer*

*Supp. Table 1.1: Bill of Materials for Construct Housing*

| Item | Vendor | Catalog No. | Description |
|---|---|---|---|
| Polycarbonate sheets, 6x6 in x 1/4 in | McMaster | 8574K281 | For the lid layer, the microscope-compatible support layer, and the basic base layer |
| Polycarbonate sheets, 12x12 in x 1/16" thick | McMaster | 8574K24 | For microscope-compatible base layer |
| Polycarb. 1/4"-28 threaded Male Luer-Lok fitting | McMaster | 51525K241 | Fittings for Luer-Lok connection |
| M5 20mm Stainless-stell Hex head screw | McMaster | 91310A124 | For screwing on the lid |
| M5 Stainless-steel Wing Nut | McMaster | 94545A225 | For screwing on the lid |
| PDMS - KRAYDEN SYLGUARD 184 KIT | Fisher Sci. | nc9285739 | For creating gaskets |
| Scigrip 4 Nonwhitening Acrylic cement, 4 oz. | McMaster | 7517A1 | For solvent welding layers and affixing Luer-Lok ports |
| Precision Needle-Tip Squeeze Bottle | McMaster | 1902T341 | For applying acrylic cement |
| Generic PLA 3D printer filament | | | For printing a rigid mold or a master for creating a soft silicone mold |
| Mold Star 15 Slow, 2 lb trial size | Reynolds | Mold-Star 15 | For casting a soft silicone mold |
| SuperSeal & Ease Release 205 Combo Pack | Reynolds | Link | Release agent for mold making |
| X-acto knife | McMaster | 35435A11 | For cutting the master out of the soft silicone mold |

### 3. Make the Gasket

In both cases, the gasket is defined by the geometry of a construct (which determines the interior dimensions) and the geometry of the support layer outside the construct (which defines the exterior dimensions). In this example, the construct dimensions are 10 mm across and 46 mm long, and the exterior dimensions are 50 mm across and 76 mm long (which is the footprint of two standard microscope slides. In any case, once the gasket has been modeled in CAD, a mold is made, silicone is mixed, and the PDMS is poured into the mold to produce the gasket. However this molding process can be achieved in two ways: by directly 3D printing a rigid mold or by printing a master and then casting a soft mold out of silicone.

#### 3.1 Modeling the Gasket

The primary step of designing the gasket is to select the desired geometry of the final constructs and the dimensions of the support layer, after which features may be added to increase the quality of the gasket's seal. While many designs were tested for this project, two examples are provided below.

##### *3.1.1. Modeling a gasket for a 60 mm dish*

This design is extremely simple. It's outer dimensions allow it to sit snugly inside of a 60 mm culture dish. Be aware, though, that some dishes include small, barely noticeable geometric features on the inside or bottom which can prevent this gasket from sitting comfortably on the bottom of the dish. This design includes two carve-outs at either end which allow the housing to be filled with ink after it is fully assembled through a dedicated fill and vent port.

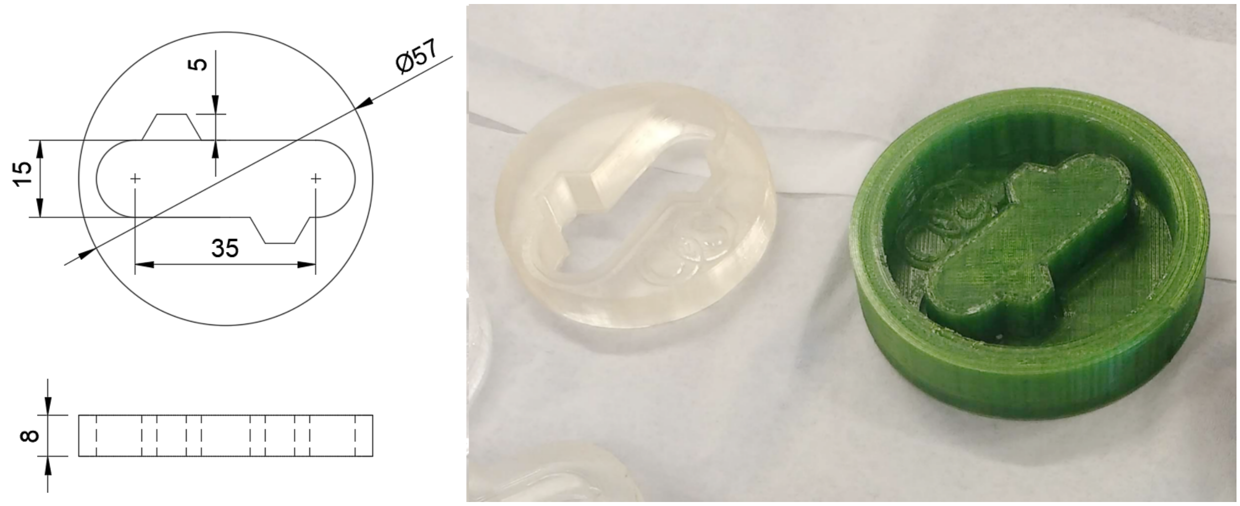

*Supp. Fig. 1.5: The dimensions of the simple gasket with the rigid 3D printed mold used to cast it.*

*3.1.2. Modeling a gasket for a a microscope-compatible housing*

The microscope-compatible gasket has exterior dimensions matching the footprint of two microscope slides and a height which equal to the ¼" thick polycarbonate support layer minus the height of a single microscope slide. The interior dimensions were selected to be be 46 mm long and 10 cm wide. Like the simple gasket above, this gasket has a raised lip designed in to improve the seal between the gasket and the microscope slide.

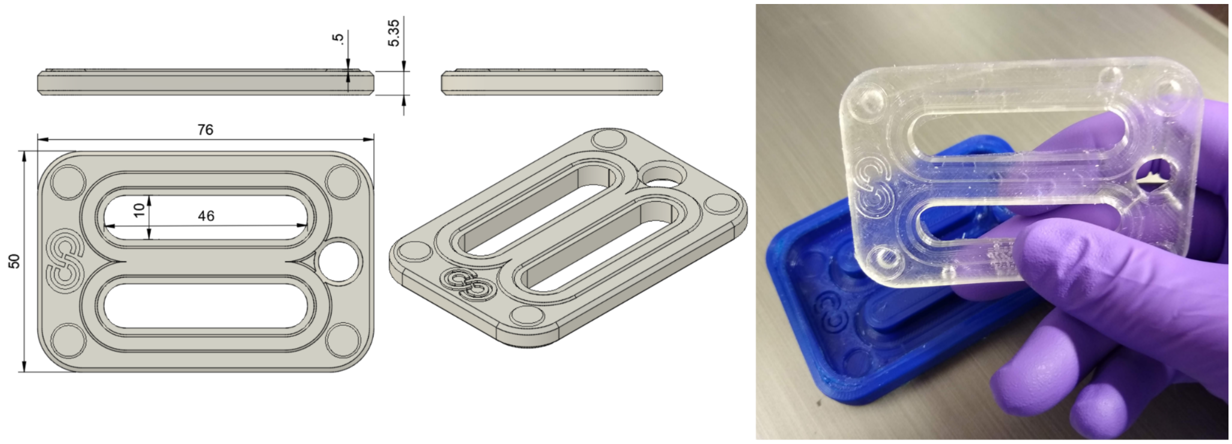

*Supp. Fig. 1.6: The dimensions of the gasket for the microscope compatible housing, with the rigid mold used to cast it.*

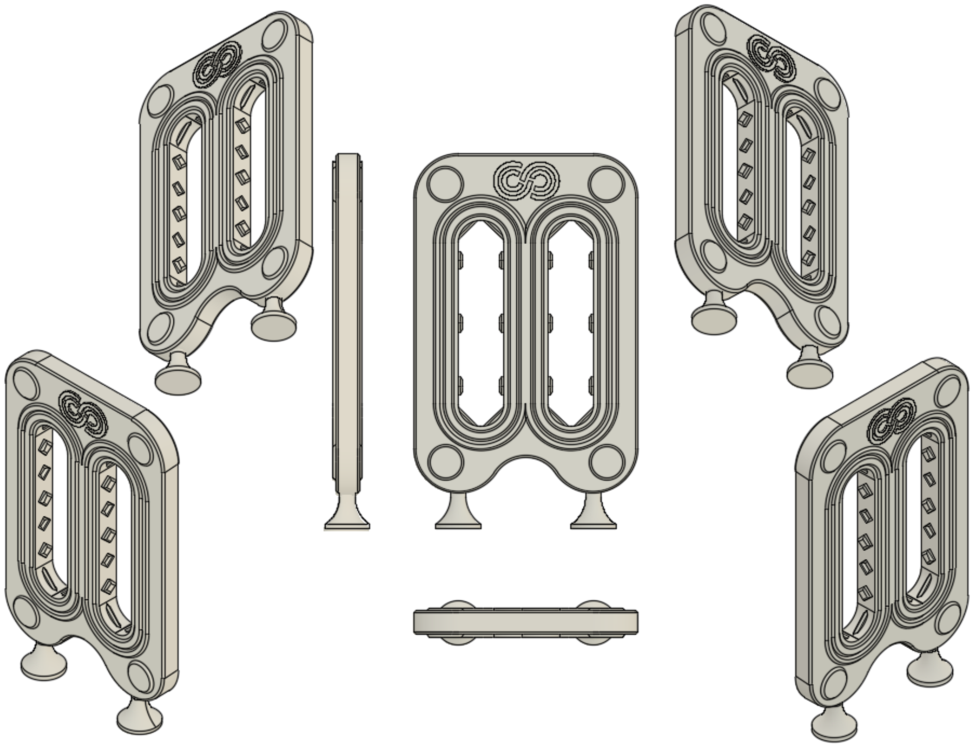

*Supp. Fig. 1.7: The microscope-compatible gasket with a lip on both sides*

A later version of this design integrated several additional features: most importantly, it included a raised lip on both sides of the gasket. A narrow channel was designed into these raised lips to allow for easier application of food safe silicone oil. Nubs were also modeled into the side walls of the channel, as it was found that constructs could occasionally lose attachment to the gasket and slide laterally along the walls. These nubs prevent lateral motion that might detach a construct from the silicone walls. This image above

Details on the process of designing the constructs and their scaffolds are covered in Supplementary Methods Part 3.

**The Challenge of Forming Reliable Seals**

Forming a reliable seal between the gasket and the glass slides below and the polycarbonate lid above is essential for maintaining bioprinted constructs within a perfusion housing. It is also deceptively challenging. Despite appearances, a flat silicone surface will not reliably prevent the escape or entrance of fluids under even modest pressure. Such a seal can be achieved, but it will not be nearly as reliable as a gasket with perimeter features. In order to avoid costly losses of time and effort, it is highly advisable to design gaskets with perimeter features like the one above.

Designing this feature on one side is easily achieved with a rigid mold, but designing this feature on both sides of the gasket likely requires a soft mold. Despite attempts, these geometries were unable to be obtained with rigid molding.

In either case, a stronger seal can be achieved by applying a continuous bead of food-safe silicone oil around the perimeter of the gasket, and by tightening the screws to compress the gasket aggressively.

#### 3.2 Preparing a Mold

##### *3.2.1. Printing a rigid mold in PLA*

In order to print a rigid mold, the master was subtracted from a larger body in Fusion360, and this resulting shape was 3D printed. The molds were printed using at least 5 walls and at least 5 floors and roofs in order to create a watertight mold. The silicone cures very slowly, and will leak if the mold is not watertight, which can occur in 3D prints if the walls and floors don't have enough layers and a high enough printing temperature for robust layer bonding.

##### *3.1.1. Casting a soft silicone mold around a PLA master*

The need for a lip on both sides of the gasket and a consistent height dimension eventually justified the additional work of producing a soft silicone mold. This mold was produced by printing a master of the gasket itself, with feet, and with a surrounding box. These feet and the surrounding box were glued to sheet of aluminum foil with ample hot glue to create a continuous seal. 400 mL Mold Star silicone was then mixed according the manufacturer's instructions and poured slowly into the box to full cover the printed masters, and then allowed several days to fully cure.

Once cured, a craft knife was used to slice a jagged-edge seam into the silicone mold. Making the seam jagged acts as indexing marks to help align the halves when preparing to use the mold. The masters were removed and mold release was applied to the interior of the new mold according to the manufacturer's instructions. This release was sprayed, brushed into all nooks, and allowed to dry. Then PDMS was prepared to cast the gasket.

#### 3.3 Casting PDMS in the mold

PDMS is prepared by mixing nine parts of silicone elastomer base with one part of the curing agent. This preparation was mixed with a power drill, as thorough mixing is essential, and pockets of unmixed elastomer will remain in corners unless they are carefully mixed in. The volumes of the gasket can be identified in your CAD program. The microscope-compatible gasket, which is the larger of the two, has a volume of 15 mL. To prepare a gasket, 20 mL of mixed silicone was stirred with a spatula attachment on a cordless drill for at least 60 seconds.

The mixed silicone was then spun in a centrifuge at high speed for thirty seconds to draw the entire mixture to the bottom of the conical tube and to remove bubbles.

The silicone was then deposited into the mold using a 50 mL serological pipet up to the height of the interior of the mold. When preparing a rigid mold, the height of the interior bodies acts as a level indicator. Covering this with a very thin layer is recommended to ensure consistent gasket height. This layer can be cut or peeled off easily after curing.

The mold containing the uncured silicone may take 3 days or more to fully cure at ambient room temperature, but it can be cured overnight in a typical labware drying oven. Longer times may be required for the soft mold.

##### 3.4 Demolding the gasket

After curing, when the gasket should be firm to the touch with little to know tackiness, at which point it can be demolded. Before demolding the gasket from a rigid mold it is recommended that the meniscus around the outside edge be gently sliced off with a craft knife. Otherwise, the meniscus may need to be trimmed off with scissors after demolding. The gasket can then be demolded by sliding a narrow spatula or knife blade between the silicone and the walls of the mold and then gently working this tool around the edge before carefully prying the gasket out of the mold.

After demolding, gaskets were ready for sterilization and use.

#### 4. Install the Luer-Lok Ports in the housing lid

The Luer-Lok ports are threaded with 1/4“-28 threads. They’re attached by drilling 7/32” holes, tapping them, and then solvent-welding the fittings in place with SciGrip 4 cement.

##### 4.1 Drill holes

Using the provided template (either on paper or 3D printed), drill 7/32” holes. Take great care to be precise. Tight dimensional tolerancing will improve the handling and repeatability of the overall process significantly, and minor but significant deviations when using hand tools are very common.

##### 4.2 Tap the Holes

Tap the holes with a 1/4“-28. Polycarbonate taps easily, without even requiring lubrication.

##### 4.3 Solvent Weld the Ports

Screw in the fittings (McMaster-Carr part 51525K241, 1/4"-28 threaded Polycarbonate Quick-Turn Tube Coupling Plugs) most but not all of the way in. Dispense solvent cement from a needle tip into the base of the threads. It will wick into the threads. Using a pliers if necessary, gently tighten the fittings into place completely. You can use a paper towel, tape, or aluminum foil to mask the surface of the polycarbonate to avoid drips. If drips occur, let them evaporate or dab with a paper towel rather than wiping them away. Wiping will only smear the solvent.

Flip the lid over and apply a bit of extra solvent cement to the threads from the underside of the lid to ensure a completely hermetic, molecular bond. Insufficient solvent welding will result in leaky ports at the welded junction. Even if the junction holds a seal initially, autoclaving stress can cause small leaks to appear over time. A well-cemented fitting, however, will remain leak-free indefinitely. Let the solvent evaporate overnight.

#### 5. Assemble the Flow System

The flow system consists of a pump, a reservoir, hoses to connect the flow, and inlet, outlet and venting needles, plus accessory tools. This setup is highly customizable. A syringe pump may be deemed more suitable to match the desired flow rate.

Assembly consists of cutting silicone tubes to the appropriate lengths sufficient to connect the housing to the appropriate reservoirs with enough additional length to accommodate the handling needed to connect the tubing to the peristaltic pump. For the arrangement pictured below, flow lines were cut to the following lengths:

*Table S1.2: Flow line lengths*

| Line | Length |
|---|---|
| Inflow line | 42 cm, with tube stop 23 cm from one end and 19 cm from other |
| Transfer line | 8 cm |
| Outflow line | 30 cm |

The pump used here has a minimum flow rate of ~ 7 µL/min when used with 1 mm diameter tubing. This comes out to 420 µL/hr or ~ 10 mL/day. Either a 50 mL or 100 mL media bottle offers a convenient size because their height is low enough for a 4” needle to reach the bottom. Constructs were fed with 20 mL of medium in cycled flow, with media replacement every other day. Media change can be performed using serological pipette in a hood, or through the septum caps using syringes and needles.

**Sourcing challenges**

The acquisition of the 4” long needles -- which are needed to draw from the bottom of the reservoir bottle -- and the peristaltic pump and pump head were among the most challenging items to source for this project. These needles are a controlled item in some areas, and this low-flow peristaltic pump is produced by a foreign manufacturer that can be difficult to reach.

*Table S1.3: Bill of Materials for perfusion system*

| Item | Vendor | Catalog No. |
|---|---|---|
| Low Flow Peristaltic Pump | Langer | BT100-2J |
| Peristaltic Pump head | Langer | DG-2-B |
| GL 45 open cap | VWR | 89059-782 |
| GL 32 cap, open with hole | VWR | 89059-780 |
| Septum for GL45 Cap, silicone/PTFE | VWR | 10099-412 |
| Septum for GL32 Cap, silicone | VWR | 10099-110 |
| Male-to-male Luer coupler, Pack of 20 | Sigma | 25064-U |
| 50ml PYREX media bottles, GL32 cap | VWR | 16157-068 |
| 100ml PYREX media bottles, GL45 cap | VWR | 16157-103 |
| Ismatec 3-stop, silicone tubing, 1.14 mm ID | Cole-Parmer | EW-95603-30 |
| 0.2 um Syringe Filters, w Acrylic Housing | VWR | 28145-477 |
| 10 mL syringe | VWR | 76290-382 |
| 20 mL Syringes | VWR | 76290-384 |
| 30 mL Syringes | VWR | 76290-386 |
| 4” 22 G Hypodermic needles | Fisher | 14-817-102 |
| 4” 16 G Hypodermic needle | Grainger | 19G370 |
| 22 G Needles | Fisher | 14-840-91 |
| 18 G Dispensing Needle, 1/2" Needle Length | McMaster | 75165A675 |

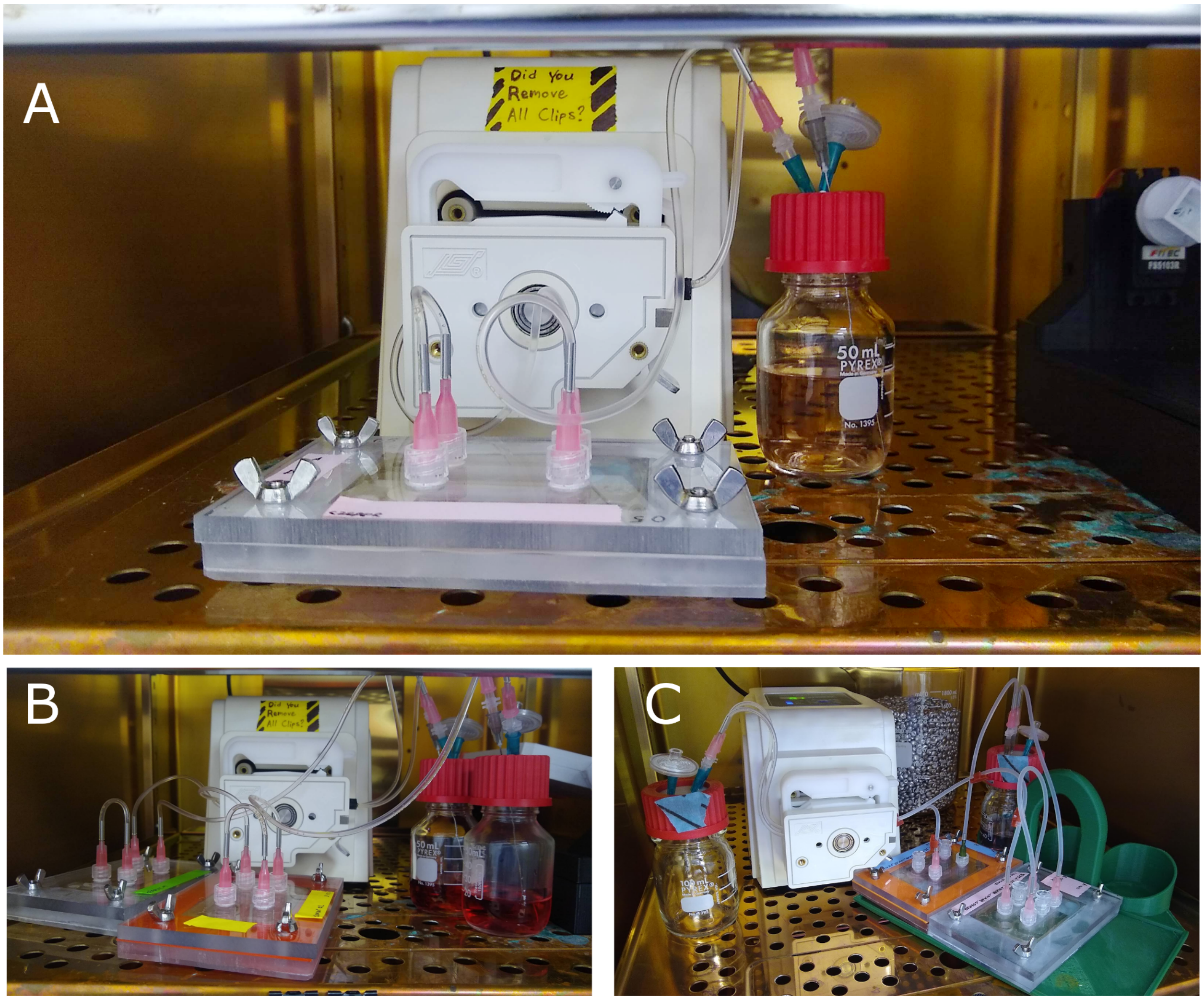

*Supp. Fig. 1.8: The assembled flow system. A) A standard cycled flow using a single 50 mL reservoir feeding two constructs through serial flow. B) Two housings, each with two constructs fed in series, fed independently from two reservoirs driven by one pump. C) Two housings with a total of three constructs fed in parallel from a single reservoir and outflowing into a separate output reservoir. The foreground housing is feeding two constructs in parallel.*

### II. Building and Integrating Compact Mechanical Extruder

**1. Designing the Compact Cartridge Extruder 17**
1.1 Syringe dimensions 17
**2. Modeling the Coupler 19**
**3. Integrating the Compact Cartridge Extruder into a Printer 22**
3.1 Mechanically integrating the extruder 22
3.2 Updating the printer's firmware 22
3.3 Electrically wiring in the new extruder 24
**4. Integrate a Stage Bracket 25**
4.1 Designing a Stage Bracket 25
**5. Setting up a Printer Control 26**

The compact cartridge extruder allows the user to rapidly load inks via a syringe cartrige that can be prepared elsewhere or elsewhen, and then printed without the loss of material associated with a tube connecting a distal syringe to the nozzle. STL files of two variants of this printhead are included with this guide, however it's likely users' exact dimensions will vary, particularly due to differences in the mounting points on different printers. For this reason, the design process is outlined here.

#### 1. Designing the Compact Cartridge Extruder

The cartridge extruder specifications are defined by three constraints: the dimensions of the syringe, the dimensions of the printer's mounting points, and the dimensions of the motor.

##### 1.1 Syringe dimensions

The Sareen lab uses 5 mL Syringes by BD with Luer-Lok Tip (BD REF 309646, Fisher Cat. #14-829-45). These syringes have a 13.5 mm outer diameter, so our socket is designed with a 14 mm inner diameter. To create a snap fit, we open the socket wide enough to allow a user to squeeze the syringe in without too much strain while keeping it narrow enough that the syringe doesn't fall out. In our printer, the opening was 13 mm wide, or ~136 degrees. The socket snap-firmness can also be adjusted by increasing or decreasing its length. Our pinch region is 9.5 mm tall.

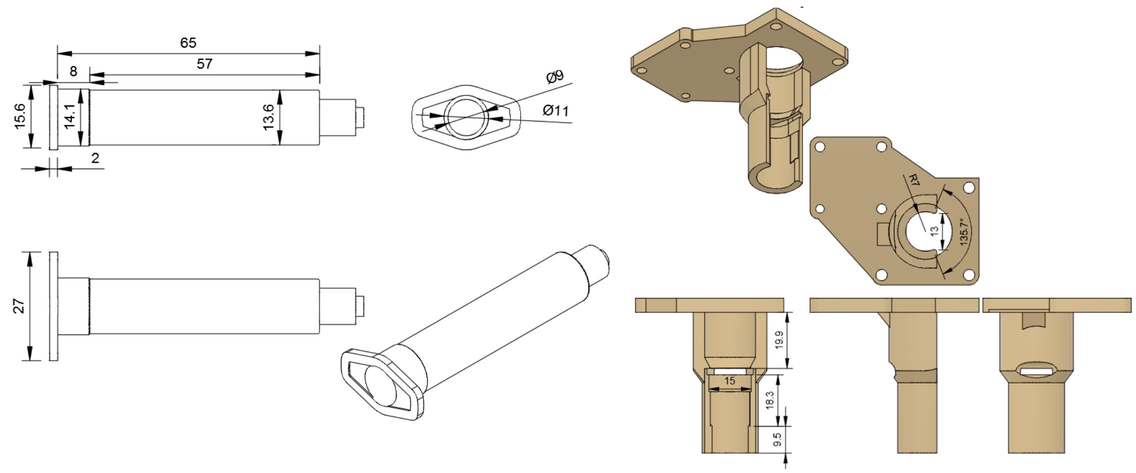

*Supp. Fig. 2.1: Printhead Carriage Mounting Points*

In order to replace the plastic-extruding hotend on a conventional FFF 3D printer, the replacement printhead must attach to whatever mounting points are designed into the printer. The Sareen lab used a Velleman Vertex, which is a reliable, affordable, hackable platform. Whatever base kit you use, prioritize hackability, as this means it's easier to find existing instructions for mounting modified printheads. On the Vertex, the printhead mounts to four 3 mm holes in a square configuration 22 mm apart. Make sure the printhead doesn't collide with the gantry rods or the sides of the printer when homing.

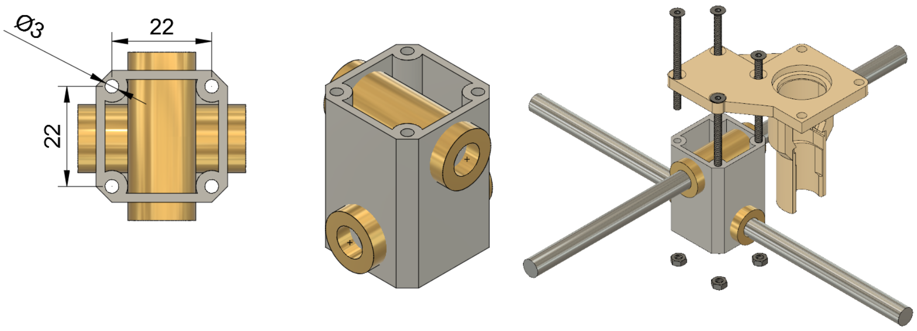

*Supp. Fig. 2.2: Motor Mounting Points*

In order to drive the syringe, we used a compact linear actuator that doesn't require rails. There are many guides online which describe ways to translate a stepper motor into a linear actuator by attaching a stepper motor to a threaded shaft, but the size, weight, and complexity of the syringe pump can be substantially reduced by using a stepper motor with a built in threaded rod. Be aware that at the time of publication, we know of only one product available from one source: ROB-10848 from SparkFun. If this isn't available, you may need to build a similar item by disassembling a stepper motor and installing a nut within the center of the rotating shaft hub.

This stepper motor is a NEMA 15 motor with M3 threaded holes in a square 31 mm apart on each side. It also has a circle 22 mm in diameter that extends out 2 mm.

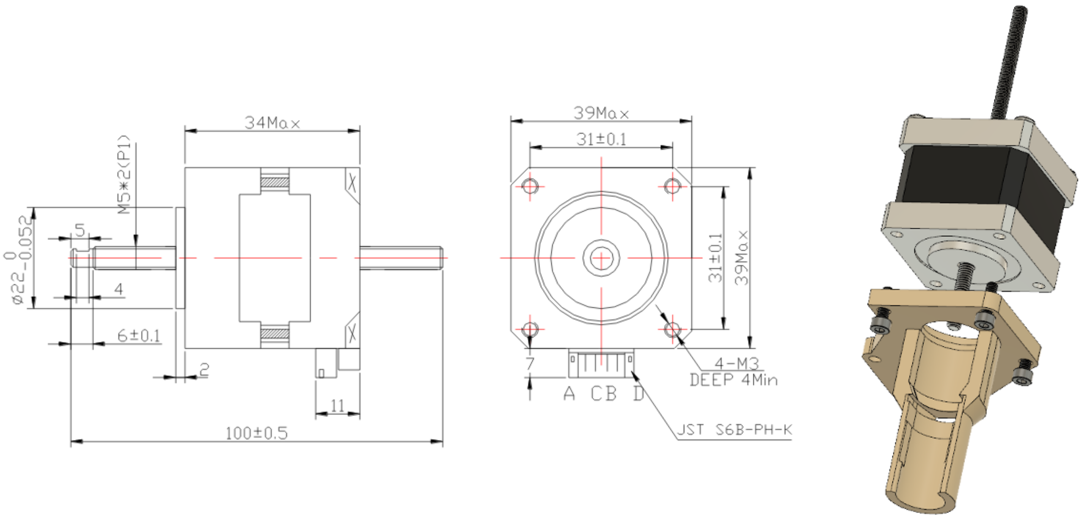

*Supp. Fig. 2.3: Dimensions and design of compact linear stepper motor*

### 2. Modeling the Coupler

The coupler is a two-piece coupling that mates the shaft of the linear actuator to the plunger of the syringe using a bulb and socket that snap fit together. A manual loader with a socket is used to load the syringe before printing.

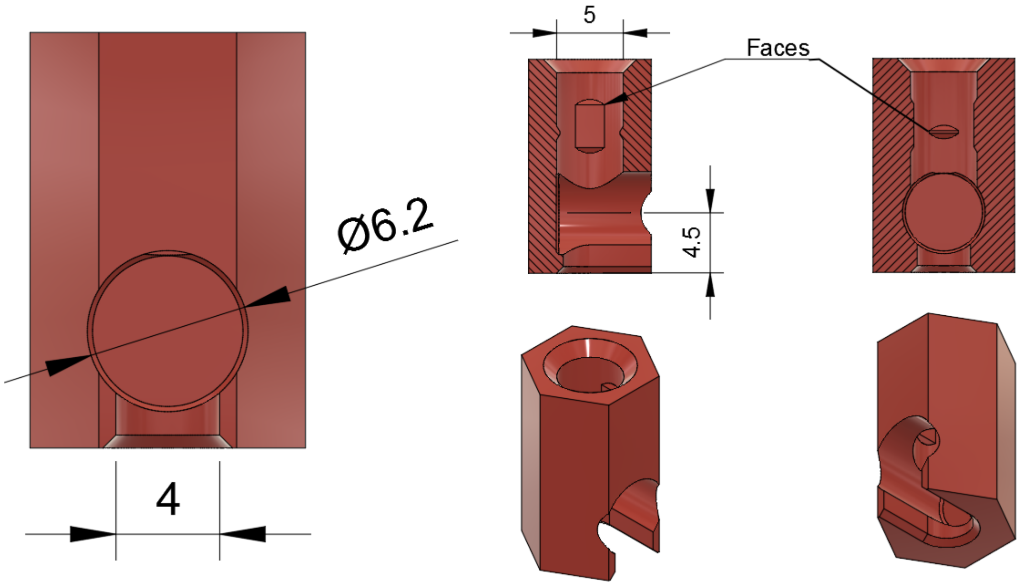

*Supp. Fig. 2.4: The Motor-side coupler*

The motor-side coupler consists of a socket on the bottom that receives the plunger-side coupler and a hole that engages with the threaded shaft.  The socket size used by the Sareen lab is 6.2 mm in diameter and 4.5 mm above the bottom of the motor-side coupler, with a 4 mm wide neck.  The screw hole is 5 mm in diameter with faces on four sides that give the threaded rod material to bite into.  The size of the hole could be adjusted to fit the thread, or the plastic could have threads tapped into it. This early design worked well enough that changes were never required.

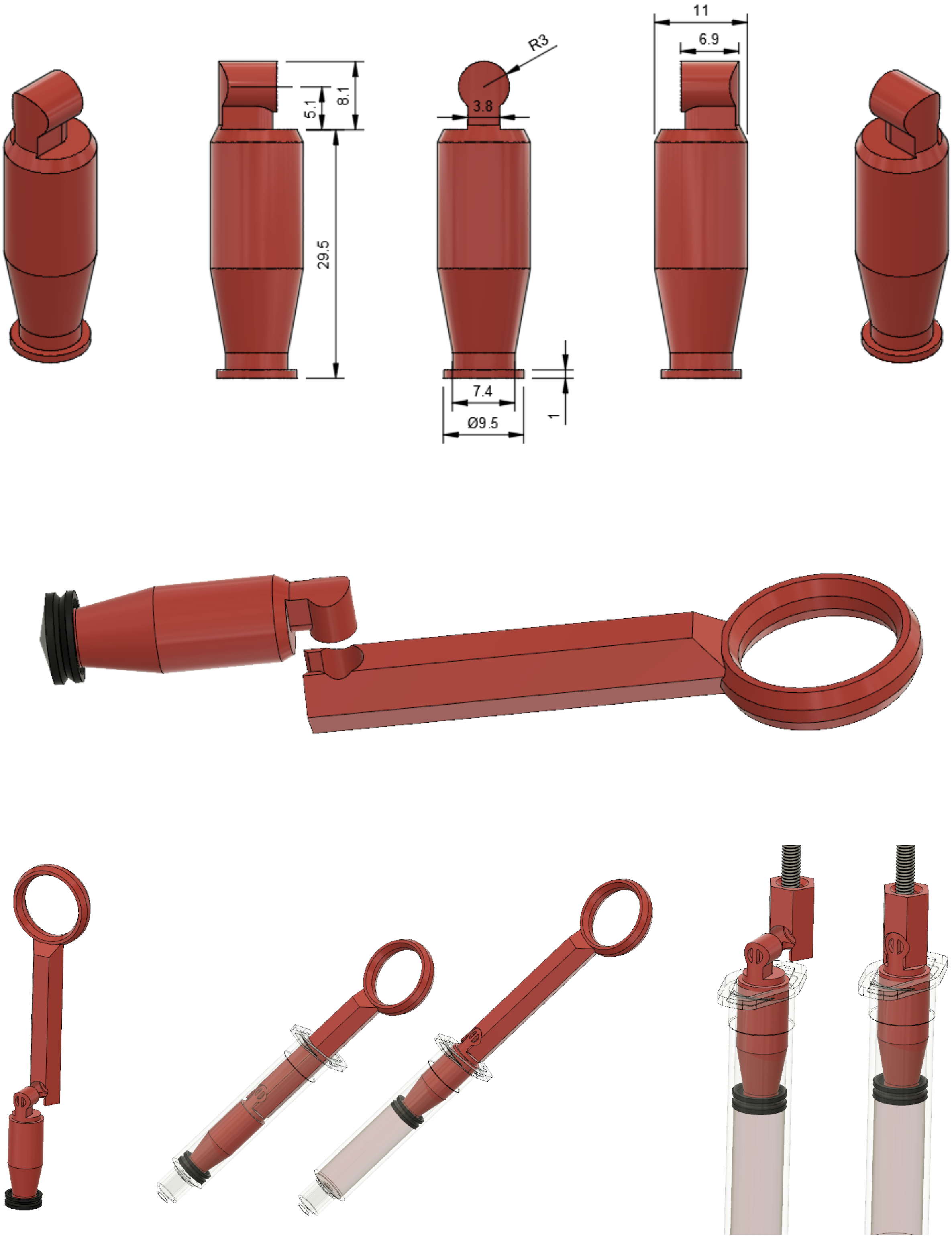

*Supp. Fig. 2.5 The Plunger-side coupler and its use with the manual loader and the motor-side coupler*

The plunger-side coupler consists of a bulb on top that couples with the motor-side coupler and a foot that inserts into the syringe. The bulb can be any size so long as it's designed to interface with the motor-side coupler.  The Sareen lab used a bulb that is 6 mm in diameter, 5.1 mm above the top of the plunger, and attached to the plunger with a neck 3.8 mm across.

The foot of the plunger is 9.5 mm in diameter and 1 mm tall. This plunger substitutes for the plunger that comes with the syringe, but it uses the same rubber foot.  The 9.5 x 1 mm cap fits the foot well, and maintains a snug fit inside of the syringe.  If necessary, the foot cap should be enlarged for shrunk to maintain a tight seal with the right level of friction inside the syringe. Keep in mind that the threaded-shaft motor turns a screw that drives the shaft if the shaft is unable to rotate. If the shaft rotates freely, 0% of its movement is along its axis, and if it is unable to rotate than 100% of its movement is axial. The friction within the syringe is higher rotationally than axially, so the motor travels axially if the foot is sized appropriately.  If the cap is rotating than the friction likely needs adjusted by increasing or reducing the diameter of the foot inside the rubber cap.

The length of the shaft of the plunger determines the starting volume of the syringe, since the plunger must be drawn back to its loading position in order to load the syringe into the 3D bioprinter. A ~30 mm shaft as shown below will draw ~4 mL of bioink into a 5 mL syringe.  STL files for other size plungers are included in supplemental files in order to allow users to load syringes with no more bioink than necessary to avoid wasting valuable reagents.

The manual syringe loader is a tool used to draw the replacement plunger in order to load syringes.  It consists of a socket of the same dimensions as the one on the motor-side coupler, a shaft, and a ring.

### 3. Integrating the Compact Cartridge Extruder into a Printer

Integrating the new extruder into a printer requires mechanically mounting the extruder, wiring the electronics, and updating the printer's firmware accordingly.

#### 3.1 Mechanically integrating the extruder

Mechanical installation may vary by printer, but in all cases will likely require bolting the new extruder onto the printhead carriage.  The new extruder should be designed with mounting holes that match the printer's, making this a fairly easy step.

The new extruder will likely extend further downward from the carriage than the previous extruder, so the Z-axis home switch will likely need to be moved accordingly.

#### 3.2 Updating the printer's firmware

A 3D printer's firmware interprets instructions and acts on them. For instance, the firmware configuration includes the number of steps a stepper motor must move to travel one mm and will interpret a command to move 3 mm by instructing the motor to turn the correct stepper motor by the correct number of steps. Most printers use an open source firmware called Marlin. Marlin is coded in C++ and can be edited and compiled in Arduino IDE.

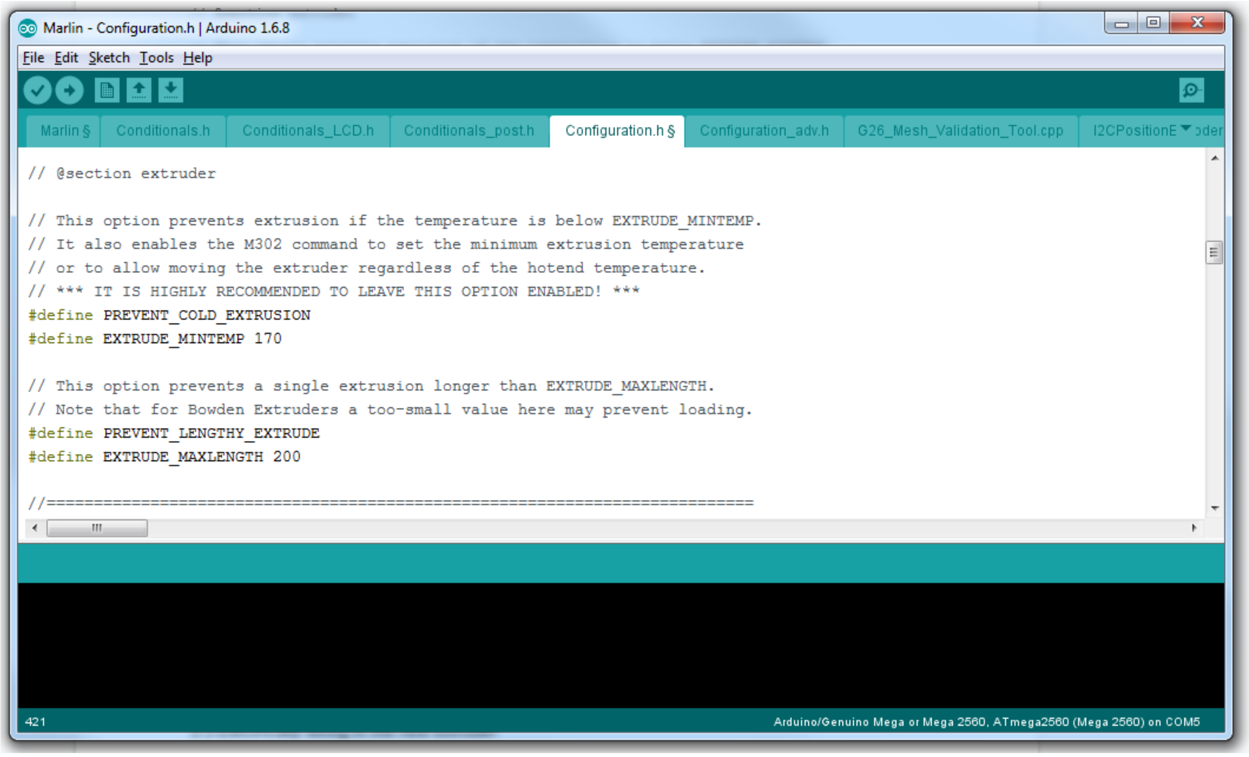

*Supp. Fig. 2.6: The key change to make within the firmware is to disable "PREVENT_COLD_EXTRUSION".*

Users can either modify the firmware that comes with their printer or they can configure the latest version of Marlin completely. Configuring the existing software is recommended, as it will save a lot of time. Configuring Marlin from the latest version will require a user to specify the dimensions of the printer, the position and polarity of end-stops, the number of extruders, the language for sending information to the display board, and many other things that are already set in the firmware that comes with a printer.  In our case, however, the firmware which came with the Velleman Vertex was of an older version that wasn't editable in the most recent version of Arduino IDE and was less user-friendly in the formatting of its configuration file.

Whichever route you take, the key change required is to disable cold extrusion protection. A conventional FFF 3D printer can't extrude filament unless the hotend is hot enough to melt the filament, so the default configuration in Marlin will ignore any instruction to move the extruder motor unless the measured temperature is above a safe level. The syringe extruder described here has no thermal control, so it has no thermister measuring temperature and will never reach the minimum temperature, so the printer can't operate unless cold extrusion prevention is disabled. To disable cold extrusion prevention open the configuration tab in the Marlin firmware and find the relevant section below. In my version it begins at line 415.

```
// @section extruder

// This option prevents extrusion if the temperature is below EXTRUDE_MINTEMP.
// It also enables the M302 command to set the minimum extrusion temperature
// or to allow moving the extruder regardless of the hotend temperature.
// *** IT IS HIGHLY RECOMMENDED TO LEAVE THIS OPTION ENABLED! ***
#define PREVENT_COLD_EXTRUSION
#define EXTRUDE_MINTEMP 170
```

Comment out the line "#define PREVENT_COLD_EXTRUSION" by adding two slashes ("//"). It should now look like this:

```
// @section extruder

// This option prevents extrusion if the temperature is below EXTRUDE_MINTEMP.
// It also enables the M302 command to set the minimum extrusion temperature
// or to allow moving the extruder regardless of the hotend temperature.
// *** IT IS HIGHLY RECOMMENDED TO LEAVE THIS OPTION ENABLED! ***
//#define PREVENT_COLD_EXTRUSION
#define EXTRUDE_MINTEMP 170
```

You'll also want to change the steps per mm of the extruder motor to reflect that the extruder motor moves 100 steps for ever mm.

##### 3.3 Electrically wiring in the new extruder

The new extruder uses a stepper motor that should be similar to the native motor it's replacing. This is a bi-polar stepper motor with four wires, which connect from the stepper motor driver board on the printer's motherboard to the stepper motor. In order to operate though, the wires coming out of the stepper motor must be connected to the appropriate pins on the printer's main board. The process for identifying the correct wiring is described on the RepRap project wiki. The easiest way to find the correct connections is through trial and error.

To do so, cut a male-female jumper wire in half and strip 20 mm of insulation from each, then solder male jumper wire ends to each of the four wires coming out of the motor. Solder female jumper wires onto the four wires coming from the main board. If the male jumper wires fit into the existing motor connector then soldering on female jumper connector ends is unnecessary.

Plug the new extruder motor's four wires into the four motor wires coming from the main board in any arrangement.  Send a command to the motor to test.  The specific menu will vary based on the printer, but will likely be under "Prepare" (or "Control") and then "Move Axis" and "Move Extruder" or something similar.

**Stepper Motor Specification Considerations**

Ideally, the stepper motor would share the electrical specs as the motor it replaces and conform to the electrical specs of the stepper driver board that translates movement commands from the mainboard to the stepper motor. Because alternatives were unavailable, we used the SparkFun ROB-10848, which is rated for 12V and 0.4A, even though the original stepper motors on the Velleman K8400 3D printer we modified were 3.1V 2.5A motors. These were the specifications expected by the Velleman printer stepper motor boards. It used VM8400DB stepper motor boards, which used the popular DRV8825 (Texas Instruments) motor driver chip. Fortunately, despite being designed to operate at a higher voltage, the SparkFun linear stepper motor can function fine in this setup. The consequence of operating this stepper motor below its intended voltage is a reduction in maximum speed and torque, and as the system does not require operation anywhere near the motor's max speed or torque, the reduction in performance is of no consequence. The difference in resistance between this motor and the default one could lead to higher operating temperatures, but in practice this has never raised the temperature of the motor above its defined operating temperature (80°C above ambient temperature, with a maximum ambient temperature of 50°C).

### 4. Integrate a Stage Bracket

Whatever is being printed onto will need to be held in place to keep it from sliding. The Sareen lab has printed in 100 mm dishes, 60 mm dishes, and on standard microscope slides.  For each of these, a bracket is needed that holds the print surface in place.

#### 4.1 Designing a Stage Bracket

The Sareen lab 3D printed a stage bracket designed to hold the print surface snugly in place while printing, which was glued with cyanoacrylate super glue to ta 3D printed plastic adapter that snapped onto the bed. The Vertex had a metal frame against which a glass build plate was supposed to be clipped, but without the glass build plate it was easy to design a snap-on adapter that clipped firmly onto the metal frame.  Brackets fitted to he desired print surface -- a 100 mm dish, a 60 mm dish, or a microscope slide -- were then printed and glued onto the snap-on build plate attachment.

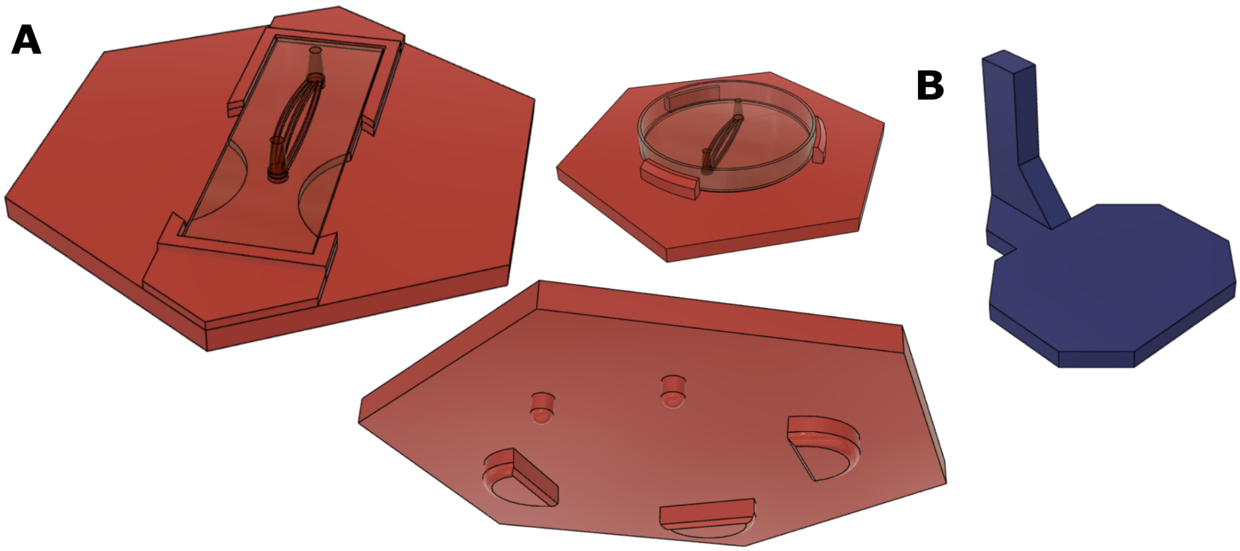

*Supp. Fig. 2.7: Printer stage accessories. A) A stage bracket adapter snaps onto the frame of the Vertex printer and allows for the placement of print surfaces. B) A z-switch extender triggers the z-axis endstop at a new height to accommodate the stage bracket adapter.*

Additionally, adding a stage bracket will change the height at which the nozzle will reach the stage, so the Z-switch will need to be modified so that it triggers a home signal when the nozzle reaches the bed instead and doesn't allow the nozzle to crash into the raised bed. On the Velleman, we printed an extension for the tab which contacts the Z-home switch to trigger the home switch and appropriate distance higher than before.

#### 5. Setting up a Printer Control

In order to execute a print, a g-code file can be run from the printer directly or over USB. We found USB control far more convenient. This was performed by running the free 3D printer control software Repetier-Host on a dedicated laptop which sat next to the benchtop PCR hood in which the printer resided.. Though technically no-longer open source as of 2014, Repetier-host was an adaptable, user-friendly interface for a custom or modified printer. It can run Cura within the program to slice files or load g-code files prepared in the Cura standalone application.

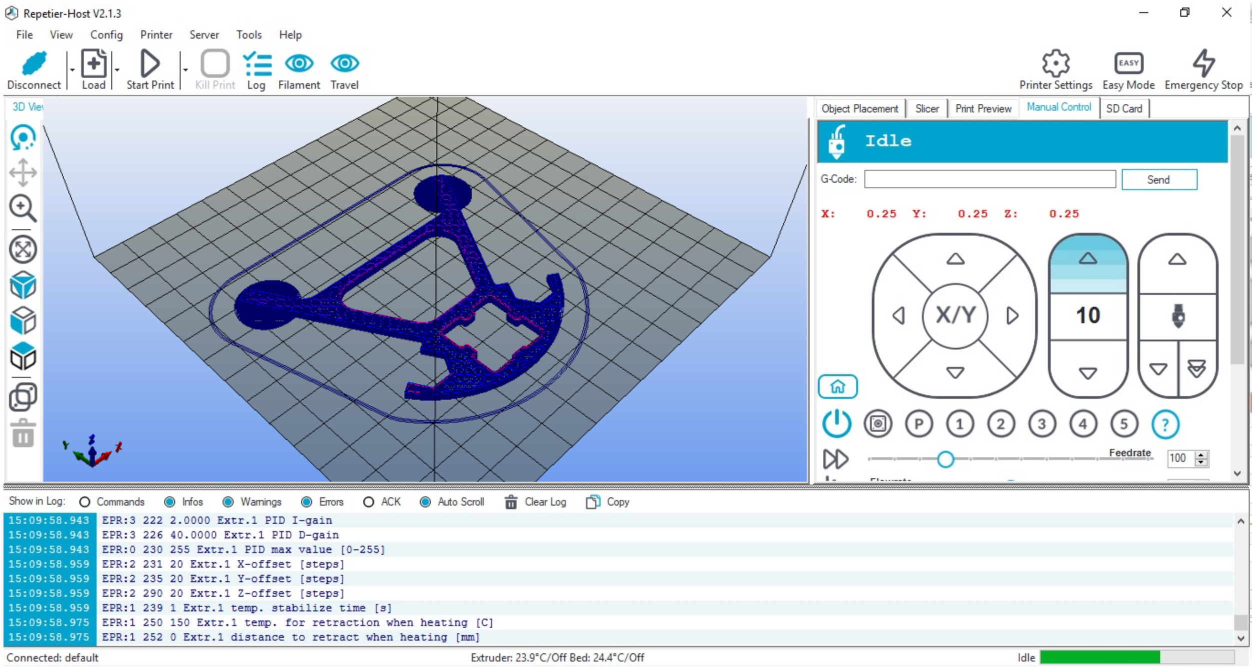

*Supp. Fig. 2.8: The manual control panel within Repetier-Host provides crucial functionality for setting up and executing prints.*

In addition to allowing a user to issue g-code commands and view the printer's log in real time, Repetier allows users to program g-code macros. We set up our printer with the following buttons:

| Button | Command |
|---|---|
| 1 | ; Move to start position to attach needle and prime flow:<br>g1 x100 y50<br><br>; Home the print head, then move it to its first position to load a cartridge and/or attach a needle<br><br>M206 x1 y1.5 ; M206 sets offsets. Use this to correct minor mismatches between the placement in the slicing software and the placement on the print bed<br><br>G28 X ; Home the X axis<br>G28 Y ; Home the Y axis<br>G1 X185 Y183 ; Move to a position where a loaded needle won't collide with the bed while homing in the Z direction<br><br>G28 Z ; Home the Z axis<br><br>G1 Z20 ; Move the bed down to create clearance<br>G1 X170 Y70 ; Move the print head into a convenient location to insert a print cartridge or change a needle.<br><br>M92 E3000 ; M92 sets the steps per unit for a motor. Here, the steps per mm for the extrusion motor are set to 3000. |
| 2 | ; Move the nozzle to what should be a known reference point in all axes |

| | |
|---|---|
| | G1 X113.564 Y90.936 ; The tip of the nozzle is moved to what should be the center of the near tower.<br>G1 Z10 ; The tip of the nozzle is moved to what should be less than 1 mm above its starting height. |
| 3 | ; After adjusting if necessary, the current position is confirmed to match the reference point.<br><br>G92 Z0 G92 X113.564 Y90.936 ; The current nozzle position is declared to be the positions listed after the G92 command<br><br>G1 Z30 F3000 ; The bed is lowered to provide clearance<br>G1 X50 Y90 F3000 ; The nozzle is moved away from the reference point so that it's no longer above the print surface while priming flow |
| 4 | ; This command also moves the nozzle into position to prime flow, but without rewriting the nozzle position in order to prime flow when position correction is not desired<br><br>G1 Z30 F3000 ; The bed is lowered to provide clearance<br>G1 X60 Y00 F3000 ; The nozzle is moved away from the reference point so that it's no longer above the print surface while priming flow. The position is slightly different from command script 3 in order to make its behavior obvious if button 4 is pressed after 3. |
| 5 | ; Flow is primed before printing<br><br>G92 E0 ; The position of the extrusion motor is set to 0.<br>G1 E0.2 ; The extrusion motor is advanced by 0.2 mm. Pressing this repeatedly slowly fills the syringe with ink.<br><br>G1 E0.1 ; The extrusion motor is retracted by 0.1 mm after each extrusion. This reduces the occurrence of oozing which can otherwise continue after priming the nozzle is complete. |

At this point, the printer should be complete and ready to print gels on a print surface. But first, a gcode file must be generated. See Supplementary methods, Part 3 for designing scaffolds and preparing printable G-Code files.

### III. Designing Scaffolds and Preparing Printable G-Code Files

**1. Designing and casting Silicone Gaskets** 30
- 1.1 Modeling the gasket 30
- 1.2 Modeling the printable gasket mold 31

**2. Designing the printable scaffold** 32
- 2.2 Generating printable files 33
- 2.3  Optimizing the model for printing 34
- 2.4 Refining the print conditions 36

The creation of a 3D bioprinted construct begins with its conception in a computer aided design (CAD) program. When drafting a design, however, it helps to understand the hands-on production process in order to design with production in mind. That production process consists of the marriage of two separate processes. The exterior geometry of the construct is produced when the surrounding matrix is poured into a mold defined by the silicone gasket. The interior geometry is produced by printing the sacrificial scaffold.

It's difficult to overstate the degree to which this process was under-described in the scientific literature available at the start of this project. Concisely, the channels described here consist of two vertical towers connected by one or more single extrusions strung between them, and these print commands are obtained by modeling such a shape and then adjusting settings in the Ultimaker Cura Slicer to translate this shape into movements which for a gel to be gently laid down without breaking.

### 1. Designing and casting Silicone Gaskets

#### 1.1 Modeling the gasket

Detailed methods for designing, modeling, and producing a PDMS gasket are described in greater detail in Supplementary Methods 1, but are revisited here to clarify the process of designing the scaffold which will sit inside the gasket. The first step of designing the silicone gasket is to select the outer dimensions of the construct. The model construct described here is 46 mm long, 10 mm wide, and 5.35 mm tall. This gives it a volume of 2.35 mL. This means that the PDMS gasket should be 5.35 mm tall and contain cutouts in the shape and size that the construct is meant to occupy.

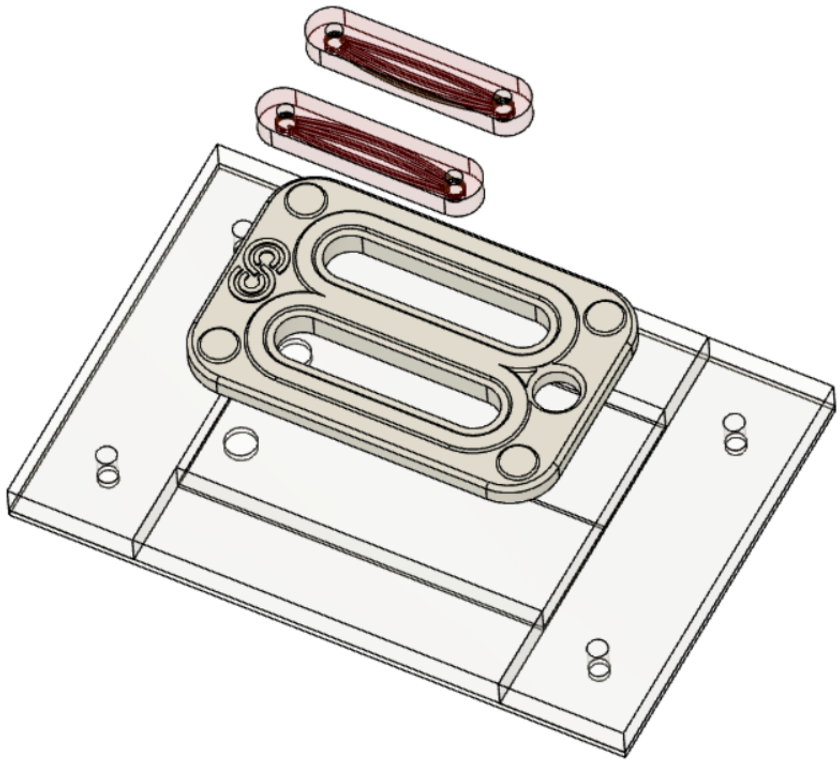

*Supp. Fig. 3.1: An exploded render of the housing base, gasket, and two constructs, with internal channels visible within the constructs.*

The outside of the PDMS gasket is 76 mm long and 50 mm wide, which matches the size of the perfusion housing's bottom half. The edges were chamfered and a perimeter was raised 0.5 mm around the top edges of the constructs in order to create slightly more robust seal when compressed.

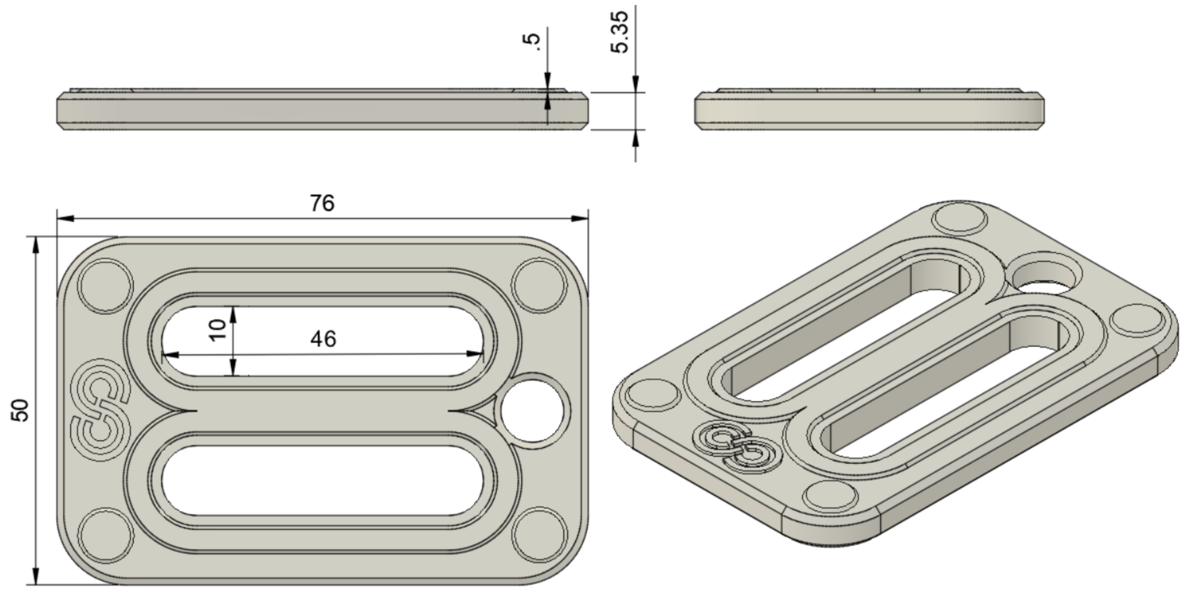

*Supp. Fig. 3.2: The dimensions of a PDMS gasket*

#### 1.2 Modeling the printable gasket mold

After the CAD model of the silicone gasket is finished, a mold file was prepared by subtracting the silicone gasket model from a larger block in Fusion360 using the Combine>Subtract function. The mold was left open on the top. Attempts were made to print two-part molds, a sufficient seal to prevent the PDMS from leaking out while also avoiding bubbles and allowing for the part to be demolded stymied these efforts. Later, these geometries were achieved using a soft silicone molding technique.

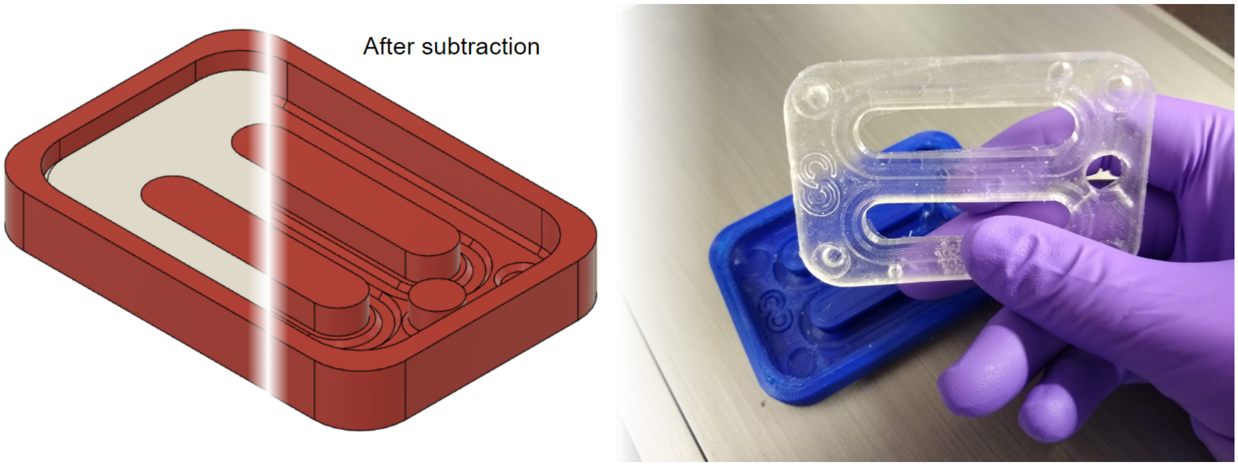

*Supp. Fig. 3.3: A rigid mold is modeled by subtracting the desired geometry from a larger body to create a printable plastic mold. The PDMS is then mixed and cast to produce a gasket.*

#### 2. Designing the printable scaffold

The printable scaffold is the heart of the bioprinting process. Its production consists of modeling the channels and then converting the model into machine instructions.

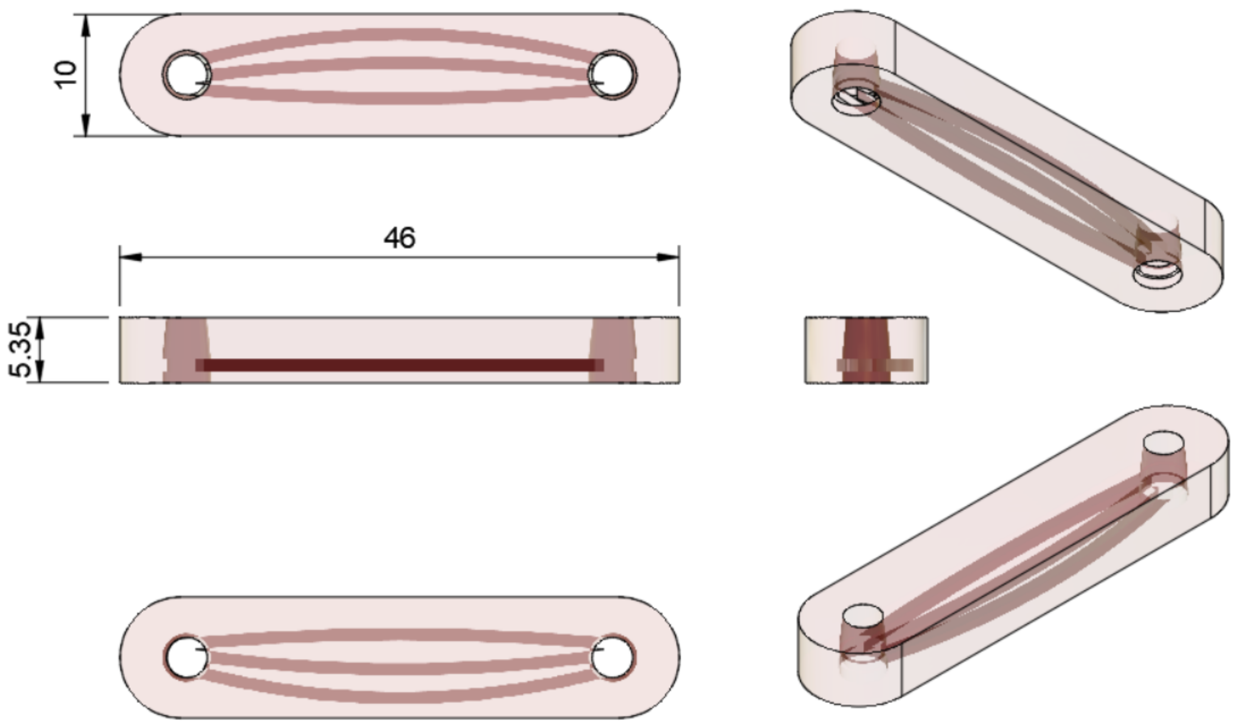

*Supp. Fig. 3.4: Modeling the basic scaffold*

The standard scaffold described here consists of two towers with channels connecting them. The width and number of the channels can take many forms depending on the interior surface area and ease of printing desired. Depending on the use, the amount of distance between the lowest channels and the bottom of the construct may need to be greater or lesser. Creating 1 mm or greater distance has the benefit of ensuring that diffusion isn't aided by spreading out of the channel underneath the construct, while having lesser or no distance improves the ease of visualizing the channels within the perfusion housing.

The basic design process consisted of drawing the channels as 2D layers and extruding them upward by their intended width, then repeating for each layer. This method of modeling vascular channels was designed specifically to cater to the process of converting a model to a printable toolpath by designing each layer of the same height that would then be used by the printer to create individual slices.

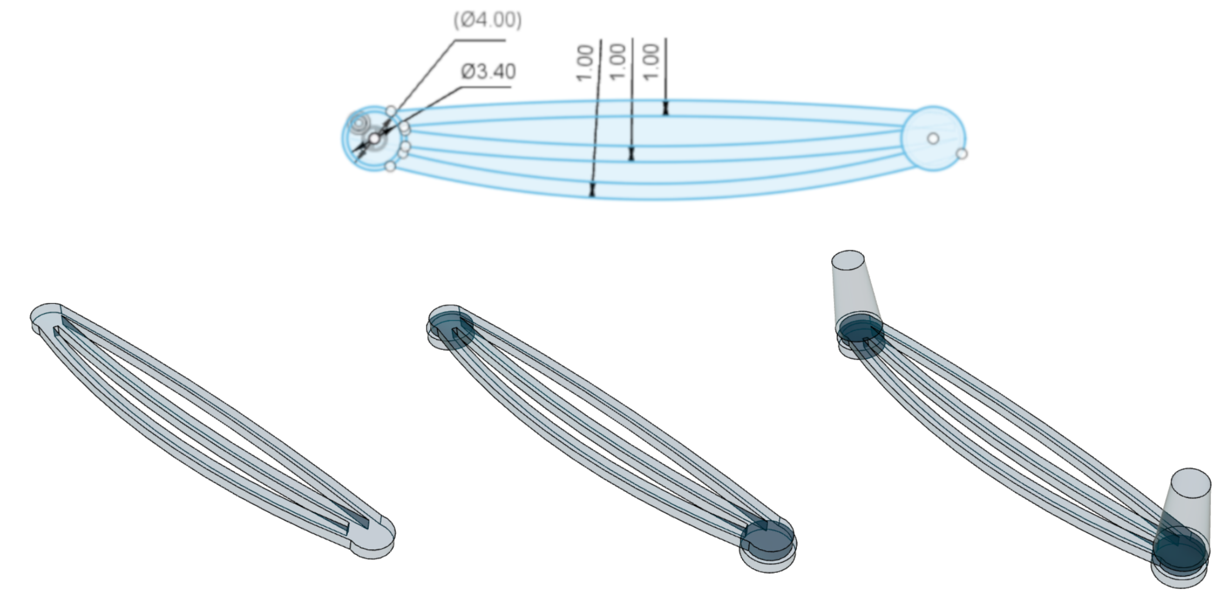

*Supp. Fig. 3.5: The 2D dimensions of a three-channel scaffold were drawn, then extended up by the same height as the slicer would be instructed to slice the model (in this case, 1 mm).*

#### 2.2 Generating printable files

In conventional fused deposition modeling (FDM) printing, a 3D object is sliced into its constitutive layers by a slicing program which then identifies a path on which to drag the print nozzle to trace out the 2D shape needed to print each layer. The resulting toolpath is saved as a printable .gcode file. This process is not designed to extrude a single thin filament by itself, however. Each layer is designed to be a closed shape with an interior and an exterior, which means that any shape thinner than twice the width of the extrusion nozzle can't be printed under standard settings. The printer instead either omits it or attempts to print a perimeter in which two sides of a rectangle overlap each other.

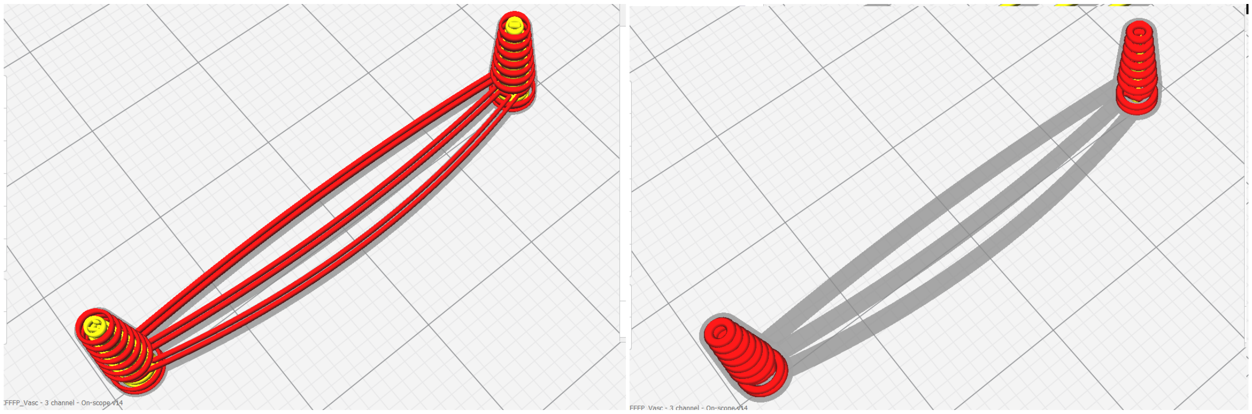

*Supp. Fig. 3.6: Outer walls are drawn in red. On the left, the toolpath traces each channel twice, destroying the scaffold. On the right, the channels are left out entirely.*

In previously published bioprinting papers in which a bioprinter is used to print a single thin extrusion, the toolpath is usually developed using entirely custom code in Matlab or a similar program. In order to provide a broadly user-friendly solution, we found a hack in the popular and free Ultimaker Cura slicing software that induced it to draw single-line channels: print the whole model only as infill.

Under normal operation, the slicer breaks each layer into a 2D shape, then designs a toolpath to trace the perimeter of the shape. The number of times it traces this shape is determined by the wall line count setting. In the figure above, the wall line count setting is “1”. The algorithm then fills in the interior of the shape by drawing parallel lines to fill it in.

In order to generate a toolpath for a vascular channel, the user must set the wall line count to zero and the infill to 100%. If the width of a single extruded line is smaller than the channel the slicer is trying to print, the channel will be printed with an infill pattern composed of parallel striated lines. If the width is equal to the width of the channel, though, this will produce a toolpath which draws a single thread along the desired path. By drawing this thread between two towers, the print should have unrestricted flow at each end of the channel.

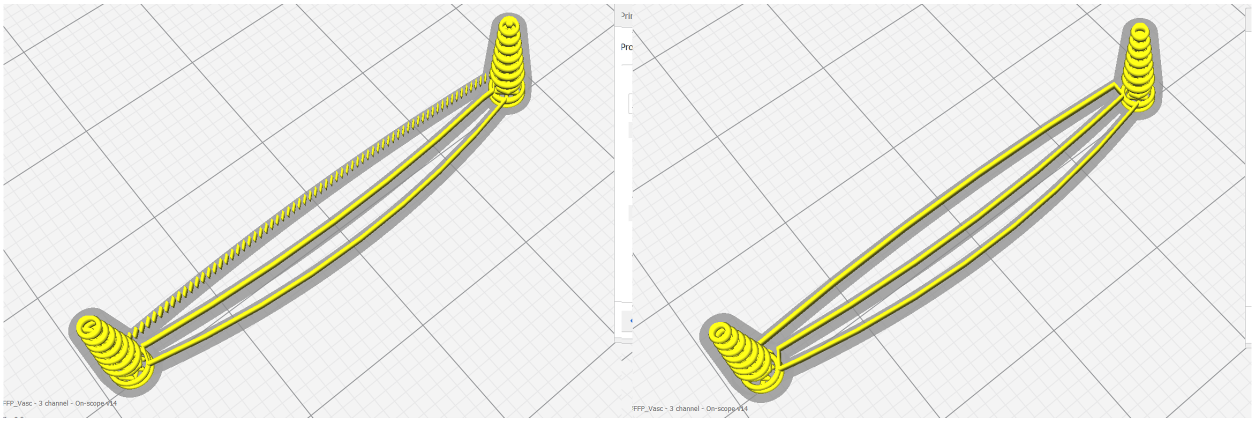

*Supp. FIg. 3.7: On the left, the infill pattern prints the channel as a series of broken parallel lines. On the right, the infill pattern draws an unbroken channel from one end to the other. This is a printable toolpath.*

#### 2.3 Optimizing the model for printing

In order to draw these channels, the original CAD model should be designed with dimensions such as the layer-height setting of the print in mind. During slicing, the user is trying to coax the slicing algorithm into predictably translate channels into a desired toolpath, so it makes sense if the CAD model is designed with planar features spaced the same distance apart as the layer height settings. In the examples provided, the channels are 1 mm tall and sliced with a 1 mm layer height. It should be noted, though that the final channel diameter doesn’t actually need to literally correspond exactly to the value of the model, as a channel’s width can be regulated by increasing or decreasing the amount of material extruded while the nozzle moves along the toolpath. Flow during printing can be increased or decreased using the Flow setting in Cura, which simply scales flow relative to 100% of the default. In fact, a user may find that in practice, a flow of anywhere from 50% to 300% may provide the ideal channel dimensions. As a result, the same model can be sliced to generate toolpaths for several prints with varied channel diameters based on the extrusion flow rate setting.

These vascular channels can also be stacked to vascularize a volume of indeterminate height. The channels can't be printed from one end to the other in air, as over this distance (35 mm in this design) gravity will pull them down. When printing sacrificial pluronic F-127 in air, a channel can typically be extended approximately 1 cm, however. This allows for multi-level printing of multiple channels provided they contact a support point every 10 mm or so. This support must be designed into the file, however. The algorithm which generates support in Cura isn't designed for this operation. Designing in support isn't difficult though: the support is just an extruded channel that crosses on top of the channels at regularly spaced intervals.

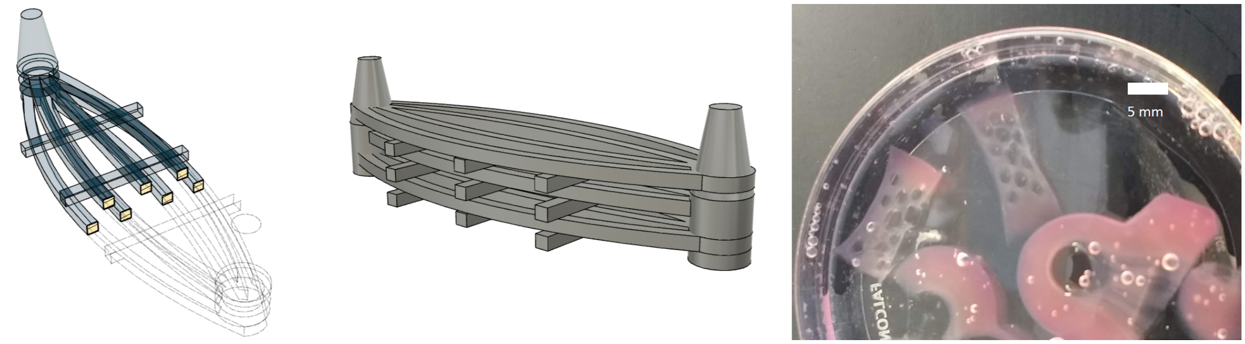

*Supp. Fig. 3.8: A multi-channel construct with three suspended layers is achievable if support material is manually designed into the model to limit the bridging distance an extruded filament needs to traverse to around 1 cm or less.*

*Table S3.1: Recommended Slicing Settings*

| Line | Length |
|---|---|
| Print Speed | 1 mm/s |
| Layer Height | 1 mm |
| Nozzle Width | 1 mm |
| Flow | 300% |
| Retraction enabled | Yes |
| Retraction distance | 0.2 |
| Prime after retraction | 0.01 |
| Z hop on retraction | Yes |

A construct with ample flow with abundant nutrient and oxygen flux into the matrix could also be much more easily cast around a loose mesh of sacrificial ink, however this paper is focused on producing channels for the purpose of replicating a liquid-matrix interface lined by endothelial cells.

#### 2.4 Refining the print conditions

Achieving precise placement of a scaffold on a slide can be done by correcting any minor mismatch between the x-y positions within the gcode and the actual desired placement using offsets. First, mark reference points, such as the center of where the towers should sit on a slide for calibration purposes and run a print. Measure the distance in the x and y directions by which the print was off of its target placement. This error can be corrected by setting x and y offsets to the printhead on the printer using the m851 command, or by setting x and y offsets to the home position using the m206 command. This can be manually entered into gcode files using a text editor, or sent as a command in Repetier-Host manually or within programmed button macros. Sending either of these commands to the printer alone (without any prescribed change) will instruct the printer to report back the current commands. These changes can be written into the printer's memory to be retained even after powering off by sending the command M500.

```
m851 x<x-value> y<y-value>
```

or

```
m206 x<x-value> y<y-value>
```

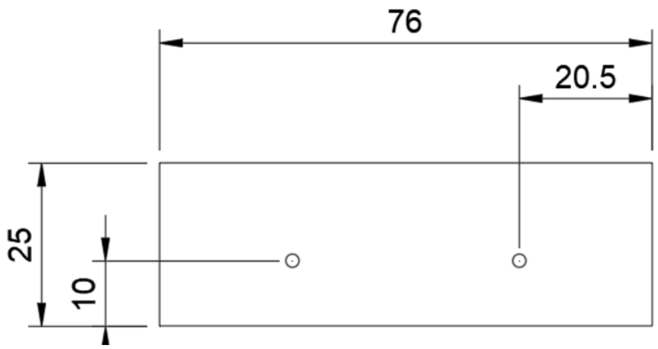

*Supp. Fig. 3.9: A reference image like this can be used to create a calibration slide.*

### IV. Preparing Bioinks, Printing, and Casting Constructs

**0. Preparing to print 37**
**1. Preparing bioinks 38**
1.1. Preparing 30% Pluronic F-127 38
1.2. Preparing Fibrinogen Bioink components in advance 40
1.3. Preparing complete fibrinogen-gelatin bioink on the day of use 41
**2. Printing a scaffold 43**
**3. Casting fibrinogen-gelatin ink around a sacrificial scaffold 44**
3.1. Prepare the housing to receive the FG ink 44
3.2. Add thrombin to initiate crosslinking and dispense quickly 45
**4. Flushing out the sacrificial scaffold 46**

#### 0. Preparing to print

Printing and casting bioprinted constructs can be a reliable, consistent process if all reagents and hardware are prepared in advance. The following checklists should be referred to in order to avoid discovering any item is not at hand when it's needed. It is recommended that items that will be used at the same time be collected together. For instance, housing components -- the base, gasket, capped lid, and two capped socket-socket adapters -- can be bagged and autoclaved together in the same sterilization bag.

| **Table S4.1: Printing Checklist** | | |
|---|---|---|
| | 18G dispensing/printing tips | Autoclaved |
| | Glass slides | Autoclaved |
| | Calibration slide | Autoclaved |
| | Transport box | Autoclaved |
| | 30% Pluronic F-127 in 5 mL printing syringe | |

| Table S4.2: Casting Checklist | | |
|---|---|---|
| | Bottom housing | Autoclaved |
| | PDMS silicone gasket | Autoclaved |
| | Upper housing (Capped) | Autoclaved |
| | Silicone oil in 1 mL syringe | Autoclaved |
| | 24G dispensing tips for applying silicone oil | Autoclaved |
| | 4x M5 screws | |
| | 4x M5 wingnuts | |
| | Thrombin | |
| | **OPTIONAL:** | |
| | A spacer | |
| | 30% Pluronic F-127 in 1 mL syringe with capped female-female adapter | |
| | Cells and cell media | |

| Table S4.3: Evacuating Scaffold Checklist | | |
|---|---|---|
| | Socket-socket Luer-Lok adapters, capped on both ends | Autoclaved |
| | 3 mL or 5 mL syringes | |
| | Chilled PBS in 10 mL wash syringe with female-female adapter and male cap | |

### 1. Preparing bioinks

The two bioinks used in this protocol were 30% Pluronic F-127 and 10% fibrin ink in gelatin. These provided the sacrificial scaffold and the bulk matrix respectively.

#### 1.1. Preparing 30% Pluronic F-127

Pluronic F-127 ink was prepared by dissolving 7.5 g of powdered Pluronic F-127 in a final volume of 25 mL of PBS supplemented with 2% pen-strep antibiotic/antimycotic (PSA). Because it is close to the saturation point of a pluronic solution, and because the solution is a solid gel at room temperature, mixing pluronic gel is very difficult. The process requires that the powder and its solvent be mixed by alternating between adding one and the other. It is recommended to consider making two 25 mL preparations of 30% pluronic ink at a time.

a. Weigh out 7.5 g of powdered Pluronic F-127.
b. Prepare 25 mL of PBS with 2% PSA (500 µL).
c. In a 50 mL conical tube, add approximately 1 mL of PBS containing 2% PSA. Then, sprinkle in just enough powder until there is dry powder sitting at the bottom of the tube and on the sides.

d. Add PBS again until all powder is wet. This will vary based on the amount of powder added in the previous step, but 1 mL is a reasonable expectation. If the added PBS is not clearly saturating all the powder, add less powder between additions of PBS.
e. Repeat the above steps until all of the powder has been added to the tube, and all of it is wet. The final volume should be very close to but slightly less than 25 mL.
f. Chill the wet pluronic down to at least 4°C. The Pluronic should shift in appearance from white to translucent. Storage at sub zero temperatures can be used to bring the temperature down faster, as freezing the gel will not hinder its preparation, but the gel will need to be above freezing to fully mix.
g. Briefly spin down the tube and add PBS to bring the final volume up to 25 mL. If done correctly, the pluronic should be completely or nearly completely dissolved, but the presence of lumpy, undissolved clumps is common.
h. Shake the tube by hand and then vortex if necessary to break up clumps or detach them from walls.
i. Place in a rotator at 4°C overnight.
j. The next day, the pluronic should have dissolved completely into a clear, viscous, fluid. If it is not fully mixed, make sure it is cold and repeat the above mixing, spinning, and over-night rotation steps until the solution is homogeneous.

Once a batch has been prepared, transfer the bioink into a large syringe in order to aliquot it into 5 mL print syringes or store for later.

k. Attach a sterile socket-socket Luer-Lok adapter to a sterile 20 mL syringe and gently load the syringe with 30% pluronic ink.
l. Remove the plunger in several 5 mL syringes and then transfer the rubber cap on the plunger to a 3D printed plunger-side coupler outlined in Supplementary Methods, Part 2, with a stylus or sterilized box cutter blade and spray down the cap with ethanol and allow to dry. Then, using a manual syringe loader (as described in Supp. 2), insert the custom plunger fully.
m. Using the socket-socket Luer-Lok adapter, mate the large syringe full of ink with each of the 5 mL printing cartridge syringes and draw back the manual plungers to fill each printing cartridge. The set volume is the volume after the plug-end of the plunger is fully exposed, and the base of the plug end is aligned with the back end of the syringe. The lengths of the custom plungers determine their volume when filled.
n. Cap each cartridge syringe with a socket Luer-Lok cap. These can be refrigerated or stored at room temperature.

#### 1.2. Preparing Fibrinogen Bioink components in advance

Fibrinogen bioink is composed of fibrinogen as the crosslinking monomer along with gelatin, which primarily adjusts the viscosity and thermostability of the ink. In order to prepare the ink, the constitutive parts must be prepared in advance.

**Selecting the right concentrations of Fibrinogen and Gelatin**

The concentration of fibrin in the ink affects the resulting firmness after crosslinking. Concentrations in literature range from 10 mg/mL (Kolesky *et al.* 2016, Homan *et al.* 2016) to 30 mg/mL (Kang *et al.* 2016, Skardal *et al.* 2017). Higher concentration will result in a firmer matrix, but 10 mg/mL appeared to be sufficient and more economical. Clarity is also a consideration. Clarity is achievable at higher concentrations, but it was easier to attain at 10 mg/mL.

Gelatin concentrations in literature range from 30 mg/mL (Wang *et al.* 2018, Skardal *et al.* 2017) to 75 mg/mL (Kolesky *et al.* 2016, Homan *et al.* 2016). The primary effect of gelatin concentration is its control over viscosity at different temperatures. Gelatin can be used to induce temperature-dependent phase transitions: at higher concentrations it can be used to print solid structures by warming it above its melting point and chilling it or letting it passively cool to room temperature after extrusion. Because it was being used in casting, printablity was not a concern. In this case, gelatin's primary role was to increase viscosity slightly, as a thinner solution lacking in gelatin underwent rapid, highly localized crosslinking upon the introduction of Thrombin. The presence of gelatin slowed the crosslinking enough to allow the solution to be mixed and deposited before it became unworkable. A concentration of 25 mg/mL was also low enough to allow filtration and to allow that the solution wouldn't solidify during the room-temperature incubation after adding transglutaminase which was necessary to produce a clear ink after crosslinking.

Kolesky, D. B. *et al.* (2016) 'Three-dimensional bioprinting of thick vascularized tissues.', *Proceedings of the National Academy of Sciences*, 113(12), pp. 3179. doi: 10.1073/pnas.1521342113.

Homan, K. A. *et al.* (2016) 'Bioprinting of 3D Convoluted Renal Proximal Tubules on Perfusable Chips', *Scientific Reports*. Nature Publishing Group, 6, pp. 1. doi: 10.1038/srep34845.

Kang, H.-W. *et al.* (2016) 'A 3D bioprinting system to produce human-scale tissue constructs with structural integrity', *Nature Biotechnology*. Nature Publishing Group, 34(3), pp. 312. doi: 10.1038/nbt.3413.

Skardal, A. *et al.* (2017) 'Multi-tissue interactions in an integrated three-tissue organ-on-a- chip platform', *Scientific Reports*. Nature Publishing Group, pp. 1–16. doi: 10.1038/s41598-017-08879-x.

Wang, Z. *et al.* (2018) 'Acta Biomaterialia 3D bioprinted functional and contractile cardiac tissue constructs', *Acta Biomaterialia*. Acta Materialia Inc., 70, pp. 48–56. doi: 10.1016/j.actbio.2018.02.007.

**Powdered Fibrinogen (60 mg)**

a. Weight out 60 mg of fibrinogen and add to a 50 mL conical tube, which will be used to produce a final volume of 6 mL of Fibrinogen-Gelatin ink containing 10 mg/mL of fibrinogen.

    Attempt to break large pieces apart if possible to improve dissolution. Use a 50 mL conical tube to enable filtration with a steriflip tube later.

    60 mg of fibrinogen will prepare a final volume of 6 mL of fibrinogen-gelatin bioink with 10 mg/mL of fibrinogen. If using another final volume (or concentration), adjust accordingly.

**100 mg/mL Gelatin solution (40 mL)**

Dissolve 4 g of gelatin in 40 mL of sterile PBS in a 100 mL bottle. Add a stir bar and autoclave with the cap sealed tightly. If the cap is loose, the solution will bubble and seep out of the bottle creating a big mess, so it is recommended to place this bottle inside a beaker with some water at the bottom and tighten the cap on the media bottle. Place on a stir plate to fully mix after autoclaving if necessary. Each construct only requires 750 µL of 100 mg/mL gelatin, so it is recommended that gelatin be stored in 10 mL volumes in 50 mL conical tubes at 4°C to allow for faster melting of the gelatin when preparing complete Fibrin-gelatin ink.

**250 mM $CaCl_2$ stock solution (10 mL)**

a. Dissolve 277.5 mg of $CaCl_2$ in 10 mL PBS (w/o Ca-Mg)

b. Sterile filter with a steriflip filter or a syringe-attached filter.

c. Aliquot in some multiple of 60 µL and freeze.

**50 mg/mL Transglutaminase stock solution (20 mL)**

a. Weigh out 1 g of Transglutamase (MooGloo)

b. Add the TG to 19 mL of PBS (w/o Ca/Mg) and bring to a final volume of 20 mL

c. Filter with a steriflip filter or by pushing the solution through a 0.22 µm Luer-Lok filter with a syringe.

d. Aliquot in some multiple of 300 µL volumes and freeze.

**50 U/mL Thrombin solution (20 mL)**

a. Dissolve 1 kU of Thrombin (Sigma T4648-1KU) in 20 mL of sterile-filtered PBS with 0.1% BSA to produce 20 mL of a solution of 50 U/mL.

b. Aliquot in multiples of 380 uL

    Crosslinking requires 60 µL of this solution per 1 mL of FG ink, so a standard 6 mL preparation of FG ink requires 360 µL of 50 U/mL thrombin (plus pipetting loss)

**20 mg/mL Aprotinin solution (2.5 mL)**

c. Dissolve 50 mg of Aprotinin (Abcam ab146286) in 2.5 mL of sterile-filtered PBS with 0.1% BSA to produce 2.5 mL of a solution of 20 mg/mL.

d. Aliquot in multiples of 40 µL.

Aprotinin is supplemented into media to produce a final concentration of 20 µg/mL, so 20 mg/mL can be added at 1 uL/mL of medium. Aliquoting 60x aliquots of 40 uL provides sufficient volume for 40 mL of medium per aliquot.

##### 1.3. Preparing complete fibrinogen-gelatin bioink on the day of use

FIbrinogen in solution will autocatalize relatively quickly, which is why its components are mixed the day the ink will be crosslinked.

a. Melt 100 mg/mL of gelatin at 37°C in a water bath, bead bath, or incubator.

b. Thaw aliquots of transglutaminase, thrombin, and aprotinin from use.

c. Add 4.5 mL of PBS or a base medium to 60 mg of fibrinogen powder.

d. Add 120 µL of 100x PSA

e. 60 µL of 250 mM $CaCl_2$.

f. Invert or vortex at a moderate speed to mix well while avoiding the formation of bubbles or applying unnecessary agitation and warm the solution for several minutes to 37°C.

g. Add 1.5 mL of 100 mg/mL gelatin to the fibrinogen solution.

h. Add 360 µL of 50 mg/mL transglutaminase.

i. Mix well by inversion, trituration, or moderate vortexing. Spin briefly to recollect the ink at the bottom of the conical tube and draw up bubbles if necessary.

j. Filter using a steriflip filter.

k. Incubate this solution at room temperature for at least 30 minutes and no more than 90 minutes.

**Incubating fibrinogen-gelatin ink after adding transglutaminase**

Transglutaminase is a secondary crosslinker that is necessary for the fibrinogen to maintain its structure after primary crosslinking occurs. Incubation at room temperature for greater than 30 minutes is necessary for the crosslinked ink to maintain its clarity, which is highly desirable for assessing the cells during the experiment and imaging the construct afterwards.

When the solution is left at room temperature for longer than 90 minutes, however, two problems may occur. First, the gelatin may congeal. The ink then loses its fluidity and cannot be effectively mixed with thrombin and dispensed in order to cast the matrix around the sacrificial scaffold. This can be corrected by gently rewarming the ink, however then the ink must be allowed to cool again, as thrombin added to warm ink will crosslink much faster than when added to room-temperature ink, and the working time may become too short to cast the matrix within.

Secondarily, transglutaminase can slowly initiate premature crosslinking if the ink is left too long at room temperature. This usually takes at least two hours, but the process is somewhat stochastic. If an experiment must be postponed after adding transglutaminase, the ink should be immediately moved to 4°C and then rewarmed the next day to melt the gelatin and then allowed to equilibrate to room temperature. This is not ideal handling, but can be used to avoid wasting complete FG ink, particularly if the goal is for testing some other part of the culture system or training, or otherwise not for the purpose of obtaining data.

Complete fibrinogen-gelatin ink has been stored at 4°C for up to a week before use successfully, but consistent use of fresh ink provided improvements to the consistency of the working time and optical clarity of the ink, which is why the use of fresh ink is strongly recommended.

#### 2. Printing a scaffold

a. Open Repetier-Host and connect to the printer.

b. Load the gcode print file.

c. Press button 1 to home the printer and move the printhead to its first position.

d. Insert a 5 mL syringe cartridge into the print head, uncap it, and attach a plastic dispensing needle.

e. Press button 1 to home the printer again, as snapping the cartridge into place can unhome the printer.

f. Place a calibration slide onto the stage adapter bracket.

g. Press button 2 to move the needle tip into position two, just above the reference point marked on the slide.

h. Use the manual arrow controls to position the needle in the x, y, and/or z axes if needed. When the needle is positioned correctly, press button 3 to assign the current position as the reference position. The needle will then move to position three, off to the side of the slide.

i. If a consistent correction is needed each time, consider modifying the M206 command in button 1's macro.

j. Remove the calibration slide and replace it with a fresh, clean, sterile microscope slide (or other print surface).

k. Press button five repeatedly to prime the nozzle. Pluronic ink should be visible advancing within the dispensing tip until it emerges. Once it has emerged, wait a few seconds to make sure it has no further residual motion and then wipe the extruded ink away using a p10 tip or other sterile tool. Wait several more seconds to confirm that no further oozing is occurring.

l. Press "Print" to begin the print process.

m. Once the print completes, move the slide into a small lidded transport box. If necessary, repeat the process to produce additional scaffolds as needed.

n. When finished printing, the tip can be removed and the cartridge recapped. Or, the tip itself can be capped with a socket-socket Luer-Lok adapter with a cap on one end. This can be wedged snuggly onto the tip, and the next print can be run using the contents of the current pluronic ink cartridge.

### 3. Casting fibrinogen-gelatin ink around a sacrificial scaffold

Casting matrix around a scaffold is not difficult, but it is a sensitive process that must be executed with fluid, confident movement in order to work. Carefully placing everything where it should be before adding the thrombin crosslinking agent will allow for the consistent preparation of high-quality 3D printed constructs.

Instructions are provided both for filling a housing completely or only partially. Filling a housing completely limits the void space exclusively to the channel, which can improve seeding, continuous feeding, and assaying the construct. However doing so also reduces the volume in which medium can reside, which may prevent manual feeding to be performed in place of continuous feeding. For process development purposes, manual feeding is recommended when attempting to isolate the influence of separate system components, which is why instructions for partial filling and manual feeding are included (and recommended for initial system tests).

#### 3.1. Prepare the housing to receive the FG ink

a. Before casting the ink, open the autoclave bag containing the housing components and lay them out in convenient reach.

b. Place the microscope slides or other print surface into the base of the housing.

c. Apply a thin, continuous bead of silicone oil around the perimeter of each side of the gasket.

d. Gently lay the gasket into place on the print surface taking care not to disturb the scaffold.

e. Place the lid on top of the base and tighten it down with wingnuts.

f. Uncap a filling port and a venting port and attach a socket-socket adapter to each filling port which is to be filled. In the microscope compatible design, the filling port is between the two flow ports, and either of the flow ports can act as the vent port.

g. IF FILLING THE HOUSING COMPLETELY: Lay a serological pipet in the hood and set the housing on top of this pipet so that it lays at a gently sloping angle, so that the capped flow port is lower than the venting port.

   IF FILLING THE HOUSING PARTIALLY: Uncap a vent port and leave the housing sitting flat.

h. Place a 24-well cell culture plate to the side of the housing without its lid.

At this point, the housing should be assembled with scaffolds sitting on glass slides (or other print surface) within the gasket. The gasket should have a visible seal against the lid and be ready to receive ink.

**If cells are to be included in the FG ink, this is the point to mix them in.**

##### 3.2. Add thrombin to initiate crosslinking and dispense quickly

a. Draw 360 μL of room temperature 50 U/mL thrombin into a p1000 pipet tip.

b. Unwrap a 10 mL serological pipet and insert it into a pipetter set to a slow draw and speed and a medium dispense speed.

c. Using a non-dominant hand, dispense the thrombin into the room temperature FG ink.

d. Draw up the ink and dispense it twice, while gently stirring before drawing up the full volume of ink.

e. Place the tip of the serological pipet firmly in the opening of the socket-socket Luer-Lok adapter in the fill port and begin to dispense.

f. <u>IF FILLING THE HOUSING COMPLETELY</u>: Watch the ink fill the housing from the lower side up to the other side and stop before the ink reaches the vent port.

   <u>IF FILLING THE HOUSING PARTIALLY</u>: Watch the ink fill the housing and stop filling when the housing is filled to the chosen level.

g. Quickly repeat for the second construct in the housing.

h. Dispense any remaining ink into wells of the waiting 24-well plate.

i. Cap the vent ports and lay the housing flat. Cap the fill ports and replace the lid on the 24-well plate.

j. Move the housing and 24-well plate into a 37ºC incubator.

At this point, the housings should be sealed and filled with an optically clear matrix and the scaffolds should be visible and undisturbed within them.

#### 4. Flushing out the sacrificial scaffold

In order to flush out the scaffolds, the matrix should be incubated at 37°C to allow for thorough crosslinking and then chilled. While initial experiments were attempted with cell-laden inks, focus was shifted towards cell-free inks in order to enable refinement of the casting and seeding process without the complicating influence of cells on the timing and handling requirements.

a. Incubate the housing at 37°C for 30 minutes.

b. Remove the housing from the incubator and allow it to cool for 10 minutes at room temperature.

c. Place the housing into a -20°C freezer for 20 minutes.

d. Examine the housing. The housing should be chilled sufficiently that the pluronic liquifies, but not so long as to allow ice crystals to form within the matrix. If the pluronic exhibits no flow, return the housing to the freezer for another 2-5 minutes.

e. Once some pluronic flow is observed, uncap both flow ports of a single construct within a cell culture hood and place the tip of a pasteur pipet connected to a vacuum aspiration line at the opening of one end and tilt the housing. Insert the aspirator tip into the port and attempt to aspirate as much pluronic ink as possible without disturbing the matrix.

f. Repeat for any additional constructs.

g. Remove the plungers from a pair of 3 mL or 5 mL syringes and attach them to both flow ports of a construct using socket-socket Luer-Lok adapters.

h. Dispense chilled PBS into one syringe, and insert an aspirating pasteur pipet into the port on the other side to aspirate the PBS out the opposing end.

i. After thoroughly flushing the channels, fill them with cell culture medium and either recap them until ready to seed cells into the channels or connect to active flow, as described in Supplementary Methods, Part 5.

**Flushing Pluronic F-127 fully is challenging**

Flushing channels of Pluronic F-127 is essential to successful cell seeding, however it requires a difficult balance. Pluronic ink is a stubborn material that both resisists washing and inhibits cell attachment. Determined, persistent flushing is required in order to remove all residue and present cells with a fibrinogen surface receptive to cell attachment, however the same forces which wash out this residue can also damage the fibrinogen substrate.

Ultimately, the best approach is to wash gently and repeatedly, with lots of chilled PBS over many minutes, exercising patience. Repeated washing on distinct days and continuous flow within an incubator can also facilitate flushing.

### V. Generating iECs and Seeding Constructs

**1. Generating iPSC-derived Endothelial Cells 47**
1.1 Culturing iPSCs 48
1.2 Differentiating Cells 48
**2. Seeding iECs into channels 48**
2.1 Determining the volume and surface area of a construct's void spaces 49
2.2 Seeding cells into a construct 49
2.3. Assessing seeded cells and feeding 50
Bioprint Construct Assessments 52

#### 1. Generating iPSC-derived Endothelial Cells

Endothelial Cells were differentiated from iPSCs based on the protocol described by Harding:

Harding A, et al. (2017) Highly Efficient Differentiation of Endothelial Cells from Pluripotent Stem Cells Requires the MAPK and the PI3K Pathways.*Stem Cells* 35(4):909.DOI: 10.1002/stem.2577

*Supp. Table 5.1: Reagents*

| Reagent | [Stock] | Supplier | Cat. # | Description |
|---|---|---|---|---|
| Matrigel | 10 mg/mL | Corning | 354263 | Basement membrane |
| mTeSR + | - | STEMCELL Technologies | 100-0276 | Stem cell medium |
| StemDiff APEL | - | STEMCELL Technologies | 05275 | Base differentiation medium |
| CHIR99012 | 10 mM | Xcess Biosciences | m60002 | Wnt signaling activator |
| EC Growth Medium MV2 | - | PromoCell | C-22022 | iEC base medium |
| BMP-4 human protein | 50 µg/ml | R&D Systems | 314-BP-010 | Vascular progenitor activator |
| VEFG Human protein | 100 µg/ml | R&D Systems | 293-VE-010 | Vascular progenitor activator |
| FGF-2 | 100 µg/ml | PeproTech | 100-18B | |
| ROCK inhibitor Y-27632 | 10 mM | Cayman | 10005583 | Apoptotic inhibitor |
| Accutase | 1x | Stem Cell Technologies | 07922 | Enzymatic dissociation reagent |

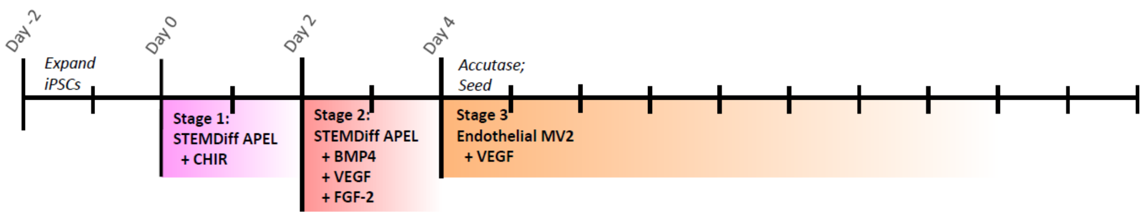

*Supp. Fig. 5.1: Schematic timeline of iEC differentiation*

#### 1.1 Culturing iPSCs

Pluripotent Stem Cells were maintained on matrigel coated plates in mTeSR+ medium and passed with EZ-Pass cell passage tools. Before beginning a differentiation, cells were plated onto matrigel-coated 100 mm dishes at approximately 5% density in order to produce medium-sized colonies with ample space between them.

#### 1.2 Differentiating Cells

a) On Day 0, administer Stage 1 iEC medium to begin mesoderm induction.

*Supp. Table 5.2: Stage 1 iEC Medium – Mesoderm Induction Medium (Day 0)*

| Reagent | [Stock] | [Final] | 10 mL |
|---|---|---|---|
| StemDiff APEL | - | - | 10 mL |
| CHIR99021 | 10 mM | 6 μM | 6 μL |

b) Do nothing on Day 1.
c) On Day 2, feed with Stage 2 iEC medium.

*Supp. Table 5.3: Stage 2 iEC Medium – Vascular Progenitor Medium (Day 2)*

| Reagent | [Stock] | [Final] | 10 mL |
|---|---|---|---|
| StemDiff APEL | - | - | 10 mL |
| BMP4 | 50 μg/mL | 25 ng/mL | 5 μL |
| VEGF | 100 μg/mL | 50 ng/mL | 5 μL |
| FGF2 (FGF-basic) | 100 μg/mL | 10 ng/mL | 1 μL |

d) On Day 4, dissociate with accutase and replate onto matrigel-coated 100 mm dishes with 10 uM ROCK inhibitor on the first day.

*Supp. Table 5.4: Stage 3 iEC Medium – Endothelial Progenitor Medium (Day 4 onward)*

| Reagent | [Stock] | [Final] | 10 mL |
|---|---|---|---|
| StemDiff APEL | - | - | 10 mL |
| VEGF | 100 μg/mL | 50 ng/mL | 5 μL |

### 2. Seeding iECs into channels

In order to seed iECs into channels, the iECs should be dissociated and suspended in endothelial medium with 10 uM ROCK inhibitor Y-27632 at a volume that will occupy the channel void spaces and with sufficient cells to cover available surface area at a density of at least 5,000 cells/cm2.

#### 2.1 Determining the volume and surface area of a construct's void spaces

The volume of medium cells are suspended into should be approximately equivalent to the volume of the space available within the construct. This volume and surface area can be estimated by calculating the volume of the void space in whatever CAD program was used to create the model of the scaffold.  For this project, a single channel was was used, which received a volume of 200 uL, in which I suspended 1 E6 cells.

*Supp. Table 5.1: Surface area estimations for various scaffolds*

| Channels | Volume [$mm^3$] | Volume [mL] | Surface Area [$mm^2$] | Surface Area [$cm^2$] |
|---|---|---|---|---|
| 1 Channel | 198 $mm^3$ | 0.198 mL | 402 $mm^2$ | 4 $cm^2$ |
| 3 Channels | 288 $mm^3$ | 0.288 mL | 707 $mm^2$ | 7 $cm^2$ |
| 6 Channels | 411 $mm^3$ | 0.411 mL | 1152 $mm^2$ | 11.5 $cm^2$ |
| 9 Channels | 536 $mm^3$ | 0.536 mL | 1603 $mm^2$ | 16 $cm^2$ |

Both higher and lower cell densities were tested, and lower cell densities (5 E5, for instance) were observed to produce fewer attached cells than 1E6, though still an ample density. Higher densities (such as 2E6) were not observed to increase the density of attached cells, and the higher metabolic consumption strain that more cells placed on a very small volume of medium was considered reason to include no more cells than were observed to be effective.

#### 2.2 Seeding cells into a construct

Seeding cells is one of the most challenging steps of this project, and it was here that the live-imaging feature of this system demonstrated its essential need. These cells attach very readily to a fibrin-gelatin substrate in a dish, but often exhibited lower attachment within a channel. Experimentally, it appeared that this was likely due to residual pluronic F127, as the channels received cells much better if washed aggressively and conditioned over several days at $37^o$ C.

In order to fully cover the interior of channels, cells were seeded, allowed to settle onto one face, and then repositioned. This was achieved initially by simply inverting a construct, however it was found that this left wide bald patches, as attachment did not occur effectively above a $45^o$ angle. To accommodate this, the housing was then rotated manually through six positions using a hexagonal bracket. Later, this was replaced with a custom rotator which would rotate a housing through a prescribed set of 8 positions on a prescribed schedule all through the night after seeding. This improved consistency, simplified the handling process on seeding days, and allowed for more positions to be placed without extending the time between seeding and refeeding or connecting to flow.

In both cases, though, a single seeding was not found to be sufficient to achieve the density of coverage desired, and three seedings over three successive days was found to be the most effective routine.

*Supp. Table 5.1: Example experiment calendar*

| Sunday | Monday | Tuesday | Wednesday | Thursday | Friday | Saturday |
|---|---|---|---|---|---|---|
| | | ExpandiPSCs | | Admin. Stg. 1<br>iEC medium | Mix ink,<br>Print,<br>Cast,<br>Flush | Admin. Stg. 2<br>iEC medium |
| | Flush again,<br>Accutase cells,<br>Seed,<br>Freeze rest | Observe,<br><br>Feed manually | Observe,<br><br>Thaw & reseed if necessary | Observe,<br><br>Thaw & reseed if necessary | Connect to flow | |
| | Observe,<br>Change med. | Observe,<br>Change med. | Observe,<br>Change med. | Observe,<br>Change med. | Assay | |

While a full channel could theoretically be fully covered in a single go, seeding half the channel on one day, then seeding the opposing half two days after was established as the standard procedure.

a. Print scaffold, cast ink, and flush.

b. Prepare a suspension of the appropriate number of iECs either from a culture dish or a frozen stock. For a single channel, 1 E6 cells was used for each channel.

c. Spin down the cells at 200 rcf for 3 minutes.

d. Aspirate and resuspend the cells in the appropriate volume of iEC medium with ROCK inhibitor. For a single channel, 200 µL was added.

e. Fully aspirate the contents of a channel using a pasteur pipet. The channel should be visibly empty of fluid.

f. Add the suspension into the channel using a micropipette. The cell suspension should fill the channel fully.

g. <u>IF USING A ROTATOR</u>: Place the housing in the rotator. and plug it in. It will cycle through eight positions over 30 seconds before stopping at position one.

   <u>IF NOT USING A ROTATOR</u>: Place the housing at the desired angle.

   i. For initial tests, simply leave the construct sitting flat overnight.

   ii. Later, consider placing the construct in the hexagonal bracket and rotating it to a new position after 45 minutes. If successful, add additional positions and adjust timing based on outcomes.

h. The next day, replace the medium with fresh iEC medium.

##### 2.3. Assessing seeded cells and feeding

Seeded cells were fed daily, either with a pipette or a continuous pump, and observed using an ECHO Revolve fluorescent microscope daily. If seeded on all sides, the cells would propagate and fill small gaps, after which point the construct could be stained or assayed. Major cell migration, however, was not observed.

Cell behavior within the construct was found to be fickle, and careful monitoring was essential to refine the system into something producing consistent seeding and stable cell health throughout an experiment. The following assessment guide was used to effectively characterize and troubleshoot cell seeding success and the behavior and morphology of cells during experiments

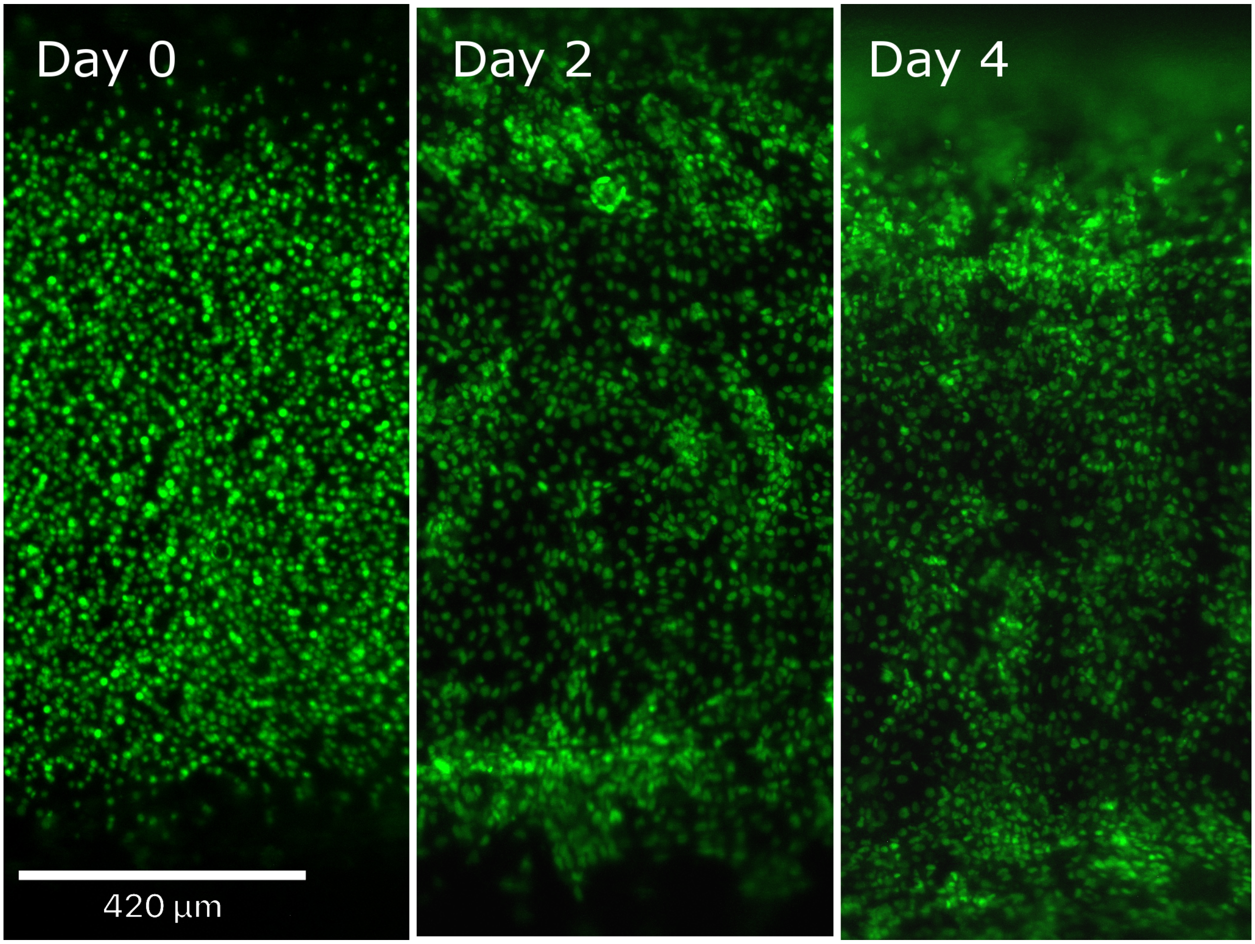

*Supp. Fig. 5.2: Images of GFP-expressing iECs seeded into a channel on the day of seeding, two days after seeding, and four days after seeding.*

### Bioprint Construct Assessments

Construct Preparation Date: ____________________
Ink Mix Time: ____________________
Print Time: ____________________
Casting Time: ____________________
Chill Time: ____________________
Flush Time: ____________________

| **Print Assessments** |
|---|
| Tower placements? |
| Channel lift? |

| **Casting Assessments** |
|---|
| Pre-crosslinking viscosity |
| Fill Level |
| Initial Seal |
| Clarity |
| Homogeneity |

| **Post-Incubation Assessments** |
|---|
| Seal |
| Clarity |
| Homogeneity |
| Channel Interface:<br>- Smoothness<br>- Color<br>- Juncture pits<br>- Description |

| **Post-Incubation** | **A** | **B** |
|---|---|---|
| Seal | | |
| Clarity | | |
| Homogeneity | | |
| Channel Interface:<br>- Smoothness<br>- Color<br>- Juncture pits<br>- Description | | |
| Other | | |

Seed time: ______________________

| Culturing | A | B |
|---|---|---|
| Day 1 | 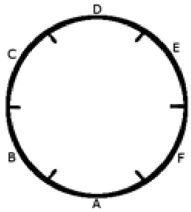 | 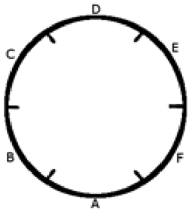 |
| | 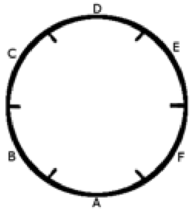 | 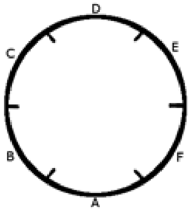 |
| | 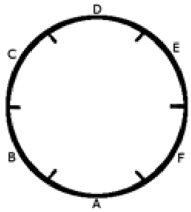 | 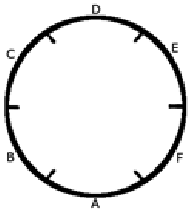 |
| | 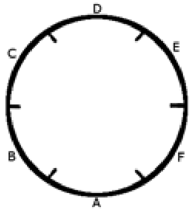 | 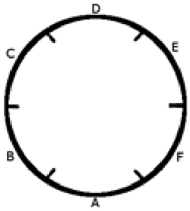 |
| | 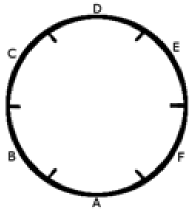 | 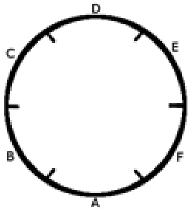 |
| | 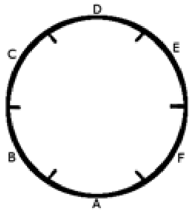 | 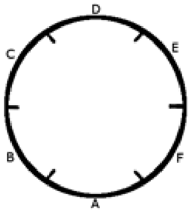 |

### VI. Culturing, Harvesting, and Imaging Constructs

**1. Maintaining Bioprinted Constructs in Culture 54**
1.1: Flow system setup 55
1.2. Feeding manually 56
**2. Live-Imaging Constructs 57**
**3. Harvesting 57**
3.1. Sectioning for staining 57
3.2. Digesting fibrin for RNA extraction 58
**4. Staining and Imaging Fixed Constructs 58**
4.1: Immunohistological staining solutions 58
4.2: Immunohistological staining procedure 59
4.3: Viability staining solutions 59
4.4: Viability staining procedure 60
**5. Confocal fluorescence imaging 60**

#### 1. Maintaining Bioprinted Constructs in Culture

Bioprinted constructs can be maintained with either continuous-flow feeding or manual batch feeding. Continuous-flow feeding offers a consistent internal environment and the ability to maintain a higher through-put of experiments by reducing hands-on time. After seeding cells into a construct, supply the construct with medium through continuous flow.

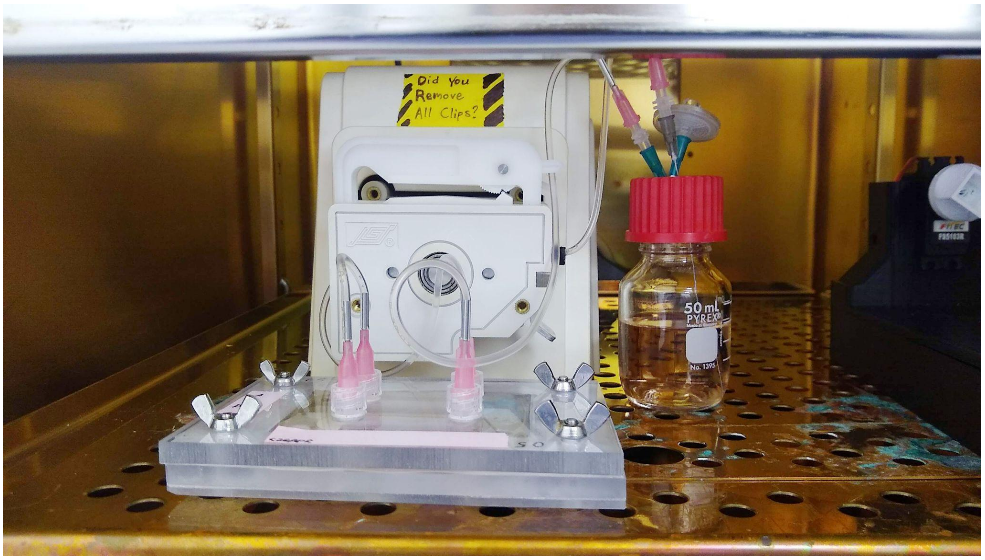

*Supp. Fig. 6.1: Cycled flow running from a 50 mL culture bottle resevoir with a septum cap, through both channels in series, back into the reservoir.*

#### 1.1: Flow system setup

a. To set up flow, first autoclave a sterilization bag containing the flow lines and their connectors, as well as the reservoir or reservoirs needed.

The parts list can be found in Supplement 1, table S1.3.

| **Table S6.1:Flow Setup Checklist** | | |
|---|---|---|
| | Reservoir(s) with septum caps | Autoclaved |
| | Flow tube bag: | Autoclaved |
| | - 2x - Male-to-male Luer coupler | Autoclaved |
| | - Inflow line with dispensing needle on each end | Autoclaved |
| | - Transfer line with dispensing needle on each end | Autoclaved |
| | - Outflow line with dispensing needle on each end | Autoclaved |
| | 4” long inflow needle | |
| | Outflow needle | |
| | Vent needle | |
| | Vent filter | |
| | 1 mL syringe | |
| | Binder clip | |

b. In a biosafety hood, fill the inflow reservoir with medium and cap it.

c. Spray down the septum and the capped ports of the construct housing with 70% isopropyl alcohol and allow them to dry.

d. Connect the 4” inflow needle to the inflow line using a male-to-male Luer coupler and push it through the septum.

e. Connect the outflow needle to the outflow line using a male-to-male coupler and push it through the septum as well.

f. Connect a filter to the vent needle and push it through the septum.

g. Connect the 1 mL syringe to the opposing end of the inflow line and draw medium in to fill the line. Clip it with a binder clip.

h. Uncap a port on the housing and connect the free end of the inflow line to it.

i. Uncap the other port of the same channel connect one end of the transfer line. Connect the other end of the transfer line to the 1 mL syringe.

j. Remove the binder clip from the inflow line and gently draw on the 1 mL syringe to pull medium into the housing and fill the transfer line. Replace the binder clip on the inflow line.

k. Uncap the port adjacent to the outflow of the first channel and connect the free end of the transfer line.

l. Uncap the outflow port of the second channel and connect the free end of the outflow line.

m. Disconnect the outflow end of the outflow line from the Luer coupler joining it to its needle and connect it to the 1 mL syringe. Draw to pull medium through the system and at least partially fill the outflow line.

n. Replace the binder clip anywhere along the flow lines and then reconnect the outflow end of the outflow line to the Luer coupler attached to the outflow needle.

o. Using the construct caddy, move the construct(s) and reservoir(s) to the incubator.

p. Unlatch the peristaltic line retaining clip and install the inflow line into the peristaltic pump, taking care not to pull on the connections at either end of the line.

q. Remove the binder clip arresting flow.

> This step is surprisingly easy to forget. REMOVE CLIPS BEFORE COMMENCING FLOW.

r. Turn on flow at the lowest speed and adjust upward until fluid movement is observable in the outflow line. Allow it to run until the outflow line is filled and begins dripping medium into the outflow reservoir (or single reservoir in a cycling system).

s. Adjust the flow speed down to the desired speed.

> It is advisable to check the flow system an hour later or at the end of the day before leaving the lab to confirm flow system operation.

##### 1.2. Feeding manually

Manual feeding removes complexity associated with the flow system which may be desirable to isolate seeding and other processes from development and testing of the flow system. The small volumes of medium involved may leave cells nutrient or oxygen limited, however, unless feeding is suitably frequent. Filling the housing only partially with fibrin ink creates a headspace above the construct that allows for a larger volume of medium to be held within the construct and is recommended if feeding manually.

a. Retrieve the construct housing from the incubator.

b. Spray down the housing ports with IPA and allow to dry.

c. Uncap the inlet and outlet of one channel.

d. Turn on the vacuum aspiration line half way and place the tip of a Pasteur pipet in the mouth of one port and gently tip the construct to drive flow towards the opening.

> Inserting the tip of the Pasteur pipet is fine, so long as the tip doesn't suck up the fibrin substrate. If this happens, the substrate is likely to tear or detach from the glass basement. The size of the ports and the clear housing make it fairly easy to insert the Pasteur pipet quite far without accidentally aspirating fibrin matrix.

e. Alternatively, medium can be removed by placing a p1000 tip into the port, tilting the construct, and drawing medium into the tip.

f. Replace the medium with a fresh 1000 uL of medium, or whatever volume fits using a similar approach to withdrawing the medium:

   - Avoid aggressive pipetting to limit unwanted sheer stress.
   - Tilt the housing as necessary to fill it.
   - Avoid overfilling. Any overflow from the alternate port presents contamination risk.

g. Recap the ports snugly. Excessive tightening is unnecessary and will wear out the ports over time.

h. Repeat for additional channels and return the housing to the incubator.

### 2. Live-Imaging Constructs

Constructs were live imaged daily on an ECHO Revolve fluorescent microscope.

### 3. Harvesting

#### 3.1. Sectioning for staining

Constructs can be removed from the housing for analyses similar to bulk tissue. They can be sectioned, as well as dissolved for RNA sequencing or RT-qPCR analysis. These approaches were developed while exploring the use of cell-laden bioprinted constructs. While these were determined to be outside the scope of the present study, the slicing jigs used are included in the printable files. After sectioning, place the sections in wells of a 24 well or 12 well plate. Deposit several drops of warm 1% agar in the dish in order to immobilize the construct sections.

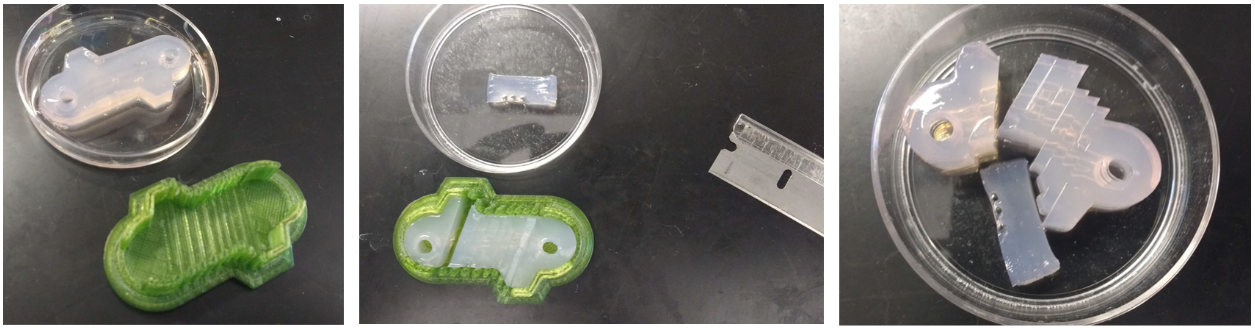

*Supp. Fig. 6.1: Sectioning a construct in a sectioning jig*

##### 3.2. Digesting fibrin for RNA extraction

While cell-impregnation was determined to fall outside the scope of this paper, a summer intern performed experiments on the use of proteinase K to digest fibrin-gelatin matrix in order to extract cells from within ink.

a. Prepare a solution of 32 U/mL of Proteinase K (New England Biolabs, P8107S) in PBS.

   For each sample, the recommended volume of 2 mL can be prepared by adding 80 μL of 800 U/mL proteinase to 1.92 mL of PBS. The use of trypsin in place of PBS was not observed to provide any benefit, although other enzyme solutions might.

b. In a 60 mm dish, dice the construct into the smallest possible pieces with a razor blade.

c. Transfer the diced pieces into a well of a 12-well plate using a spatula.

d. Add 1 mL of proteinase K digestion solution into the 60 mm dish to suspend any pieces of matrix too small to lift with a spatula and then transfer the solution into the well of the 12-well plate.

e. Add an additional mL to ensure sufficient enzyme to digest the fibrin-gelatin matrix.

f. Place the dish in an incubator for 30 minutes on a rocker or orbital shaker.

g. If the contents are suitably dissolved, transfer to a 2 mL tube, 15 mL tube, or two 1.5 mL tubes and spin to pellet.

### 4. Staining and Imaging Fixed Constructs

##### 4.1: Immunohistological staining solutions

**Blocking & Permeabilization Buffer (10 mL)**

9 mL PBS
1 mL Donkey Serum

15 uL Triton X100

**TWEEN Wash Buffer (100 mL)**
100 uL TWEEN in 100 mL PBS

**Hoescht Stain 33342, 20 mM (5 mL)**
2 uL Hoescht in 5 mL PBS (8 uM)

##### 4.2: Immunohistological staining procedure

Staining can be performed both on sections in a dish or on intact constructs within their housing. In either case, the staining procedure is the same.

a. Samples should be fixed for 20 minutes with 4% paraformaldehyde or other fixative.

b. After fixation, wash samples three times with PBS.

c. Fully cover the sample in blocking buffer and block for at least 1 hour, or overnight if staining cells within the matrix.

d. Prepare primary antibodies such as the following:

| Protein | Host | Manufacturer | Product Number | Dilution |
|---|---|---|---|---|
| CD31 | Mouse | Cell Signalling | 3528S | 1:50 |
| VEGF R2 | Rabbit | Cell Signalling | 2479S | 1:50 |

e. Administer 1$^{o}$ antibody solution and incubate at 4$^{o}$ C overnight.

f. Rinse with a wash buffer consisting of PBS + TWEEN three times for at least 5 minutes each, preferably on a rocker.

g. Prepare 2$^{o}$ antibody solution such as the following:

| Antibody | Target | Manufacturer | Product Number | Dilution |
|---|---|---|---|---|
| Alexafluor 568 | Mouse | Thermo Fisher | A10037 | 1:250 |
| Alexafluor 568 | Rabbit | Thermo Fisher | A10042 | 1:250 |
| Alexafluor 405 | Rabbit | Thermo Fisher | A48258 | 1:250 |

h. Apply the 2$^{o}$ antibody solution and stain overnight.

##### 4.3: Viability staining solutions

**Live/Dead/Hoechst Stain**

a. Prepare 50 uL of 1 mM stock Calcein solution by diluting 50 ug Calcein in 50 uL DMSO

b. To 2 mL of PBS, add:

   i. 6 uL of 1mg/mL Propidium Iodide
   ii. 6 uL of 1 mM Calcein
   iii. 2 uL of 20 mM Hoechst 33342
   iv.

##### 4.4: Viability staining procedure

Add Calcein/PI/Hoechst stain and incubate for 30 minutes to 1 hr in the dark, then rinse in PBS and image.

### 5. Confocal fluorescence imaging

The cells were imaged using a Nikon confocal microscope running Nikon's Elements software.

a. The construct was placed on the stage and moved into an appropriate x-y position.

b. Light was directed through the microscope's eyepiece, which was used to find the correct z-position of the sample of interest.

c. The light was directed back to the camera collector. Initial view settings were 512 px resolution, 1 frame per second, with a laser power of 2.

d. The sample was viewed by either turning on live viewing or taking a series of captures to identify the area of greatest interest in all relevant dimensions.

e. Brightness and any other settings were adjusted as needed for each fluorescent channel.

f. A test capture was performed with 1/10th as many Z-stack slices as intended in order to confirm that the sample is appropriately framed and that the Z-intensity correction settings are right.

   It is advisable to load the test capture into IMARIS to confirm the fluorescence signal at the top of the channel is visible in the reconstructed 3D volume.

g. Repeat test captures until the fluorescence is ideal, then set the Z-stack slice number to its full value and begin the capture.

h. Load the full capture into the Imaris high content imaging program.

### VII. Assessing Diffusional Barriers

**1. Prepare the flow system and reagents 62**
- 1.1: Prepare the FITC flow system 62
- 1.2: Prepare the FITC-Dextran solution 63

**2. Calibrate the microscope settings 63**
- 2.1: Frame the point of interest in their X-Y positions 64
- 2.2: Locate the ideal Z-position for capture 64

**3. Capture diffusion 64**
- 3.1: Preparing the channel 64
- 3.2: Flush channel and begin capture. 65

**4. Post-process image stack 66**

**5. Analyze and quantify the rate of diffusion 66**

**6. Compare the speed of diffusion between time serieses 67**

Visualizing dye movements within a printed construct provides a flexible template for leveraging bioprinting for meaningful research aims. Here, we describe how one can watch a fluorescent dye diffuse out of a channel in order to measure the permeability of a layer of cells lining a channel. In the following example we use a Nikon A1 confocal microscope with Nikon's NX Elements software to capture our images, FIJI to process them, and the R mathematics language to analyze them.

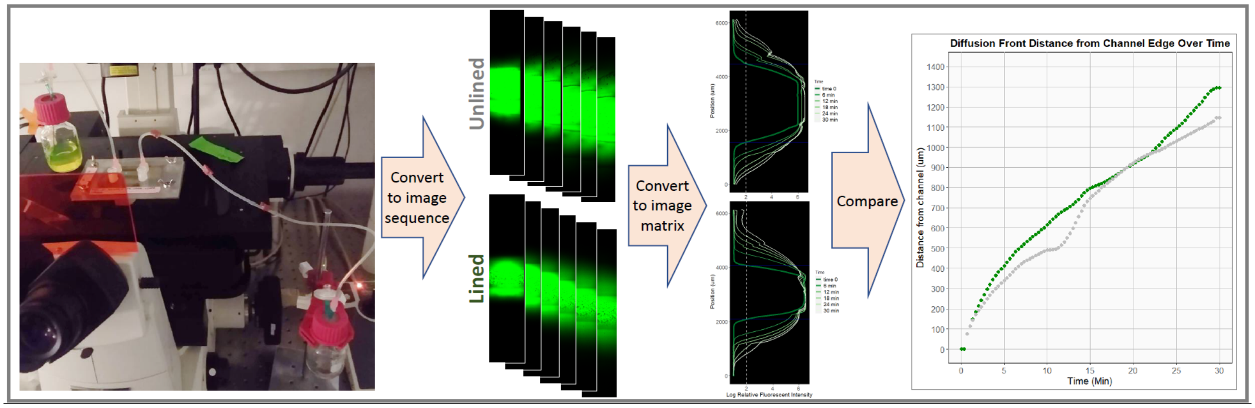

*Supp. Fig. 7.1: Schematic of the FITC diffusion assay pipeline*

### 1. Prepare the flow system and reagents

#### 1.1: Prepare the FITC flow system

The assay cannot be performed with static FITC dye inside the construct. The fluorescent intensity over the course of 30 minutes will decrease too much to observe outward diffusion, presumably due to a combination of photobleaching and because the decline in concentration within the channel reduces the concentration gradient that drives diffusion. In order to combat this, we need to have constant-rate flow through the channel to maintain the concentration gradient that drives FITC to diffuse outward. Because setting up a pump system in a microscope room is cumbersome, we instead connected a modified version of the flow system used to feed the constructs and created a gravity-driven flow system consisting of an inflow reservoir, an inflow line, an outflow line, and an outflow reservoir mounted on a test tube stand.

| **Table S7.1: Flow Setup Checklist** The parts list can be found in Supplement 1, table S1.3. | |
|---|---|
| | 2x - Reservoirs with septum caps |
| | Inflow line with dispensing needle on each end |
| | Outflow line with dispensing needle on each end |
| | 4” long inflow needle |
| | Outflow needle |
| | 2x - Male-to-male Luer coupler |
| | 2x - Vent needles |
| | 2x - Vent filter |
| | 1 mL syringe |
| | Binder clip |
| | Test tube stand |

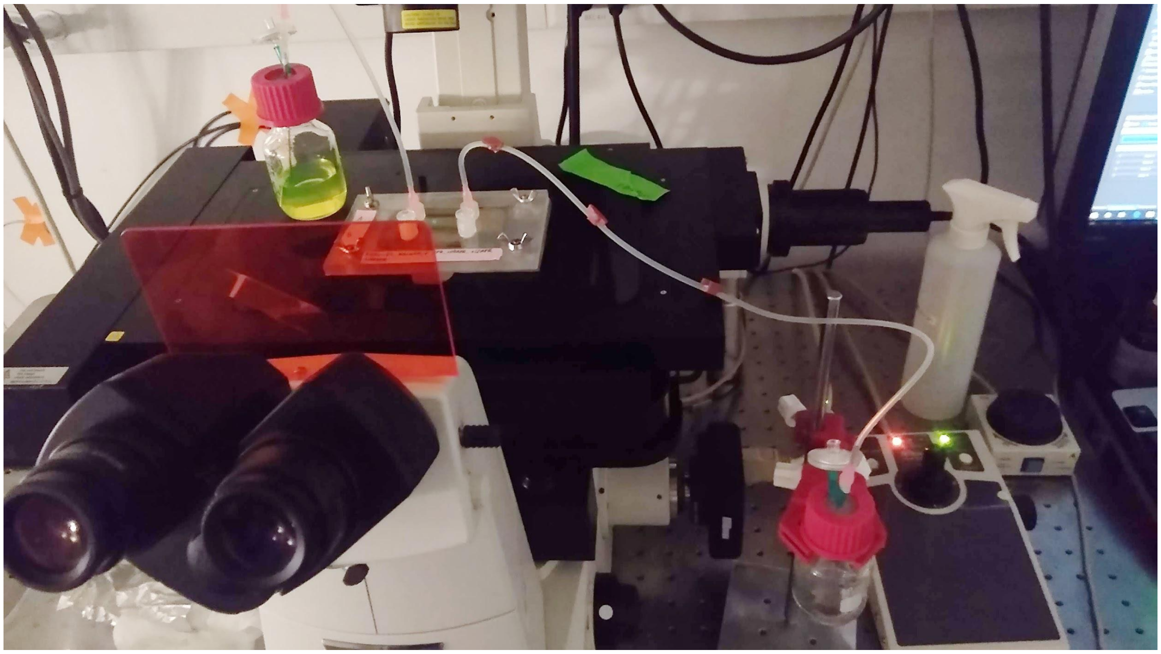

*Supp. Fig. 7.2: The FITC flow system uses the same components as the feeding system, but replaces the pump with gravity. A test tube stand allows for control of the height difference in order to adjust flow rate.*

##### 1.2: Prepare the FITC-Dextran solution

**FITC-Dextran (4 kDa) (Sigma, FD4-100MG) stock solution (10 mg/mL)**
Dissolve 100 mg of FITC-Dextran in 10 mL of PBS, aliquot, and freeze.

**FITC-Dextran (4kDa) working solution ( 0.5 mg/mL)**
Dilute 2 mL of 10 mg/mL FITC-Dextran into 38 mL of PBS to prepare 40 mL

#### 2. Calibrate the microscope settings

There are many settings which must be optimized to complete the assay. These include the choice of dye; concentration of dye; laser power and gain; image size; and frames per second in addition to the general workflow. Below are the settings we used:

a) Dye: FITC-Dextran
b) Concentration: 0.5 mg/mL
c) Laser power: 3, with a gain of 100
d) Image size: 512 pixels
e) Frames per second: 1

##### 2.1: Frame the point of interest in their X-Y positions

To get started, find the channel in the X-Y position. Depending on the microscope and its interface, this may require the user to know the coordinate placement, or it may be achieved by visually aligning the objective with the point of interest.

##### 2.2: Locate the ideal Z-position for capture

Effective imaging requires focusing confocal optics at the correct height. Focusing two low may fail to capture any fluorescent signal, while focusing too high may introduce optical distortions which compromise the collection of clear data. Locating the ideal Z-position presents two separate challenges that must both be addressed.

a) Identify what position in the sample you wish to target

It may seem intuitive to assume that the ideal z-position might be the midline of a channel, or a relatively broad area. We found that the z-plane needed to image the sample just slightly above the bottom of the channel in order to obtain the sharpest visible channel wall. It is recommended that users flush a channel with dye and then collect a Z-stack of at least five positions ranging from the lowest position at which the dye is visible to the highest in order to identify the best height relative to their channel. In our case, we found that the edge of the channel and cells appeared to produce shadows that grew worse as the target height got further from the floor of the channel.

b) Find the desired position

Once the ideal position in the channel is known, the user must locate the bottom of the channel. Identifying the top and bottom of channels seeded with a fluorescent marker-expressing cell line is fairly easy. Once the approximate height is known, the z-position can be found in future experiments by starting at the expected position and searching carefully while looking at a vacant channel with a high fluorescent power and a high gain.

#### 3. Capture diffusion

##### 3.1: Preparing the channel

The fluid contents of a channel must be evacuated in order to assess diffusion of a fluorescent dye. If the channel is filled with medium, the dye will be diluted and will not offer a reliable image demonstrating the channel boundary at image 0. Before aspirating the contents of the channel, however, all hardware and software settings (including framing) should be ready in order to begin capture immediately after flushing the channel.

a) Frame the channel in all dimensions.

b) Set the framerate, resolution, laser power, and gain. Make sure only the desired channel is active.

c) Set the time-series settings and the image stitching settings.

d) Prime the inflow line of the FITC flow system by connecting it to a syringe and gently filling the line. Clip it with a binder clip to prevent it from draining, and place the outflow bottle at the desired position.

e) Aspirate the contents of the channel using a pasteur pipet connected to a vacuum line. A low pressure is recommended. After extended culturing, the fibrin-gelatin matrix is likely softer and liable to tear or detach from the glass slide if handled roughly. It may be necessary to place the tip of the pasteur pipet at the mouth of the channel with the vacuum line disconnected or closed off before connecting or opening the line.

##### 3.2: Flush channel and begin capture.

a) Once the channel is empty, connect the primed, clipped, inlet flow line to one port of the housing and connect the outlet flow line to the other.

b) Seet the housing on the stage and reconfirm that the objective is positioned to capture the diffusion time series.

c) Before starting flow, tilt the housing to elevate the outflow side.

d) With the outflow reservoir more than six inches below the stage, attach a 10 mL syringe to the vent needle. Unclip the inflow line and gently pull to suction the FITC into the channel.

e) As soon as FITC enters the channel, reseat the housing in the stage bracket and begin capturing the time series.

f) Confirm that the FITC is dripping from the outflow needle in the outflow reservoir at a steady pace, then raise the outflow reservoir to slow the drips to the desired pace.

> The flow rate will depend on tube friction, which is determined by the tube length and diameter, as well as the needle length and diameter. Placing the inlet reservoir at the height of the stage and the outflow needle ~4” below the stage, however, provided a flow rate of ~ 0.3 mL/min, which used ~ 10 mL over a 30 min capture.

g) Let the time series run to completion.

> Repeat with additional channels as necessary.

#### 4. Post-process image stack

In order to quantify the data, the FITC assay images need to be re-registered to remove drift and normalized in intensity. They are then output as a series of separate tiff files and a gif.

a. Open FIJI and load the time series of interest

b. Convert the images from 16-bit to 8-bit in order to save as an animated gif (Image > Type > 8-Bit)

c. Register to correct for drift by selecting Plugins > StackReg

> Set Transformation to “Rigid Body”. *All slices will be registered towards the current slice when StackReg is run, so select the starting slice accordingly.*

d. Adjust brightness and contrast by selecting Image > Adjust > Brightness/Contrast and then, while viewing the darkest image, clicking “Auto” and “Apply”.

e. Save As an Image Sequence to generate a separate Tiff file for each slice (File > Save As > Image Sequence)

f. Save as an animated gif (File > Save As > Animated gif. “Set Delay in Milliseconds” to 50 and “Number of plays…” = 0)

#### 5. Analyze and quantify the rate of diffusion

Once a time series has been converted into individual images, these images must be loaded into R and converted into a matrix of pixel intensities. These intensities are then averaged across the length of the channel in order to create a vector of average fluorescent intensity perpendicular to the channel for each image.

The starting image is then used to identify a cutoff intensity value used to discern the signal of FITC diffusing outward from background noise. This cutoff is based on the outside edges of the profile. The pixels at the edge of the profile are assumed to contain no dye, so their maximum intensity benchmarks the background noise.

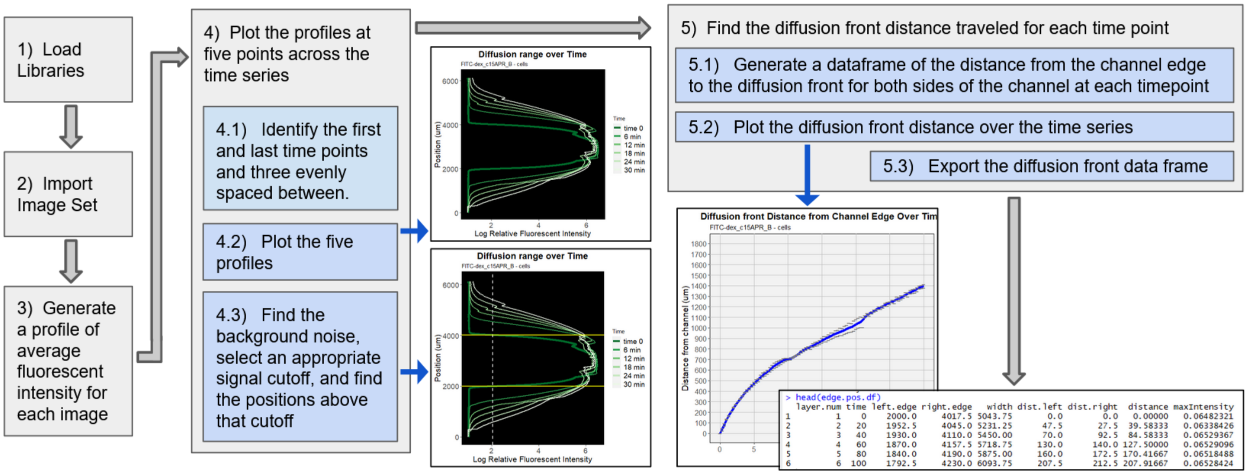

*Supp. Fig. 7.3: Code pipeline for converting image series into a table listing the distance between the diffusion front from the channel wall on each side of the channel for each timepoint.*

The cutoff is used to identify the front boundary of the FITC on each side of the channel. For the starting image, this boundary is assumed to be the edge of the channel. For each fluorescent profile, the boundary positions on each side are found and logged.

The distance between each boundary position in the time series from the starting boundary position is calculated, smoothed, and averaged between the two sides of the channel. This distance is then plotted to visualize the speed with which the FITC diffused out of the channel.

### 6. Compare the speed of diffusion between time serieses

Once the diffusion profiles and diffusion distance over time has been generated for a sample, it can be exported as a table in CSV format. These tables can be imported in a second script that allows for each to be plotted alongside any other.

**Applying a sanity check**

Before running an analysis, examine the gifs of various runs. While the quantification method provided is designed to make the analysis more objective, each time series should still be subjected to a sanity check. If an observable difference in diffusion is visible to the eye, the quantification methods can provide an average in the difference in slope. However if no discernable difference can be identified between conditions by eye, do not expect the quantification to be sensitive enough to provide evidence of a real diffusional barrier effect too small for observation by the human eye.
